# A Late Cretaceous Lonchodectid?

**DOI:** 10.1101/2019.12.17.879783

**Authors:** Carlos Albuquerque

## Abstract

A pterosaur ulnar specimen (NZMS CD 467) from the Mangahouanga Stream of New Zealand’s North Island has been first described by Wiffen et al 1988. Assumed to belong to a *“Santanadactylus-*like” pterosaur, this taxon has not since been extensively described, with only a few tentative claims that it represents an azhdarchid. Here, I re-examine the specimen and compare it to other pterodactyloid taxa, noting peculiar features such as its plug-like (obdurate) ulnar end. Christened *Parirau ataroa*, this taxon is found to be a lonchodectid, which alongside the North American *Navajodactylus boerei* extends this clade into the world’s youngest pterosaur faunas.

## Introduction

Through most of the 20th and early 21th centuries, the world’s last pterosaur faunas were assumed to be relictual, composed of only a few azhdarchids and nyctosaurids that were in the process of being outcompeted by birds. Several pterosaur remains dating to the Campanian and the Maastrichtian were in particular lumped with the former on the assumption that all other pterosaur clades were extinct (Fowler 2011, Witton 2013). However, recent studies have demonstrated that pterosaurs were not outcompeted by birds (Butler 2009, Witton 2013) and that their diversity in the younger stages of the Cretaceous was relatively high, including at least nyctosaurid, pteranodontid, thalassodromedid and tapejarid species in addition to a large azhdarchid diversity (Vremir 2013, Longrich 2018). Furthermore, the pterosaur fossil record is understood to suffer from severe taphonomic biases, lending to often long ghost-lineages (Lü 2017, Longrich 2018). This renders the status of several putative azhdarchids as at least worth re-examination.

One such taxon is NZMS CD 467, an ulnar specimen from the Mangahouanga Stream, part of New Zealand’s Tahora Formation. Found in Campanian estuarine deposits, this specimen was originally identified as an anhanguerid similar to “Santanadactylus” (now *Anhanguera) araripensis* albeit very tentatively so (Wiffen 1988). More recent sources have listed it as an azhdarchid remain (Witton 2013, Averianov 2014), however none of them have re-examined the fossil and all operate under the assumption that all Campanian and Maastrichtian pterosaur remains belong to azhdarchids or nyctosaurids. Therefore, a re-examination of the fossil is required, with an understanding that Late Cretaceous pterosaur ghost lineages are now plausible.

Understanding the phylogenetic positions of “problematic” taxa such as NZMS CD 467 will provide a clearer picture of the youngest pterosaur faunas.

## Materials and Methods

NZMS CD 467 was provided by Marianna Terezow from the National Paleontological Collection at GNS Science, Lower Hutt, while BEXHM 2015.18 was provided by Julian Porter from the Bexhill Museum and SMP VP-1853 was provided by Steven Jasinski from the State Museum of Pennsylvania. An Azhdarcho lancicollis picture (reversed for analogy) was provided by Alexander Averianov under free use stipulations:

https://commons.wikimedia.org/wiki/File:Azhdarcho.jpg

https://zookeys.pensoft.net/articles.php?id=4754

Phylogenetic studies and associated character matrix are discussed below.

## Results and Discussion

### Systematic Paleontology

Archosauria Cope, 1869

Pterosauria Kaup, 1834

Pterodactyloidea Plieninger, 1901

Coelobrachia Albuquerque, 2019

Lonchodectidae Hooley, 1914

### *Parirau ataroa* gen. et sp. nova

urn:lsid:zoobank.org:pub:73FCA0E4-D286-4174-8785-9E73CB83BD0F

### Etymology

“Parirau” is Māori for “wing” and “Ataroa” is a word that simultaneously means “night”, “darkness”, “moon” and “old”/“vestige”. Thus, “Ancient Wing”, mirroring Pterodactylus antiquus.

### Holotype

NZMS CD 467, left ulna.

### Horizon and Locality

Mangahouanga Stream, conglomeratic facies of the Maungataniwha Member of the Tahora Formation, North Island, New Zealand.

### Referred Material

NZMS CD 467 ulna.

### Diagnosis

Lonchodectid with asymmetrical distal ulna, bearing both dorsal and ventral depressions. The anterior section of the distal ulna is massively expanded, with dorsal and ventral elevations and a crest with both dorsal and ventral curvatures. The fovea carpalis is proportionally small, 6.2 mm in diameter. Ulna is pneumatic, with 1.4 mm bone walls on average. Possible 3.75 meter wingspan (see Lonchodectid Ecology section).

### Description

NZMS CD 467 is incomplete, with only the distal shaft and posterior part of the ulnar articulation remaining well-preserved. Nonetheless, the element does not appear to be flattened or bear extensive erosion facets aside from the missing elements. The missing section of the ulnar articulation is comparatively small, indicating that the posterior section of the ulnar joint was indeed reduced. This, alongside the straight shape of the ulna and its distal expansion and compression, supports a pterosaurian identity over an avian one. NZMS CD 467 measures 8.3 cm, the proportional wideness and thin (up to 1.4 mm) bone walls indicating a pneumatic state. This differs from the condition seen in azhdarchids, pteranodontids and nyctosaurids which have proportionally even thinner bone walls, in similar sized taxa rarely exceeding 0.31 mm and only approaching similar values at larger sizes (Bennett 1991, de Ricqlès 2000, Unwin 2003, Witton 2013) and more in line with lonchodectid ulnae which also show signs of pneumacy but have ulnar bone walls up to 2 mm thick (Rigal 2017).

**Figure 1.**
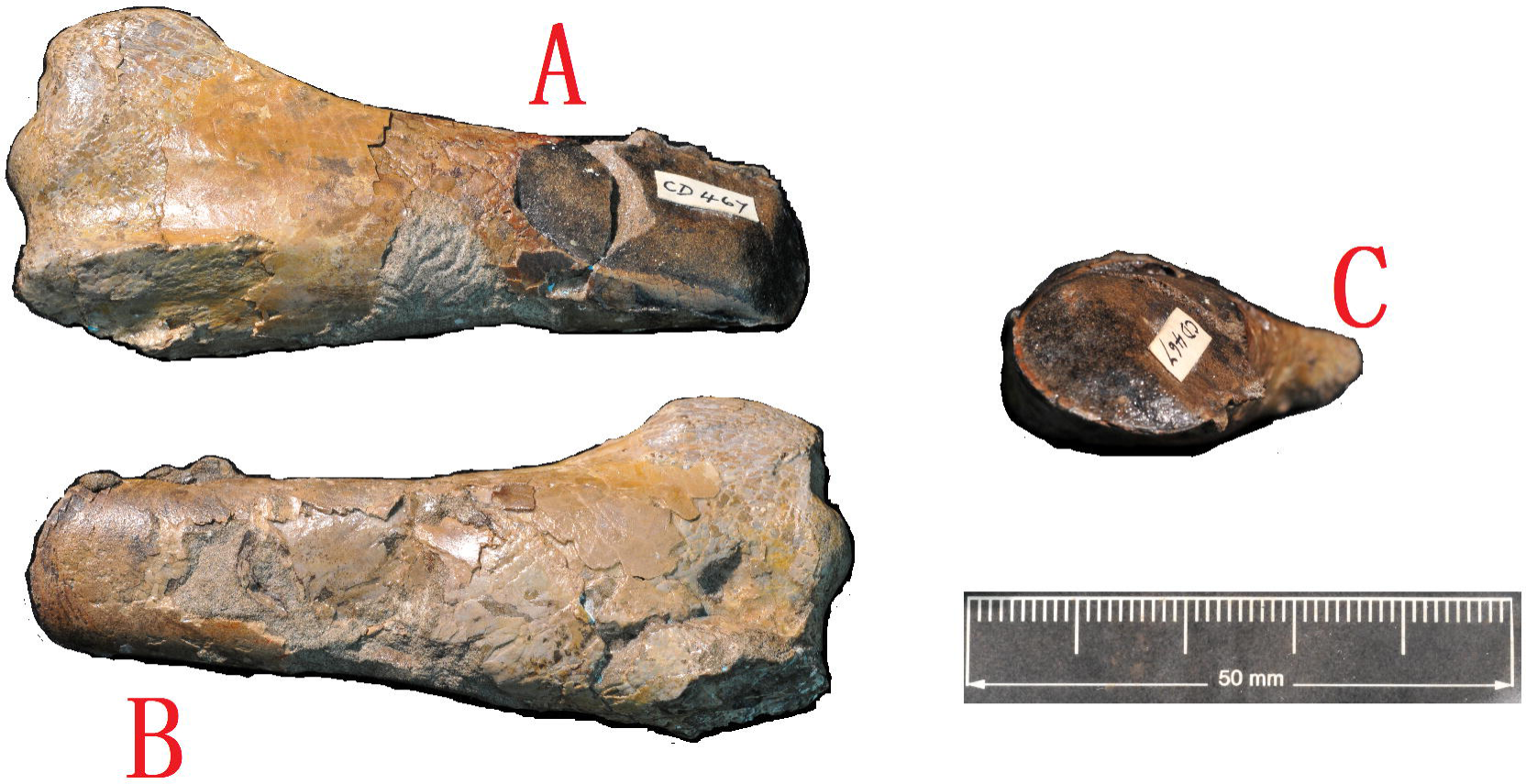
NZMS CD 467 shaft in dorsal (A), ventral (B) and mid-section (C) displays respectively.

Even more distinct from azhdarchid morphology is the anatomy of the ulnar articulation. The distal ulna is divided in anterior and posterior sections by both dorsal and ventral depressions which are immediately followed by dorsal and ventral ridges in the immediate area of the anterior ulna. These are in turn followed by a long, flattened crest with diagonal upwards and downwards curvatures, resulting in a triangular shape at cross-section. The *fovea carpalis* meanwhile occupies a fourth of the anterior ulnar joint surface, being oval in shape. In effect, the distal ulnar articulation strongly resembles a plug, a condition only seen in lonchodectid pterosaurs like *Serradraco* and *Prejanopterus* (Rigal 2017) as well as on *Navajodactylus boerei*, a taxon only tentatively identified as an azhdarchid and lacking diagnostic traits of the group (Fowler 2011, Witton 2013).

By contrast, the ulnae of azhdarchoid and ornithocheiroid pterosaurs have deep, short, highly expanded anterior distal joints, with a pronounced upwards curvature in derived ornithocheiroids. Istiodactylids meanwhile have a rectangular shape with a slight downwards curvature, while other pterosaurs have downwards-curving, hook-like anterior ulnae joints due to the lack of pneumatisation (see Wiffen 1988 Fig 2 amidst several others for comparisons). This therefore results in four broad types of pterosaur distal anterior ulnar joints: **bulbous** (azhdarchoids, derived ornithocheiroids), **obdurate** (lonchodectids), **rectangular** (istiodactylids) and **uncinated** (most pterosaurs).

**Figure 2.**
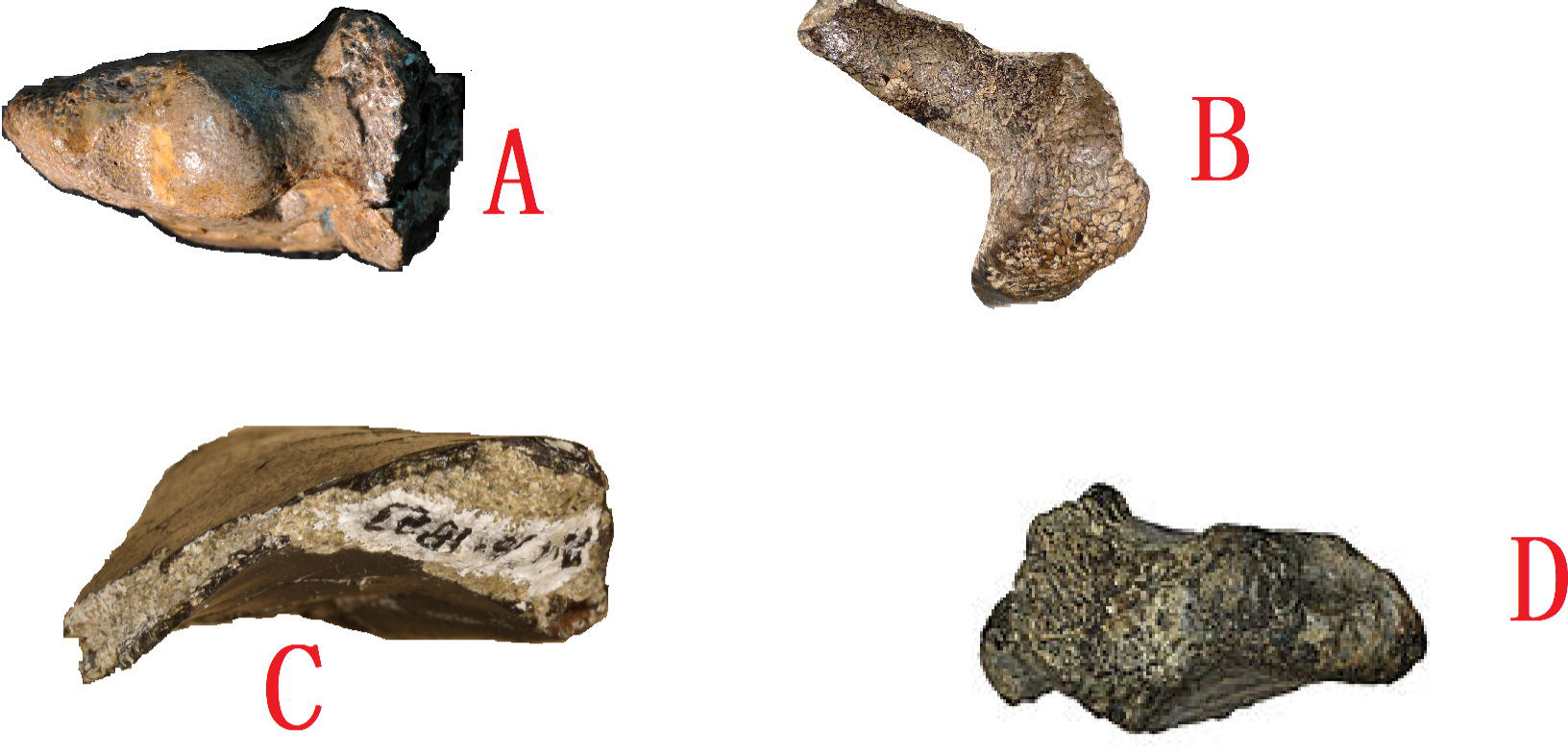
Distal ulnar ends of various pterosaur taxa: *Parirau ataroa* (A), *Serradraco sagittirostris* (B), *Navajodactylus boerei* (C) and *Azhdarcho lancicollis* (D). C and D (SMP VP- 1853 and ZIN PH 56/43 respectively) are right ulnas, so they were reversed in order to better compare to the other specimens, which are all left ulnas.

Both bulbous and uncinated morphologies are present in taxa covering a wide range of flight styles and ecological roles (Witton 2008, Witton 2010, Witton 2013, Longrich 2018 among others), while rectangular and obdurate morphologies are much more narrowly located, the former in an unambiguously monophyletic clade of scavenging soarers (Witton 2013) while the latter are in an ostensibly monophyletic clade with an unclear ecological niche (see Phylogenetic Analysis and Lonchodectid Ecology below). Therefore, it seems that anterior ulnar joint morphologies correlate much more strongly with phylogenetic signals than biomechanical ones. By contrast the size of the *fovea carpalis* and posterior ulnar joint seem to be more variable.

A tooth has been found in association with NZMS CD 467 (Wiffen 1988). Cut longitudinally, it cannot be currently diagnosed with confidence to any particular vertebrate group, as the extent of its enamel is difficult to ascertain. Bearing a somewhat tubular, recurved shape, it does superficially resemble unspecialised pterosaur teeth, but this morphology is also known in several crocodilian and actinopterygian groups, and conversely it does not resemble lonchodectid teeth, which are quasi-ziphodont. As such, it is not considered diagnostic of *Parirau ataroa* as to avoid rendering it a potential chimaera.

### Phylogenetic Analysis

**Fig 3.**
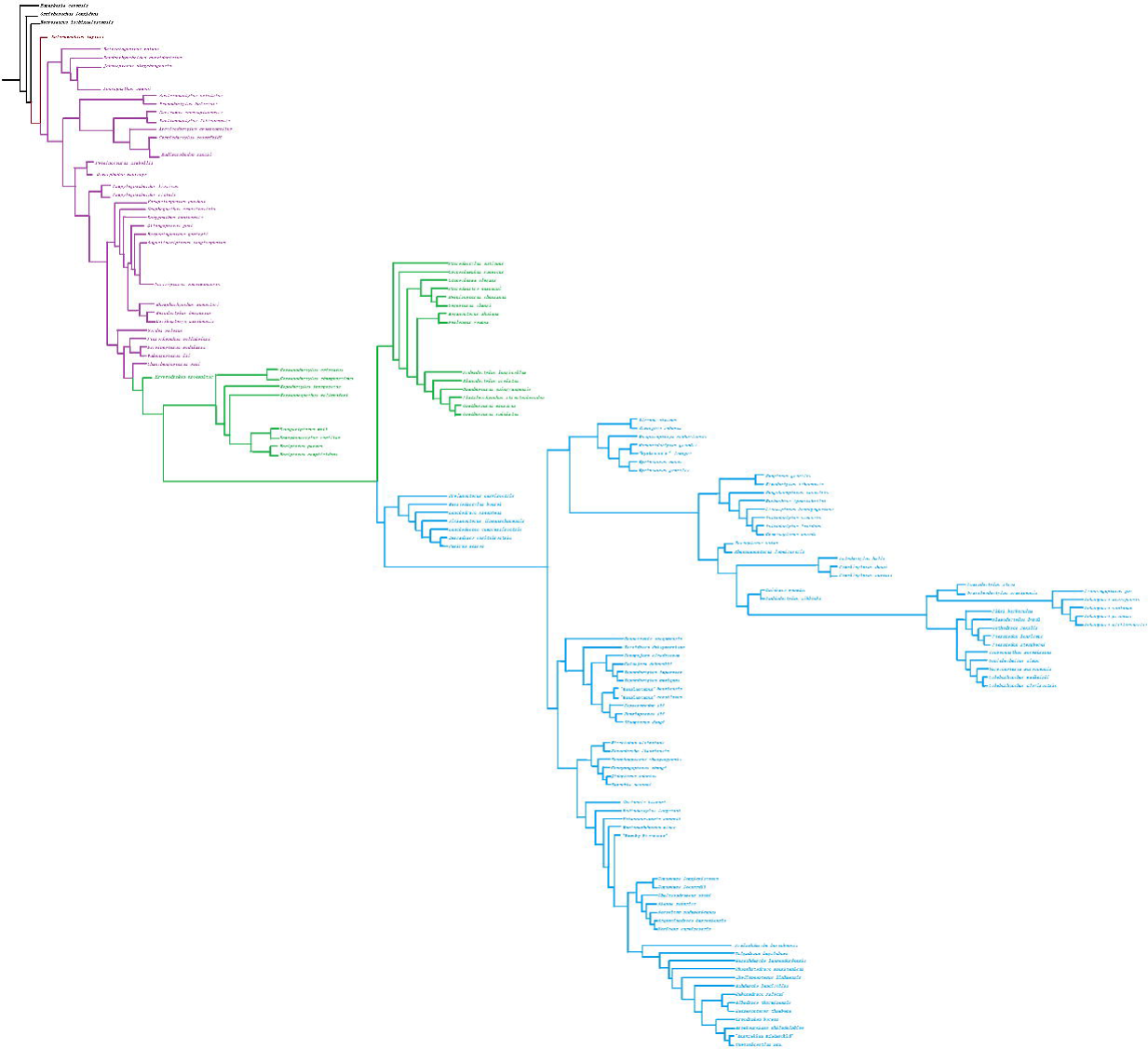
Strict consensus phylogenetic tree. Branch lengths are aesthetic rather than indicative of divergence rates or timing. Black = Archosauriformes, red = Pterosauromorpha, purple = Pterosauria, green = Pterodactyloidea, blue = Coelobrachia.

The dataset used in this study is based on the one presented by Longrich 2018, itself a derivative from the dataset utilised by Andres 2013 and ostensibly the most extensive character set derived from that one. It was done through the use of the software TNT 1.5 (Goloboff 2016) and modifications include added characters on the dataset (see Supplementary), the reasoning for their inclusion discussed below. As a whole, this analysis aims to emulate Hartman et al 2019’s methodology, examining the characters themselves and explaining why the results offer the currently most parsimonious interpretation of pterosaur phylogeny available. Naturally, the absence of genetic material means that crucial elements of pterosaur phylogeny will never be resolved, but the available morphology-driven phylogenetic analysis has wielded a surprising amount of consistency with pterosaur evolution and known fossil record, suggesting a relatively high degree of accuracy.

The results overall are consistent with previous studies in finding a monophyletic Pterosauromorpha, Pterosauria, Eopterosauria (though *Peteinosaurus zambellii* is now recovered as a dimorphodontid while *Parapsicephalus purdoni* becomes a basal rhamphorhynchid), Monofenestrata, Pterodactyloidea, Ctenochasmatoidea and Azhdarchoidea. Anurognathidae is however recovered as the most basal pterosaur clade as per Lü 2017 but *contra* several other Andres 2013 derived analyses, as supposed shared characters with pterodactyloids have developed differently. For instance, the dataset fails to acknowledge that anurognathid similarities with pterodactyloid pterosaurs such as their reduced tails clearly evolved convergently due to structural differences in the same characters (Witton 2013). This is an example of the organic character examination I seek to cultivate in this analysis.

### Coelobrachia

As with previous analyses, Azhdarchoidea and Ornithocheiroidea (*sensu* Unwin 2003; synonymous with “Pteranodontia”/”Pteranodontoidea” in other analyses but see below) are sister taxa. These two clades are united by a wide array of morphological characteristics such as proportionally small torsos, vestigial metacarpals l-lll that have lost contact with the wrist in many species, tall ridged cervicals (ridges secondarily reduced or lost in neoazhdarchians) and, most notably, the development of limb pneumacy, which in turn lead to the expansion of limb bones and ensuing similar shapes and proportions, the reduction of bone wall length. Pneumatic foramina are also known in *Pterodaustro guinazui* (Witton 2013), but this taxon lacks the extensive adaptations for pneumacy and multiple foramina seen in azhdarchoids and ornithocheiroids, suggesting that limb pneumacy might have happened as little as twice in pterosaur evolution. The development of forelimb pneumacy in the Azhdarchoidea + Ornithocheiroidea clade almost certainly happened earlier than in *Pterodaustro*, an Albian taxon which is nested among taxa with apneumatic limbs, and its development in the former group must have been a true innovation in pterosaur evolution, as it allowed lineages of this group to explode in diversity during the Early Cretaceous and quickly become the most speciose and ecologically diverse pterosaur clade known.

Because of the relevance of limb pneumacy in these pterosaurs, I propose the name Coelobrachia (Greek, “hollow arms”). This clade is defined as the group including the last common ancestor between *Azhdarcho lancicollis, Ornithocheirus simus* and *Lonchodectes compressirostris* and all its descendants. It is vaguely analogous to post-Andres 2013 definitions of Ornithocheiroidea, which are no longer applicable due to the removal of dsungaripterids from Azhdarchoidea and the complete polyphyly of Pteranodontia (see below). Therefore, Ornithocheiroidea returns to its Unwin 2003 definition.

Within Coelobrachia, Lonchodectidae is recovered as the most basal clade. This is *contra* previous post-Andres 2013 recoveries within Ornithocheiroidea in varying but usually basal positions. With the added characters and the inclusion of several other taxa suspected of being lonchodectids, an additional 11 steps are required to recover lonchodectids as ornithocheiroids. Due to the fragmentary nature of most remains the affinities between lonchodectid taxa remain tentative, with *Prejanopterus curvirostris* being rendered more basal due to its lack of phalange pneumacy and atypical jaw morphology.

Pteranodontia as originally defined in Unwin 2003 and later post-Andres 2013 studies is found to be completely polyphyletic: pteranodontids are found as part of Ornithocheirae (most parsimonious as a sister taxon to Ornithocheiridae, but positions within Ornithocheiridae, as sister to or within Anhangueria or sister to or within Targaryendraconia or basal to these all occur within two to three additional steps; see below for the status or Ornithocheirae as a whole) while nyctosaurids are found in a more basal position, most parsimoniously as within Ornithocheiroidea but taking an additional two steps as basal to all coelobrachians aside from lonchodectids. While both clades do strongly resemble each other in possessing toothless jaws and similar limb proportions, they differ radically in aspects of the pectoral girdle, pteranodontids grouping more closely with derived ornithocheiroids in possessing warped deltopectoral crests and similar scapulocoracoid morphology while nyctosaurids differ radically from most other coelobrachians in having hatchet-shaped deltopectoral crests and scapulocoracoids that do not articulate with the thoracic vertebrae/notarium. This is further consistent with the fossil record of both clades: putative nyctosaurids are known from the Berriasian (Dyke 2010) while unambiguous pteranodontids do not predate the Cenomanian, and the shared toothlessness might be associated with the general trends towards the prevalence of toothless taxa in Late Cretaceous pterosaur faunas (see On The Youngest Pterosaur Faunas).

*Piksi* is recovered as a basal pteranodontid in this analysis. This taxon is notoriously controversial as its status as either a pterosaur or a bird is ambiguous, being found as an avian theropod in Martin-Silverstone 2016 but as a pterosaur in Longrich 2018. As noted in the latter’s supplementary material a few decisive characters might only be available under special preparation, so it is beyond the scope of this paper to conclusively determine whereas *Piksi* is a pterosaur or a bird. However, its unique phylogenetic position in Longrich 2018 required a re¬evaluation of this taxon. *Contra* Longrich 2018’s results but following some observations in the supplementary material where similarities to pteranodontids, nyctosaurids and azhdarchids are noted, *Piksi* is found to be most parsimoniously a pteranodontid. Its basal position within Pteranodontidae might be deceptive however, as the holotype’s bone surface is rugose and could therefore indicate that the animal is a flapling.

Similarly, *Gwawinapterus beardi* was also included in this analyses. This taxon was originally interpreted as an istiodactylid pterosaur, but has since been re-identified as a saurodontid fish based on its dental replacement pattern (Vullo 2012, Witton 2013). However, more recently this type of dental replacement pattern has been identified in *Nurhachius* (Zhou 2019) and non-istiodactylid istiodactyliforms have been identified in the Cenomanian (Kellner 2019), so a re-examination of this specimen might be necessary. Here, it is recovered as an istiodactylid and the genus *Istiodactylus* is paraphyletic in relation to it.

Relationships within the rest of Ornithocheiroidea are found to be consistent with most recent studies, with Istiodactyliformes, Boreopteridae and Ornithocheirae being found as monophyletic and well defined. Ornithocheirae is found consistent with Pêgas 2019 in the presence of a Targaryendraconia, Ornithocheiridae and Anhangueria, but the definitions of these three clades are founded on relatively few characters and might flux in different analyses.

Azhdarchoidea’s topology is largely consistent with most studies since Unwin 2003, there being a distinction between Tapejaromorpha and Neoazhdarchia. Chaoyangopteridae and Thalassodromedidae are found to be within the latter with a considerably stronger support than for the former, requiring an additional 17 steps to place them in Tapejaromorpha. Within Neoazhdarchia, Thalassodromedidae is found to be closer to Azhdarchidae as opposed to several post-Andres 2013 studies demonstrating a “Neopterodactyloidea” composed of Azhdarchidae and Chaoyangopteridae; however such a clade can be produced within three additional steps. As with Longrich 2018, *Alanqa* and *Aerotitan* are found to be thalassodromedids as is *Argentinadraco* and *Xericeps* while *Cretornis, Radiodactylus, Palaeocursornis, Montanazhdarcho* and the “Hornby pterosaur” are found to be basal to the Thalassodromedidae + Azhdarchidae clade, but all of these taxa can be recovered as azhdarchids within one to three additional steps. *Samrukia* is recovered as a chaoyangopterid, but it is fragmentary enough to render such a placement over others as tentative for now.

A distinction between “slender” and “robust” azhdarchids has long been recognised (Witton 2013), and indeed at least a clade composed of *Cryodrakon + Arambourgiana + Quetzalcoatlus* and a clade composed of *Hatzegopteryx* + *Bakonydraco* + *Albadraco* are both strongly supported. However, the fragmentary nature of several specimens and taxa with “mosaic” characteristics such as *Volgadraco* and *Zhejiangopterus* render a deeper division within Azhdarchidae tentative at best, and it is possible that a clear answer to the overall topology of this clade may never come. *Contra* previous post-Andres 2013 studies, *Bakonydraco* is recovered as an azhdarchid as known post-cranial material hadn’t previously been considered.

### Dsungaripteroidea

Initially defined as all pterosaurs possessing a notarium (Kellner 2003), Dsungaripteroidea has since been redefined as a group of pterosaurs including all taxa derived from the last common ancestor between *Dsungaripterus* and *Germanodactylus* (Unwin 2003, Witton 2013). However, several post Andres 2013 studies have found this definition of Dsungaripteroidea to be polyphyletic, with germanodactylids being more closely related to Ctenochasmatoidea (thus forming the clade Archaeopterodactyloidea) while dsungaripterids are rendered neoazhdarchians usually closely related to thalassodromedids.

The problem with these results is that they are heavily reliant on the notarium as a character, when it has been noted that all pterosaurs with wingspans above two meters have notaria and none under that size value have it in spite of phylogenetic affiliation, while ignoring common characteristics between germanodactylids and dsungaripterids such as expanded opisthotic processes, highly recurved femora and non-pneumatic limb bones with thick bone walls (Witton 2013). With these features taken into consideration, dsungaripterids unsurprisingly nest with germanodactylids even when the notarium is listed as a character.

They do so most parsimoniously near the base of Pterodactyloidea, following perhaps previous observations that germanodactylids and wukongopterid skulls strongly resemble each other (Lü 2010), but it takes a single additional step to instead recover them as sister taxa to Ctenochasmatoidea, thus recovering a massively expanded Archaeopterodactyloidea. With this in mind, it might be worth investigating which characters are plesiomorphic to Pterodactyloidea as a whole or which ones can provide a monophyletic Archaeopterodactyloidea.

### On The Youngest Pterosaur Faunas

Table - A summary of known Campanian and Maastrichtian pterosaurs.

As noted previously, the pterosaur fossil record is strongly impacted by taphonomic biases. As such, a complete picture of the composition of the world’s last pterosaur faunas may always be out of reach, but some broad generalisations can be made:

- Even with the allocation of several taxa to other groups, Azhdarchidae remains the most speciose (or at least best represented) group. This seems to represent a genuine adaptive radiation as unambiguous Early Cretaceous azhdarchids are unknown and several Late Cretaceous fossil sites such as the Javelina Formation and the SebeşFormation bear multiple azhdarchid species at varying size ranges and morphologies (Vremir 2013). Whereas as solely represented by Azhdarchidae or with additional taxa such as thalassodromedids, Neoazhdarchia displays the broadest size range among Late Cretaceous pterosaurs, from raven sized species to the largest flying vertebrates of all time. Neoazhdarchians are understood to be terrestrially foraging carnivores and omnivores comparable to modern ground hornbills (Naish 2008, Witton 2013, Wu 2017), but the comparatively much higher species diversity, size range and morphology types hints at a greater nuance in terms of functional ecology.
- Nyctosauridae is the second most diverse clade. They and Pteranodontidae are the only known piscivorous pterosaurs and seem to have encroached into the ecological niches occupied by toothed ornithocheiroids in the pre-Cenomanian Cretaceous.
- Toothed pterosaurs appear to be rare at best or absent altogether, the only known examples being lonchodectids (which in the absence of cranial materials doesn’t preclude the possibility of tooth loss in *Navajodactylus* and *Parirau)* though istiodactylids and anurognathids may have also been present (see Table). Because toothless and toothed taxa co-exist in most Early Cretaceous locales and the extinction of at least some toothed pterosaurs occurred abruptly in the Cenomanian/Turonian boundary (Pentland 2019), it seems more likely that toothless species simply moved to occupy niches left vacated by the extinction of toothed pterosaurs. Toothless beaks are thought to correlate with more generalistic diets and more efficient manipulation of objects in the mouth among dinosaurs (Field 2019), while the teeth of toothed pterosaurs are noted as being very specialised (Witton 2013), suggesting that toothless pterosaurs likely had broader diets and thus were better equipped to deal with the ecological turnovers of the Cenomanian and Turonian. A similar situation happened with contemporary mammalian faunas, with the hypercarnivorous eutriconodonts and symmetrodonts declining while more omnivorous therians, dryolestoids and multituberculates thrived (Grossnickle 2013). The toothed pterosaurs that survived were thus either similarly generalistic or were specialised to relatively stable niches.

Longrich 2018 proposed that Late Cretaceous pterosaurs were displaced at small sizes by birds and that in turn these were unable to grow to large sizes due to competition from pterosaurs (the noted threshold for both being wingspans of 2 meters), mirroring the niche partitioning between flightless dinosaurs and mammals. However, small pterosaurs are known (“Hornby pterosaur”, *Piksi)* and their existence is implied regardless due to the superprecocial lifecycle of pterosaurs in general (Witton 2013), meaning that birds would have regularly faced competition from adult and flapling pterosaurs at small sizes. Similarly, Late Cretaceous birds are noted to be constrained at small sizes (Longrich 2011), implying that miniscule pterosaurs might have also been present. Therefore, pterosaurs were clearly not restricted in terms of body size and the niche partitioning between both groups might have been more similar to that of modern birds and bats, in which both clades are highly diverse but one is restricted in terms of body size (Mesozoic birds, bats) while the other is considerably more diverse (pterosaurs, Cenozoic birds).

This, combined with the general prevalence of highly productive ecosystems such as forests and wetlands in the warmer Late Cretaceous climate, suggests that the youngest pterosaur faunas, much like modern bat and bird faunas, might very well have had hundreds if not thousands of species. Even relatively unproductive environments such as arid plains (i.e. Javelina Formation) and the open sea (i.e. Ouled Abdoun Basin) produce highly diverse pterosaur faunas, further hinting at the sheer diversity of less extreme ecosystems.

Thus, rather than declining, the last pterosaurs were thriving, and it took the KT mass extinction event to wipe them out alongside nearly all birds aside from a few ornithurines. After the extinction of pterosaurs, flying birds quickly radiated into both large and small sizes (Longrich 2011, Longrich 2018), further attesting to the ecological constraints Mesozoic birds faced.

## Lonchodectid Ecology

The ecology of lonchodectid pterosaurs has always been a controversial subject. Unwin 2006 suggested a piscivorous lifestyle while Witton 2013 suggested terrestrial azhdarchid-like habits based on the proportions of an undescribed specimen, the “Moon Goddess”. Complicating matters is the reassignment of some of *Lonchodectes’* post-cranial remains to *Ornithostoma* (Averianov 2012), and the post-cranial remains of *Yixianopterus, Lonchodraco, Prejanopterus* and *Serradraco* not being discussed in terms of proportions or relations to the “Moon Goddess” specimen. The “Moon Goddess” specimen itself may be a composite, and its manuscript will not be published (Witton 2016).

The lonchodectid femur is known in *Yixianopterus* and *Prejanopterus.* It is somewhat comparable in size and thickness to that of ctenochasmatoids (Pereda-Suberbiola 2012), suggesting that these pterosaurs could in fact walk competently on the ground. Lonchodectid wing material is more broadly known and far more complete; unlike azhdarchids and ctenochasmatoids, lonchodectids had proportionally massive wings comparable in proportions to those of ornithocheiroids. This suggests that lonchodectids spend most of their time on the wing or were rather acrobatic flyers; whereas their unique obdurate ulnar morphology might have played a role in their flight style is worth investigating.

Most lonchodectid remains occur in inland freshwater sites (exceptions like *Parirau* and *Lonchodectes* are usually fragmentary, suggesting long post-mortem aquatic transportation); combined with their general rarity compared to other pterosaur remains, this suggests that lonchodectids weren’t aquatic and only occasionally would have foraged at the water’s edge. Their teeth are similarly highly distinct from those of piscivorous pterosaurs; rather than conical, they are quasi-ziphodont and spaced and raised by elevated tooth sockets. While adequate to catch fish, they were more suited to rip flesh. Conversely, lonchodectids lack the adaptations for scavenging seen in istiodactylids (Witton 2013), so while they would have likely scavenged on occasion they were more specialised to hunt living prey.

The overall picture seems to be that of pterosaurs analogous to modern kites and falcons, hunting vertebrates on land or on the wing. Their long wings would have suited both dynamic soaring flight as well as continuous flapping, allowing lonchodectids to both travel long distances in search for prey as well as to pursue it on the wing. Prey might have been killed and consumed on the ground as in modern birds of prey and bats, but modern frigatebirds are known to snatch turtles and other small prey items from the ground on flight, so lonchodectids might similarly have been a rare case of non-avian aerial predators snatching prey from the ground on the wing. They likely could have similarly caught fish on the water surface this same way, but their terrestrial lifestyle preferences, speciations to process meat and competition from ornithocheiroid pterosaurs might have rendered this a rare occurrence.

Because of the specialised nature of lonchodectid dentition, it seems unlikely that lonchodectids represent the ancestral condition for Coelobrachia. However, their jaws do resemble those of both ctenochasmatoids and several ornithocheiroids like *Cearadactylus* and they display both competent terrestrial locomotion like azhdarchoids and spéciations to a highly aerial lifestyle like ornithocheiroids, suggesting that the ancestral coelobrachian likely led a mosaic lifestyle.

Wiffen 1988 proposed a 3.75 meter wingspan for NZMS CD 467, based on comparisons to anhanguerids. This size value likely remains the same for *Parirau ataroa*, and indeed it is similar to the one proposed for *Navajodactylus boerei* (Fowler 2011), though the latter taxon would have been larger. Regardless they would be closest in size to *Unwindia* among lonchodectids. They are comparable to the largest living flying birds, but mid-sized by Late Cretaceous pterosaur standards. *Navajodactylus boerei* appears to have been reasonably successful, its remains not only found in opposite parts of Laramidia but also common for pterosaur remains of the time.

It is possible that some Campanian and Maastrichtian *Pteraichnus* tracks were made by lonchodectids. These tracks are different from those associated with azhdarchid pterosaurs, *Haenamichnus* (Pérez-Lorente 2014).

## Conclusion

With the re-description of NZMS CD 467, Lonchodectidae is extended into the Late Cretaceous and New Zealand acquires its first named pterosaur taxon. *Parirau ataroa* further demonstrates that the world’s last pterosaur faunas were richer than previously imagined and that taxa should not be classified solely on the basis of temporal distribution. Additionally a clearer insight towards pterosaur phylogenetics was hopefully achieved.

