## Supplementary for "A Late Cretaceous Lonchodectid?"

**Supplementary Material**

Appendix: Taxon-character matrix for the phylogenetic analysis of Pterosauria.

Euparkeria_capensis 0 - 1 0 0 - 0 0 0 0 0 0 0 1 0 0 0 0 0 0 0 - 0 0 0 0 0 0 0 0 0 - - - - - - - 0 0 0 0 0 0 0 0 1 0 0 - - 0 0 0 0 - 0 0 0 0 0 0 0 - 0 0 0 0 0 0 0 0 0 0 0 0 0 0 0 0 - - 0 0 0 0 0 0 1 0 0 0 0 - 0 0 0 0 0 0 0 0 0 0 0 0 - 0 0 1 0 0 - - 0 0 1 0 - 0 1 0 0 0 0 0 0 0 0 0 1 0 0 0 1 1 0 0 0 0 0 0 0 0 0 0 0 0 0 0 0 0 0 0 0 0 0 0 0 0 0 0 0 0 0 0 0 0 0 0 0 0 0 0 0 0 0 0 0 0 0 0 0 0 0 - 0 - - - - 0 0 0 0 0 ? ? ? 0 0 0 - - 0 0 0 0 0 0 0 0 0 0 0 0 0 0 1 0 0 0 0 0 0 1 0 0 1

Ornithosuchus_longidens 0 - 1 ? 0 - 0 0 0 0 0 0 0 1 ? 0 0 0 0 0 0 ? 2 0 0 1 0 0 1 ? 0 - - - - - - - 0 0 0 0 0 0 0 0 0 ? 0 - - 1 0 1 0 - 0 0 0 0 0 0 0 - 0 0 0 0 0 0 0 0 0 ? ? ? 0 ? ? ? ? ? 0 ? 0 ? 0 0 1 0 0 0 0 - 0 0 0 0 0 0 ? ? ? ? ? ? ? 0 0 1 0 0 - - 0 0 1 0 - 0 0 0 0 0 0 0 0 0 0 0 1 0 0 0 1 1 0 0 0 ? 0 0 0 0 0 0 0 0 0 [01] 0 0 ? 0 0 0 0 0 0 0 0 0 0 0 1 0 1 0 0 0 0 0 0 0 0 0 0 0 1 0 0 0 ? 0 0 - ? - - - - 0 0 0 ? ? 0 ? ? 0 0 0 - - 0 0 0 0 0 0 0 0 0 0 0 0 1 ? 0 ? 0 1 0 0 0 0 0 0 1 0 0 1

Herrerasaurus_ischigualastensis 0 - 1 ? 0 - 0 0 0 0 0 0 0 1 0 0 0 0 1 0 0 - 0 0 0 0 0 0 0 0 0 - - - - - - - 0 0 0 0 2 0 0 0 0 0 0 - - 1 0 0 0 - 0 0 0 0 0 0 0 - 0 0 0 0 0 0 0 0 0 0 0 0 0 0 ? ? ? ? 0 ? 0 ? 0 0 1 0 0 0 0 - 0 0 0 0 0 0 0 0 ? ? ? ? ? 0 0 2 0 0 - - 0 0 1 0 - 0 0 0 0 0 0 0 0 0 0 0 1 0 0 0 1 1 0 0 0 0 0 0 0 0 0 0 0 ? 0 0 0 0 1 0 0 0 0 0 0 0 0 ? ? ? ? ? ? ? ? ? 0 0 0 0 1 0 0 0 1 0 1 0 0 0 0 - 0 - - - - 0 0 1 1 0 0 0 0 0 0 0 - - 0 0 0 0 0 0 1 0 0 0 0 0 0 0 3 0 0 1 0 0 0 0 0 0 1 0 0 1

Scleromochlus_taylori 1 - ? 0 0 - 0 0 0 0 1 0 ? 1 ? ? 0 1 0 0 0 ? 0 0 ? 0 ? 0 1 0 0 - - - - - - - 0 1 0 0 0 0 0 0 0 ? 0 - - 1 ? ? 0 - 0 ? 0 2 0 ? ? - 1 0 ? ? ? 0 0 1 ? 0 0 0 0 ? ? ? ? ? 0 0 ? ? 0 ? 1 0 0 0 0 - 0 0 ? 0 0 0 0 0 ? ? ? ? ? 0 0 1 2 0 - - 0 0 1 0 ? ? ? 0 0 0 0 ? ? ? ? ? 1 ? ? ? 1 ? ? ? 0 0 ? ? ? 1 0 0 0 0 ? ? 0 0 0 0 0 0 0 0 0 0 0 0 0 ? ? ? ? ? ? ? ? 0 ? 0 1 ? ? ? 1 0 0 0 ? ? ? ? ? - - - - 0 1 ? ? ? ? ? ? 0 0 0 - - 0 0 0 0 0 0 0 1 ? 0 0 ? 0 0 4 - 0 1 0 0 0 0 0 0 1 0 0 1

Batrachognathus_volans 1 - 1 - 0 - 0 0 0 1 ? 0 3 0 - 0 ? - - 1 0 0 0 0 0 1 1 1 - 1 0 - - - - - - - 0 - - - - 0 - - - ? 0 - - 1 ? 0 0 - 0 0 ? 0 0 1 ? 0 ? 0 0 1 1 0 ? ? ? ? ? ? 0 ? - - - - 1 0 ? 1 0 1 1 0 1 0 ? - 0 0 2 0 0 0 1 - 0 0 0 0 - 1 ? 2 2 0 - - 1 0 1 0 - 3 0 0 0 2 2 0 1 0 0 0 1 1 - - 1 0 0 0 0 ? ? ? ? ? ? 0 0 1 ? ? 0 0 ? ? ? ? ? ? 0 0 0 0 2 0 1 0 0 0 0 4 0 0 0 1 1 0 ? ? 1 0 0 0 0 0 0 ? ? ? ? ? ? 0 1 0 1 0 1 ? ? ? ? ? ? 1 ? 0 1 0 0 ? 0 1 ? 1 ? ? 0 0 ? ? 0 1 0 0 0 0 0 0 1 0 0 1

Jeholopterus_ningchengensis 1 - 1 - 0 - 0 0 0 1 ? 0 3 0 - 0 ? - - 1 0 0 0 0 0 1 1 1 - 1 0 - - - - - - - 0 - - - - 0 - - - ? 0 - - 1 0 0 0 - 0 0 ? 0 0 1 ? 0 ? 0 0 1 ? 0 ? ? ? ? ? ? 0 ? - - - - 1 0 ? 1 ? 1 1 0 1 0 0 - ? 0 2 0 0 0 ? ? ? ? 0 0 - 1 ? 2 2 ? ? ? ? 0 1 0 - 3 0 0 0 2 2 0 1 0 0 0 1 1 - - 1 0 0 0 0 ? ? ? ? 1 0 ? ? 1 1 0 0 ? ? 0 1 0 0 0 0 0 0 0 2 ? ? ? ? 0 0 ? ? 0 0 1 1 1 0 ? 1 0 4 0 0 0 0 0 ? ? ? 5 0 0 1 0 1 0 1 0 0 ? ? 1 1 1 0 0 1 0 0 ? 0 1 ? 1 1 ? 0 0 2 0 0 1 0 0 0 0 0 0 1 0 0 1

Anurognathus_ammoni 1 ? 1 - 0 - 0 0 0 1 1 0 3 0 - 0 ? - - 1 0 0 0 0 0 1 ? 1 - 1 0 - - - - - - - 0 - - - - 0 - - - 0 0 - - 1 0 0 0 - 0 0 ? 0 0 1 ? 0 ? 0 0 1 1 0 ? ? ? ? ? ? 0 0 - - - - 1 0 ? 1 0 1 1 0 1 0 0 - 0 0 2 0 0 0 1 - 0 0 0 0 - 1 ? 2 2 0 - - 1 0 1 0 - 3 0 0 0 2 2 0 1 0 0 0 1 1 - - 1 0 0 0 0 1 ? ? ? ? ? ? ? 1 ? 0 0 0 1 0 1 0 0 0 0 0 0 0 2 ? ? ? ? ? ? ? ? 0 0 1 1 1 ? ? 1 0 4 0 0 0 ? ? 0 ? ? 5 0 0 1 0 1 0 1 ? ? 1 0 1 ? ? 0 0 1 0 0 ? 0 1 0 1 1 0 0 0 2 0 0 1 0 0 0 0 0 0 1 0 0 1

Dendrorhynchoides_curvidentatus 1 - 1 - 0 - 0 0 0 1 ? 0 3 0 - 0 ? ? ? ? ? 0 ? ? ? ? ? 1 ? ? 0 - - - - - - - 0 ? ? ? ? ? ? ? ? ? ? ? ? ? ? ? ? ? ? ? ? ? 0 1 ? ? ? ? ? ? ? ? ? ? ? 0 0 1 ? ? - - - - 1 0 ? 1 0 ? 1 0 1 0 0 - 0 0 2 0 0 0 ? ? 0 ? 0 0 - 1 ? ? ? 0 - - ? 0 1 0 - 3 0 0 0 2 2 ? 1 0 0 0 1 1 - - ? ? ? 0 0 ? ? ? ? 1 0 ? ? 1 ? ? 0 ? ? ? 1 0 0 0 0 0 0 0 2 ? ? ? ? 0 0 4 ? 0 0 1 1 1 0 ? 1 0 0 0 0 0 ? ? 0 ? ? 5 0 0 1 0 1 ? 1 ? 0 1 ? ? ? ? 0 0 ? 0 0 ? ? 1 ? ? ? 0 ? 0 2 1 0 1 0 0 0 0 0 0 1 0 0 1

Eudimorphodon_ranzii 1 - 1 0 0 - 0 0 0 0 1 0 ? 1 0 1 0 1 0 0 0 - 0 0 1 1 1 0 2 0 0 - - - - - - - 0 1 0 0 3 0 0 0 0 0 0 - - 1 0 0 0 - 0 0 0 2 1 0 0 - 1 0 0 0 0 0 0 1 0 ? ? ? 0 ? 0 0 - - 0 1 ? ? 0 0 2 0 0 0 1 0 0 0 0 0 0 0 0 1 0 0 0 0 - 0 0 1 1 0 - - 0 0 0 1 0 1 1 0 0 2 0 1 1 0 0 1 1 0 4 2 1 1 0 1 0 ? 1 0 ? ? ? 0 0 0 ? 0 0 ? 1 ? ? ? 0 ? 0 0 0 0 2 0 1 0 1 1 0 1 0 0 0 1 1 0 0 0 1 0 1 0 0 0 ? ? 0 ? ? 1 0 0 0 0 1 0 ? ? ? ? ? ? 0 0 ? ? 0 0 0 ? 0 1 ? 0 ? 0 0 ? ? ? 0 1 0 0 0 0 0 0 1 0 0 1

Carniadactylus_rosenfeldi 1 - 1 ? 0 - 0 0 0 0 1 ? ? 1 0 1 0 1 0 0 0 - 0 0 1 1 1 0 2 0 0 - - - - - - - 0 1 0 0 3 0 0 0 0 0 0 - - 1 ? 0 0 - 0 0 0 2 1 0 0 - 1 0 0 0 0 ? 0 1 0 0 0 1 0 0 ? ? ? ? ? ? ? ? 0 0 1 0 0 0 1 0 0 0 0 0 0 0 0 1 ? ? ? ? ? 0 0 1 1 0 - - 0 0 0 1 0 1 0 0 0 2 0 1 1 0 0 1 1 0 4 2 1 1 0 1 0 ? 1 0 ? 0 0 0 0 0 0 ? 0 ? ? ? 0 0 0 0 ? 0 0 0 1 0 1 0 1 0 0 1 ? ? 0 1 1 0 0 ? 1 0 1 0 0 0 0 0 0 ? 0 0 0 0 0 0 1 0 0 0 0 0 0 0 ? ? 0 0 ? 0 0 ? 0 1 ? ? ? 0 ? 0 2 0 0 1 0 0 0 0 0 0 1 0 0 1

Arcticodactylus_cromptonellus ? ? ? ? ? - ? 0 0 ? ? 0 0 ? 0 ? ? 1 0 0 ? - ? ? ? ? ? ? ? ? ? ? ? ? ? ? ? ? 0 1 0 ? 2 0 0 ? 0 0 ? ? ? ? ? 0 ? ? 0 ? ? ? ? 0 ? - ? 0 ? ? ? 0 ? ? ? ? ? ? ? ? ? ? ? ? ? ? ? ? ? 0 1 0 0 0 ? ? 0 0 0 ? 0 0 ? ? ? ? ? ? ? ? ? ? ? 0 - - ? 0 0 1 0 1 0 0 0 ? 0 1 1 0 0 [01] 1 0 4 2 ? 1 0 ? ? ? ? 0 ? ? ? 0 ? ? ? ? ? 0 ? ? ? 0 0 ? 0 0 ? 0 1 0 1 ? 0 0 0 1 ? 0 ? 1 1 ? 0 0 1 0 1 0 ? ? ? ? 0 ? ? ? ? 0 ? 0 1 0 ? ? 0 ? ? ? ? ? ? ? ? ? ? ? ? 1 0 0 0 0 0 ? 2 0 0 1 0 0 0 0 0 0 1 0 0 1

Caviramus_schesaplanensis ? ? ? ? 0 - ? ? ? ? ? ? ? ? ? ? ? 1 ? ? ? ? ? ? ? ? ? ? ? ? ? ? ? ? ? ? ? ? ? ? ? ? ? ? ? ? ? ? ? ? ? ? ? ? ? ? ? ? ? ? ? ? ? ? ? ? ? ? ? ? ? ? ? ? ? ? ? ? ? ? ? ? ? ? ? ? ? 0 1 0 0 0 1 1 1 1 1 0 0 0 0 0 ? ? ? ? ? 0 0 0 2 0 - - 0 0 ? ? ? 1 0 0 0 1 0 ? ? 0 ? [01] 1 0 3 2 ? ? 0 1 0 ? ? ? ? ? ? ? ? ? ? ? ? ? ? ? ? ? ? ? ? ? ? ? ? ? ? ? ? ? ? ? ? ? ? ? ? ? ? ? ? ? ? ? ? ? ? ? ? ? ? ? ? ? ? ? ? ? ? ? ? ? ? ? ? ? ? ? ? ? ? ? ? ? ? ? ? ? ? ? ? ? ? ? ? ? ? ? ? ? 1 ? ? ?

Raeticodactylus_filisurensis 1 - 1 ? 0 - 0 0 0 0 1 ? ? 1 0 1 0 1 1 0 0 - 0 0 ? 0 1 0 2 0 1 0 1 0 1 0 0 0 0 1 0 0 2 0 0 0 0 0 0 - - ? 0 0 0 - 0 0 0 2 0 0 0 - 1 0 0 0 0 0 0 1 0 ? ? ? 0 ? ? ? ? ? 0 1 ? ? ? 0 1 2 0 0 1 1 1 1 1 0 0 0 0 0 ? ? ? ? ? 0 0 0 2 1 - - 0 0 0 1 2 1 1 0 0 1 0 1 0 0 0 1 1 0 3 2 1 1 0 1 0 ? 1 0 ? 0 0 0 0 ? ? ? 0 0 ? ? ? ? ? ? ? ? 0 ? ? ? ? ? ? ? ? ? 0 0 ? 1 1 ? ? ? 1 0 1 0 0 0 0 ? ? ? ? 0 ? 0 0 0 ? ? 0 ? 0 ? ? ? ? ? ? ? ? ? ? ? ? 1 0 0 0 ? 0 ? ? 0 0 1 0 0 0 0 0 0 1 0 0 1

Austriadactylus_cristatus 1 - 1 ? 0 - ? 0 0 0 1 0 ? 1 0 1 ? 1 0 0 0 - 0 0 ? 0 0 0 1 ? 1 0 1 0 2 0 0 0 0 1 0 0 1 0 0 1 0 0 0 - - 1 0 0 0 - 0 0 ? 1 0 0 0 - ? 0 0 0 0 0 ? ? 0 ? ? 1 0 ? ? ? ? ? 0 ? ? ? ? ? 2 0 0 0 0 - 0 0 0 ? 0 0 0 0 ? ? ? ? ? 0 0 1 2 0 - - 0 0 0 1 [01] 1 1 0 0 2 0 1 0 0 0 0 1 0 1 1 1 1 0 0 0 ? ? ? ? 0 0 0 0 0 0 0 0 ? ? ? 0 0 0 0 0 0 ? 0 1 ? ? ? ? 0 0 1 ? ? ? 1 1 ? ? ? ? 0 1 0 0 0 ? ? ? ? ? ? 0 0 0 0 1 ? ? ? 0 ? 0 0 ? ? ? ? 0 0 0 ? 0 1 ? ? ? ? ? ? ? ? 0 1 0 0 0 0 0 0 1 0 0 1

Preondactylus_bufarinii 1 - 1 ? 0 - 0 0 0 0 1 ? ? 1 0 1 0 1 0 0 0 - ? 0 ? ? ? 0 1 0 0 - - - - - - - 0 1 0 0 1 0 0 1 0 0 0 - - ? ? 0 ? ? ? ? ? 1 0 0 0 - 1 0 0 0 0 ? ? 1 ? ? ? ? ? ? ? ? ? ? ? ? ? ? ? 0 1 0 0 0 0 - 0 0 0 0 0 0 0 0 ? ? ? ? ? 0 0 1 1 0 - - ? 0 0 1 0 1 ? 0 0 2 0 1 0 0 0 0 1 0 1 1 1 1 0 0 0 0 ? ? ? ? 0 ? 0 0 ? ? ? 0 ? ? ? 0 0 0 ? 0 ? 0 1 ? ? ? ? ? ? ? ? ? ? 1 1 ? ? ? ? 0 1 0 0 0 ? ? ? ? ? 0 0 0 0 0 1 0 0 ? 0 0 0 0 0 0 0 0 0 0 0 0 0 1 ? 0 0 ? 0 0 2 0 ? ? ? ? ? ? ? ? 1 0 0 1

Dimorphodon_macronyx 1 - 1 ? 0 - 0 0 0 0 1 ? ? 0 0 1 1 1 0 0 0 - 2 0 3 0 0 0 2 1 0 - - - - - - - 0 1 1 1 1 0 0 1 0 0 0 - - 1 1 0 ? ? 0 0 0 1 0 0 0 - 1 0 0 0 0 0 0 1 ? ? ? 1 0 ? ? ? ? ? 0 1 ? ? ? 0 1 2 0 0 0 - 0 0 0 0 0 0 0 0 0 0 0 0 - 2 0 1 3 2 0 0 1 0 0 1 - 4 1 0 0 2 1 1 0 0 0 0 1 1 - - 1 1 0 0 0 0 1 1 ? ? ? 0 0 0 ? ? 0 0 1 0 0 2 0 ? 0 0 0 1 2 ? ? ? ? 0 0 ? 0 0 0 1 1 1 0 0 1 0 2 0 0 0 ? ? 0 0 0 2 0 0 0 0 1 0 0 0 0 1 0 1 1 3 0 0 1 0 0 1 0 1 0 0 0 0 0 0 2 0 0 1 0 0 0 0 0 0 1 0 0 1

Peteinosaurus_zambellii ? ? ? ? ? - ? ? ? ? ? ? ? ? ? ? ? ? ? ? ? ? ? ? ? ? ? ? ? ? ? ? ? ? ? ? ? ? ? ? ? ? ? ? ? ? ? ? ? ? ? ? ? ? ? ? ? ? ? ? ? 0 ? ? ? ? ? ? ? ? ? ? ? ? ? ? ? ? ? ? ? ? ? ? ? ? 0 0 ? ? 0 0 0 - 0 ? ? ? 0 0 0 0 ? ? ? ? ? 0 ? ? ? ? - ? 0 0 0 1 0 1 0 0 0 2 0 1 ? 0 0 0 2 0 2 2 ? ? 0 0 0 ? ? ? ? ? ? ? ? ? ? ? ? ? ? ? ? ? ? ? ? 0 ? ? ? ? ? ? ? ? ? ? ? ? ? ? ? ? ? ? ? ? ? ? ? ? ? ? ? ? ? ? ? ? 0 0 1 0 0 0 0 0 0 0 ? ? 0 0 0 0 0 ? 0 ? ? ? ? ? ? ? ? 0 0 1 0 0 0 0 0 0 1 0 0 1

Parapsicephalus_purdoni ? ? ? 0 ? ? ? 0 1 0 1 0 0 0 0 1 1 1 0 0 0 - 2 0 3 0 0 0 2 1 0 - - - - - - - 0 1 1 1 2 0 0 1 0 0 0 - - 1 0 0 0 - 0 0 0 1 0 0 0 - 1 0 0 0 0 0 0 1 0 0 0 1 0 0 0 0 - - 0 1 0 0 0 ? ? ? ? ? ? ? ? ? ? ? ? ? ? ? ? ? ? ? ? ? ? ? ? ? ? ? ? 0 ? ? ? ? ? ? 0 ? 1 ? ? ? ? [01] 1 ? ? ? ? ? ? ? 0 ? ? ? ? ? ? ? ? ? ? ? ? ? ? ? ? ? ? ? ? ? ? ? ? ? ? ? ? ? ? ? ? ? ? ? ? ? ? ? ? ? ? ? ? ? ? ? ? ? ? ? ? ? ? ? ? ? ? ? ? ? ? ? ? ? ? ? ? ? ? ? ? ? ? ? ? ? ? ? ? ? ? ? ? ? 0 ? ? ? 1 ? ? ?

Campylognathoides_liasicus 1 - 1 0 0 - 0 0 0 0 1 0 0 1 0 1 1 1 0 0 0 - 0 0 2 1 1 0 2 1 0 - - - - - - - 0 2 1 0 2 0 0 0 0 0 0 - - 1 0 0 0 - 0 0 0 2 0 0 0 - 0 0 0 1 0 0 0 0 0 0 0 1 0 0 0 0 - - 0 1 ? ? 0 1 2 2 0 0 0 - 0 0 0 0 0 0 1 - 0 0 0 0 - 1 0 1 3 0 - - 1 0 0 0 - 0 0 0 0 2 1 1 1 0 0 0 1 1 - - 1 0 0 1 0 0 1 0 ? 0 0 0 0 0 0 0 0 0 0 0 0 2 0 0 0 0 0 0 2 1 1 1 0 0 0 1 0 0 1 1 1 1 ? ? 1 0 1 0 0 0 0 0 1 ? 0 2 0 0 0 0 1 0 0 0 1 1 0 1 1 3 1 0 ? 0 0 ? 0 1 0 1 1 0 0 0 2 1 0 1 0 0 0 0 0 0 1 0 0 1

Campylognathoides_zitteli 1 - 1 ? 0 - 0 0 0 0 1 0 ? 1 0 1 1 1 0 0 0 - 0 0 ? ? ? 0 2 1 0 - - - - - - - 0 2 1 0 2 0 0 0 0 0 0 - - 1 0 0 0 - 0 0 ? ? 0 0 0 - 0 0 0 1 0 0 ? 0 ? ? ? ? 0 ? 0 ? ? ? ? ? ? ? 0 1 2 2 0 0 0 - 0 0 0 0 0 0 1 - 0 0 0 ? ? 1 0 1 3 0 - - 1 0 0 0 - 0 0 0 0 2 1 1 1 0 0 0 1 1 - - 1 0 0 1 0 ? ? ? ? 0 0 0 0 0 ? ? 0 0 0 0 0 2 0 ? 0 0 0 0 2 ? 1 1 0 0 0 1 ? 0 1 1 1 1 ? ? 1 0 1 0 ? ? 0 0 1 ? ? 2 0 0 0 0 1 0 ? 0 1 1 0 1 1 3 1 0 1 0 0 1 0 1 0 1 1 ? 0 0 2 1 0 1 0 0 0 0 0 0 1 0 0 1

Sericipterus_wucaiwanensis 3 1 1 ? 1 0 1 0 1 0 1 0 1 1 1 1 1 1 1 0 1 - ? ? ? ? ? 0 ? 1 1 0 0 1 ? 0 0 1 0 1 1 ? 2 0 0 0 0 ? 0 - - ? ? ? 0 - 0 ? 0 2 0 0 1 - 1 ? 0 0 ? ? 1 1 ? 0 ? ? 0 0 ? 0 - - ? ? ? ? 1 1 ? ? ? ? ? ? 1 0 0 1 0 0 1 - 0 0 0 0 - 0 0 ? 3 1 ? - 1 0 2 0 - 2 2 1 1 2 2 0 1 1 1 1 2 1 - - 0 0 ? ? 0 ? 1 0 0 0 0 0 0 ? ? ? ? 0 ? ? ? ? ? ? 0 0 ? 0 2 ? ? ? ? ? ? ? 0 0 0 1 1 1 ? 0 1 0 3 0 0 1 0 0 ? ? ? ? ? ? ? ? ? ? ? 0 2 2 0 1 ? ? ? ? ? ? ? ? ? ? ? ? ? ? ? ? ? ? 0 1 0 0 0 0 0 ? 1 0 0 1

Angustinaripterus_longicephalus 3 1 1 ? 1 0 1 0 1 0 1 0 1 1 1 1 1 1 1 0 1 - 0 0 2 1 1 0 2 ? 1 0 0 1 ? 0 0 1 0 1 1 0 2 0 0 0 0 0 0 - - 1 ? 0 ? ? 0 0 0 2 1 0 1 - 1 0 0 0 0 0 1 1 ? ? ? ? ? ? ? ? ? ? 0 1 ? ? ? 1 1 2 0 1 0 - 1 0 0 1 0 1 1 - ? ? ? 0 - 0 0 1 3 1 - - 1 0 2 0 - 2 2 1 1 1 2 0 1 1 1 1 2 1 - - 0 0 0 ? 0 ? ? ? ? ? ? ? ? ? ? ? ? ? ? ? ? ? ? ? ? ? ? ? ? ? ? ? ? ? ? ? ? ? ? ? ? ? ? ? ? ? ? ? ? ? ? ? ? ? ? ? ? ? ? ? ? ? ? ? ? ? ? ? ? ? ? ? ? ? ? ? ? ? ? ? ? ? ? ? ? ? ? ? ? ? 0 ? ? ? ? ? ? ?

Harpactognathus_gentryii 3 0 1 0 1 0 1 0 1 2 1 0 1 1 1 1 ? 1 1 0 ? - ? ? ? ? ? 0 ? ? 1 0 0 1 ? 0 0 1 0 0 1 ? 2 0 0 ? 0 ? ? ? ? ? ? ? ? ? ? ? ? ? ? ? ? - ? ? ? ? ? ? ? ? ? ? ? ? ? ? 0 0 - - 0 1 ? ? ? 1 ? ? ? ? ? ? ? ? ? ? ? ? ? ? ? ? ? ? ? ? ? ? ? ? ? ? ? 0 2 0 - ? ? ? 1 2 2 0 ? ? ? [01] 2 ? ? ? 0 0 ? ? 0 ? ? ? ? ? ? ? ? ? ? ? ? ? ? ? ? ? ? ? ? ? ? ? ? ? ? ? ? ? ? ? ? ? ? ? ? ? ? ? ? ? ? ? ? ? ? ? ? ? ? ? ? ? ? ? ? ? ? ? ? ? ? ? ? ? ? ? ? ? ? ? ? ? ? ? ? ? ? ? ? ? ? ? ? ? 0 ? ? ? ? ? ? ?

Cacibupteryx_caribensis ? ? 1 0 ? ? ? 0 1 0 1 0 1 1 1 1 1 0 1 0 1 - 0 0 2 0 0 0 ? 1 0 - - - - - - - 0 0 1 0 2 0 0 0 0 0 0 - - 0 0 0 0 - 0 0 0 1 0 0 1 - ? 0 0 0 0 0 0 ? 0 0 0 1 0 0 0 0 - - 0 1 0 0 ? 1 ? ? ? ? ? ? ? ? ? ? ? ? ? ? ? ? ? ? ? ? ? ? ? ? ? ? ? 0 2 0 - ? ? ? 0 ? 2 ? ? ? ? [01] 2 ? ? ? ? 0 0 ? 0 ? ? ? ? ? ? ? ? ? ? ? ? ? ? ? ? ? ? ? ? ? ? ? ? ? ? ? ? ? ? ? ? ? ? ? ? ? ? ? ? ? ? ? ? ? ? ? 0 ? ? ? ? ? ? ? ? ? ? ? ? ? ? ? ? ? ? ? ? ? ? ? ? ? ? ? ? ? ? ? ? ? ? ? ? 0 ? ? ? ? 1 ? ? ?

Rhamphorhynchus_muensteri 2 - 1 0 0 - 0 0 0 0 1 0 1 1 1 1 1 0 1 0 1 - 0 0 2 1 1 0 2 1 0 - - - - - - - 0 0 1 0 2 0 0 0 0 0 0 - - 0 0 0 0 - 0 0 1 2 2 0 1 - 1 0 0 0 0 0 - 1 0 0 0 1 0 0 0 0 - - 0 1 0 0 1 1 0 2 0 0 0 - 1 0 0 0 0 0 1 - 0 0 0 0 - 0 1 1 3 1 - - 1 0 2 0 - 2 2 1 0 2 2 0 1 1 0 2 2 1 - - 0 0 0 1 0 0 1 1 0 0 0 0 0 0 0 0 0 0 1 0 0 2 0 0 0 0 0 0 2 1 1 0 1 0 0 [12] 0 0 0 1 1 1 0 0 1 0 6 0 0 0 0 1 0 0 0 2 0 0 1 0 1 0 1 0 1 2 0 1 1 [12] 0 1 1 0 0 1 0 1 0 1 1 0 0 1 2 [12] 0 1 0 0 0 0 0 0 1 0 0 1

Qinglongopterus_guoi 3 ? 1 ? 0 - ? 0 ? 0 ? 0 1 ? 1 ? ? 1 1 0 1 - 0 0 2 ? 1 ? 2 ? ? ? ? ? ? ? ? ? ? 0 ? ? 2 0 0 0 0 ? ? ? ? ? ? ? ? ? 0 ? ? 2 0 0 ? - ? 0 0 ? 0 0 ? ? ? 0 0 1 ? ? ? ? ? ? 0 ? ? ? ? 1 1 2 0 0 ? ? 1 ? 0 ? 0 ? 1 - ? ? ? ? ? 0 ? 1 3 0 - - 1 0 2 0 - 2 0 1 1 2 2 0 1 1 1 1 ? 1 - - 0 0 0 1 ? ? 1 0 ? ? ? 0 0 0 ? 0 0 ? 1 ? 0 2 0 0 0 0 0 0 2 ? ? ? ? 0 0 ? ? ? ? 1 1 1 ? ? 1 0 3 0 0 0 ? 0 0 ? ? 3 ? 0 1 0 1 ? 1 0 0 ? 0 1 1 1 0 1 ? ? ? ? ? 1 ? 1 1 0 0 1 2 1 0 1 0 0 0 0 0 0 1 0 0 1

Nesodactylus_hesperius ? ? ? ? ? ? ? ? ? ? ? ? ? ? ? ? ? ? ? ? ? ? ? ? ? ? 0 ? ? ? ? ? ? ? ? ? ? ? ? ? ? ? ? ? ? ? ? ? ? ? ? ? ? ? ? ? ? ? ? 1 ? 0 ? ? ? ? ? ? ? ? ? ? ? ? ? ? ? ? ? ? ? ? ? ? ? ? ? ? ? ? ? ? ? ? ? ? ? ? ? ? ? ? ? ? ? ? ? ? ? ? ? ? ? ? ? ? ? ? ? ? ? ? ? ? ? ? ? ? ? ? ? ? ? ? ? ? ? ? ? ? ? 1 ? 0 0 0 0 ? ? ? ? ? ? ? 0 2 0 ? 0 0 ? 0 2 ? 1 1 0 0 0 2 0 0 ? 1 1 1 ? ? 1 0 6 0 0 0 0 1 0 0 ? ? 0 0 1 0 1 0 ? 0 1 ? ? ? 0 0 ? 1 0 0 0 1 0 ? ? 1 ? ? ? ? ? ? 0 1 0 0 ? 0 0 0 1 0 0 1

Dorygnathus_banthensis 1 - 1 0 0 - 0 0 0 0 1 0 1 1 1 1 1 1 1 0 1 - 0 0 2 1 1 0 2 1 0 - - - - - - - 0 0 1 0 2 0 0 0 0 0 0 - - 1 0 0 0 - 0 0 0 2 0 0 1 - 1 0 0 0 0 0 0 1 0 0 0 1 0 0 0 0 - - 0 1 0 0 1 1 1 2 0 0 1 0 1 0 0 0 0 0 1 - 0 2 0 0 - 0 0 1 3 1 - - 1 0 2 0 - 2 0 1 0 2 2 0 1 1 0 1 1 1 - - 1 0 0 1 0 0 1 0 ? 0 0 0 0 0 0 0 0 0 1 0 0 2 0 0 0 0 0 0 2 1 1 0 1 0 0 3 0 0 1 1 1 1 0 0 1 0 3 0 0 0 0 1 0 0 0 3 0 0 1 0 1 0 1 0 0 2 0 1 1 [13] 0 1 1 0 0 1 0 1 0 1 1 0 0 1 2 2 0 1 0 0 0 0 0 0 1 0 0 1

Scaphognathus_crassirostris 1 - 1 ? 0 - 0 0 0 0 1 0 3 1 0 1 1 1 1 0 0 - 0 0 2 1 1 0 2 1 0 - - - - - - - 0 1 1 0 2 0 0 0 0 0 0 - - 1 0 0 0 - 0 0 0 2 0 0 1 - 1 0 0 0 0 0 0 1 0 ? ? 1 0 ? ? ? ? ? 0 1 ? 0 0 1 1 2 0 0 0 - 1 0 0 0 0 0 1 - ? ? ? ? ? 0 0 1 3 1 - - 1 0 2 0 - 2 0 1 0 2 2 0 1 0 0 0 1 1 - - 1 0 0 0 0 0 ? ? 0 0 0 0 0 0 0 0 0 0 0 0 0 2 0 0 0 0 0 0 2 1 1 ? ? 0 0 2 0 0 0 1 1 0 ? ? 1 0 3 0 0 0 0 1 0 0 0 2 0 0 1 0 1 0 1 0 0 2 ? 1 1 1 0 1 [01] 0 0 0 0 1 0 1 1 0 0 1 2 2 0 1 0 0 0 0 0 0 1 0 0 1

Sordes_pilosus 1 - 1 0 0 - 0 0 0 0 1 0 1 1 0 1 1 1 0 0 0 - 0 0 2 1 1 0 2 1 0 - - - - - - - 0 1 1 0 2 0 0 1 0 0 0 - - 1 0 0 0 - 0 0 1 2 1 0 1 - 1 0 0 0 1 0 - 1 0 0 0 1 0 0 0 0 - - 0 1 ? 0 0 1 1 0 0 0 0 - 1 0 0 0 0 0 1 - 0 0 0 1 0 1 0 1 3 0 - - 1 0 2 0 - 3 0 0 0 2 2 0 1 0 0 0 1 1 - - 1 0 0 0 0 0 1 1 ? 0 0 0 0 0 0 0 0 0 1 0 0 2 0 0 0 0 0 0 2 ? 1 1 0 0 0 4 0 0 1 1 1 1 ? ? 1 0 0 0 0 0 0 0 0 0 ? [45] 0 0 1 0 1 0 1 0 0 2 0 1 1 1 0 1 1 0 0 0 0 1 0 1 1 0 0 1 2 2 0 1 0 0 0 0 0 0 1 0 0 1

Darwinopterus_modularis 1 - 1 0 0 - 0 0 0 0 1 0 1 1 - 1 1 - - 1 0 0 0 0 4 1 1 0 - 1 1 1 1 2 2 0 0 0 0 - - - - 0 0 1 0 1 0 - - 1 0 0 0 - 0 0 1 2 1 0 1 0 1 0 0 1 1 0 - 1 1 ? ? 1 0 ? 0 1 0 0 0 1 ? ? ? 1 1 0 0 0 0 - 1 0 0 0 0 0 1 - ? ? ? ? ? 1 0 1 0 0 - - 1 0 2 0 - 3 0 0 0 2 2 0 1 0 1 0 3 1 - - 2 0 0 0 0 1 ? ? ? 1 0 1 0 0 ? 0 0 ? 1 ? 0 2 0 0 0 0 0 0 2 ? 1 ? ? 0 0 2 0 0 ? 1 1 0 ? ? ? 1 2 0 0 0 ? ? ? ? 0 4 ? 0 1 0 ? ? 0 ? 0 2 0 1 1 3 1 1 0 0 0 0 0 1 ? 1 1 0 0 1 2 2 0 1 0 0 1 0 1 0 1 0 0 1

Wukongopterus_lii 1 - 1 ? 0 - 0 0 0 0 1 0 1 1 - 1 ? - - 1 ? 0 ? ? ? ? ? 0 - ? ? ? ? ? ? ? ? ? 0 - - - - ? ? ? ? ? ? ? ? ? ? ? ? ? ? ? ? 2 ? ? ? 0 1 ? ? ? ? ? ? 1 ? ? ? ? ? ? ? ? ? ? 0 1 ? ? ? 1 1 0 0 0 0 - 1 0 0 0 0 0 1 - ? ? ? ? ? 1 0 1 0 0 - - 1 0 2 0 - 3 0 0 0 2 2 0 1 0 1 0 3 1 - - 2 0 0 0 0 ? ? 0 ? 1 0 ? 0 0 1 0 0 0 1 0 0 2 0 0 0 0 0 0 2 ? ? ? ? 0 0 2 ? ? 0 1 1 0 ? ? 1 ? ? ? ? ? ? ? 0 ? ? 4 ? 0 1 0 1 0 0 ? ? ? ? ? ? ? 0 1 ? 0 0 0 0 1 ? 1 1 ? 0 1 2 2 0 1 0 0 0 0 1 0 1 0 0 1

Pterorhynchus_wellnhoferi 1 - 1 0 0 - 0 0 0 0 1 0 1 1 - 1 1 - - 1 0 0 0 0 4 1 1 0 - ? 1 1 0 1 1 0 0 0 0 - - - - 0 0 1 0 0 0 - - ? ? 0 0 - 0 0 ? 2 2 0 1 0 1 0 0 0 1 0 ? 1 0 0 0 1 0 ? ? ? ? ? ? ? ? ? ? 1 1 0 0 0 0 - 1 0 0 0 0 0 1 - ? ? ? ? ? 1 0 1 0 0 - - 1 0 2 0 - 3 0 0 0 2 2 0 1 0 1 0 1 1 - - 1 0 0 0 0 1 ? 0 ? ? ? 0 0 ? ? ? ? 0 1 0 0 2 0 0 ? 0 ? ? ? ? ? ? ? ? ? ? ? ? ? 1 1 0 ? ? 1 0 2 0 0 0 ? ? ? ? ? 0 0 0 1 0 1 0 ? ? 0 2 0 1 ? ? 0 1 0 0 0 1 0 1 ? ? ? 0 ? ? 2 ? 0 1 0 0 0 1 0 1 0 0 1

Changchengopterus_pani ? ? ? ? ? ? ? ? ? ? ? ? ? ? ? ? ? ? ? ? ? ? ? ? ? ? ? ? ? ? ? ? ? ? ? ? ? ? ? ? ? ? ? ? ? ? ? ? ? ? ? ? ? ? ? ? ? ? ? ? ? ? ? ? ? ? ? ? ? ? ? ? ? ? ? ? ? ? ? ? ? ? ? ? ? ? ? ? ? ? ? ? ? ? ? ? ? ? ? ? ? ? ? ? ? ? ? ? ? ? ? ? ? ? ? ? ? ? ? ? ? ? ? ? ? ? ? ? ? ? ? ? ? ? ? ? ? ? ? ? ? 0 ? 1 0 0 0 0 1 ? 0 ? ? ? 0 1 0 ? 0 0 ? 0 2 ? ? ? ? 0 0 ? 0 0 ? 1 1 1 ? ? ? 0 0 0 0 0 ? ? ? ? ? 4 0 0 1 0 1 ? ? 0 0 2 0 1 ? ? 0 1 ? 0 ? ? ? 1 ? 1 1 ? 1 1 2 1 0 1 0 0 ? 0 0 0 1 0 0 1 0 1 0 0 0 1 0 1 0 0 1

Kryptodrakon_progenitor ? ? ? ? ? ? ? ? ? ? ? ? ? ? ? ? ? ? ? ? ? ? ? ? ? ? ? ? ? ? ? ? ? ? ? ? ? ? ? ? ? ? ? ? ? ? ? ? ? ? ? ? ? ? ? ? ? ? ? ? ? ? ? ? ? ? ? ? ? ? ? ? ? ? ? ? ? ? ? ? ? ? ? ? ? ? ? ? ? ? ? ? ? ? ? ? ? ? ? ? ? ? ? ? ? ? ? ? ? ? ? ? ? ? ? ? ? ? ? ? ? ? ? ? ? ? ? ? ? ? ? ? ? ? ? ? ? ? ? ? ? ? ? ? ? ? ? ? ? ? ? ? ? 0 ? ? ? ? ? ? ? ? 2 ? ? ? ? ? ? ? 0 0 ? ? 1 ? ? ? ? 0 ? ? ? ? ? ? 0 0 0 ? 1 0 ? 0 0 0 ? 0 0 ? ? ? ? ? ? ? ? ? ? ? ? ? ? ? ? ? ? ? ? ? ? ? ? ? ? ? ? ? 1 0 ? ?

Germanodactylus_cristatus 2 - 1 0 0 - 0 0 0 0 1 0 1 1 - 1 1 - - 1 0 0 0 0 4 1 1 0 - 1 1 1 0 3 2 0 0 0 0 - - - - 0 0 1 1 0 0 - - 1 1 0 0 - 2 0 1 3 2 0 1 0 1 0 0 0 1 0 - 1 1 ? ? ? 0 ? 1 ? ? ? ? ? ? ? ? 1 1 1 0 0 0 - 1 0 0 0 0 0 1 - 0 ? 0 ? ? 1 0 1 3 0 - - 1 0 2 0 - 3 0 0 0 1 2 0 1 0 0 0 1 1 - - 0 1 0 1 0 0 ? ? ? ? ? 0 0 1 1 0 0 0 1 0 ? ? 0 0 0 0 0 0 2 ? ? ? ? 0 0 2 0 0 0 1 1 1 0 ? 1 0 5 0 ? ? 0 0 0 ? ? 4 ? ? 1 1 0 0 1 0 ? 3 0 1 1 3 0 1 0 0 0 1 0 1 0 1 1 0 0 1 3 3 0 1 0 0 1 1 1 0 0 1

Germanodactylus_rhamphastinus 2 - 1 0 0 - 0 0 0 0 1 0 1 1 - 1 1 - - 1 0 0 0 0 4 1 1 0 - 1 1 1 0 3 2 0 0 0 0 - - - - 0 0 1 1 0 0 - - 1 1 0 0 - 2 0 1 3 2 0 1 0 1 0 0 0 1 0 - 1 1 ? ? ? 0 ? 1 ? ? ? ? ? ? ? ? 1 1 1 0 0 0 - ? 0 0 0 0 0 1 - 0 ? 0 ? - 1 0 1 3 0 - - 1 0 2 0 - 3 0 0 0 1 2 0 1 0 0 0 1 1 - - 1 1 0 0 0 0 ? ? ? 1 0 0 0 1 ? ? 0 ? 1 0 ? ? ? 0 0 0 0 ? 2 0 ? 0 0 0 0 2 ? 0 ? 1 1 1 ? ? 1 ? ? ? ? ? 0 ? 0 0 ? 4 ? ? 1 ? 0 0 1 0 ? 3 0 1 1 3 0 ? 0 0 0 ? 0 1 0 1 1 0 0 1 ? ? 0 1 0 0 1 1 1 0 0 1

Normannognathus_wellnhoferi 1 - 0 0 0 - 0 0 0 ? 1 0 ? 2 ? 1 ? ? ? ? ? ? ? ? ? ? ? ? ? ? 1 1 2 2 ? 0 0 0 ? - ? ? ? ? ? ? ? ? ? ? ? ? ? ? ? ? ? ? ? ? ? ? ? ? ? ? ? ? ? ? ? ? ? ? ? ? ? ? 1 0 - - ? ? ? ? ? 1 ? 0 2 0 0 - 1 ? 0 0 0 0 ? ? 1 0 0 ? ? ? ? ? ? ? ? ? ? 0 2 0 - 2 0 1 0 1 1 0 1 0 0 ? 1 1 - - 1 1 0 0 0 ? ? ? ? ? ? ? ? ? ? ? ? ? ? ? ? ? ? ? ? ? ? ? ? ? ? ? ? ? ? ? ? ? ? ? ? ? ? ? ? ? ? ? ? ? ? ? ? ? ? ? ? ? ? ? ? ? ? ? ? ? ? ? ? ? ? ? ? ? ? ? ? ? ? ? ? ? ? ? ? ? 0 1 0 0 1 1 1 0 0 1

Dsungaripterus_weii 3 0 0 0 0 - 1 0 0 0 1 0 0 1 - 1 1 - - 1 0 0 0 1 2 1 1 0 - 1 1 1 2 3 3 0 1 0 1 - - - - 1 - - - - 1 1 2 0 0 0 1 2 2 0 1 2 2 0 1 0 1 1 0 1 1 0 - 1 1 1 1 0 1 1 0 1 0 0 0 1 0 1 1 1 0 1 0 0 0 - 1 0 0 1 0 0 1 - 1 0 0 1 0 0 0 1 0 1 - - 1 0 2 0 - 1 0 0 0 2 2 0 1 0 1 0 4 1 - - 0 2 0 1 0 1 0 1 1 0 0 1 0 1 0 1 1 1 1 1 1 0 0 1 0 0 1 1 2 1 1 1 0 0 0 1 ? ? ? 0 ? 0 ? ? 1 0 8 0 1 1 1 1 1 ? 0 ? 1 0 ? 1 0 0 1 1 2 3 ? 1 ? ? ? ? 1 1 1 1 ? 0 0 1 2 1 0 ? ? ? 0 1 0 0 1 1 1 1 0 1

Domeykodactylus_ceciliae ? ? ? ? ? ? ? ? ? ? ? 0 ? ? ? ? ? ? ? ? ? ? ? ? ? ? ? 0 ? ? 1 ? 2 ? ? ? 1 0 ? ? ? ? ? ? ? ? ? ? ? ? ? ? ? ? ? ? ? ? ? ? ? ? ? ? ? ? ? ? ? ? ? ? ? ? ? ? ? ? ? ? ? ? ? ? ? ? ? 1 1 ? 0 ? ? ? 1 0 ? ? 0 0 1 - ? ? ? 1 0 ? 0 ? ? 1 ? - ? 0 2 0 - 1 ? ? 0 2 2 0 ? ? ? 0 4 ? ? ? ? ? 0 1 ? ? ? ? ? ? ? ? ? ? ? ? ? ? ? ? ? ? ? ? ? ? ? ? ? ? ? ? ? ? ? ? ? ? ? ? ? ? ? ? ? ? ? ? ? ? ? ? ? ? ? ? ? ? ? ? ? ? ? ? ? ? ? ? ? ? ? ? ? ? ? ? ? ? ? ? ? ? ? ? ? ? 0 1 0 0 1 1 1 0 0 1

Noripterus_parvus 3 0 1 0 0 - 1 0 0 0 1 0 0 1 - 1 1 - - 1 0 0 0 1 2 0 1 0 - 1 1 1 1 3 3 0 1 0 1 - - - - 1 - - - - 1 1 2 0 0 0 1 2 2 0 1 2 2 0 1 ? 1 1 0 1 1 0 - 1 1 1 1 0 1 ? 0 1 0 0 0 1 0 ? 1 1 1 1 0 0 0 - 1 0 0 1 0 0 1 - 1 0 0 1 1 0 0 1 0 1 - - 1 0 2 0 - 1 0 0 0 2 2 0 1 0 1 0 3 1 - - 0 1 0 1 0 1 0 1 1 0 0 1 0 1 0 1 1 1 ? ? ? ? ? ? 0 0 ? 1 2 ? ? ? ? 0 0 ? 0 0 0 0 1 1 0 0 1 0 8 0 1 1 1 1 1 0 0 4 1 ? 1 1 0 0 ? 0 2 ? ? 1 ? ? ? ? ? ? ? ? ? 0 0 1 2 1 0 ? ? ? 0 1 0 0 1 1 1 1 0 1

Noripterus_complicidens ? ? ? ? ? - ? ? ? ? ? 0 0 ? ? ? ? ? ? ? ? ? ? ? ? ? ? ? ? ? ? ? ? ? ? ? ? ? ? ? ? ? ? ? ? ? ? ? ? ? ? ? ? ? ? ? ? ? ? ? ? ? ? ? ? ? ? ? ? ? ? ? ? ? ? ? ? ? ? ? ? ? ? ? ? ? ? 1 ? ? 0 0 0 - 1 0 0 ? 0 0 ? ? 1 0 0 1 1 0 ? ? ? 1 - - ? 0 2 0 - 1 0 ? 0 2 2 0 ? ? 1 ? 3 1 - - ? ? ? ? ? ? 0 1 1 0 0 1 0 1 ? ? 1 ? 1 1 1 0 0 ? ? ? ? 1 2 ? ? ? ? 0 0 ? 0 0 1 0 1 1 1 0 1 0 8 0 1 1 1 1 1 0 0 ? ? 0 1 1 0 0 ? 0 2 1 0 1 1 3 ? 1 1 1 1 1 1 0 0 1 2 1 0 ? 4 - 0 1 0 0 1 1 1 1 0 1

Kepodactylus_insperatus ? ? ? ? ? ? ? ? ? ? ? ? ? ? ? ? ? ? ? ? ? ? ? ? ? ? ? ? ? ? ? ? ? ? ? ? ? ? ? ? ? ? ? ? ? ? ? ? ? ? ? ? ? ? ? ? ? ? ? ? ? ? ? ? ? ? ? ? ? ? ? ? ? ? ? ? ? ? ? ? ? ? ? ? ? ? ? ? ? ? ? ? ? ? ? ? ? ? ? ? ? ? ? ? ? ? ? ? ? ? ? ? ? ? ? ? ? ? ? ? ? ? ? ? ? ? ? ? ? ? ? ? ? ? ? ? ? ? ? ? 0 1 0 1 2 1 0 1 ? ? ? ? ? ? ? ? ? ? ? ? ? ? ? ? ? ? ? ? ? ? 0 0 1 1 1 1 ? 0 1 0 5 0 0 0 ? ? 0 ? ? ? ? ? ? 1 0 0 ? ? 0 ? ? ? ? ? 0 ? ? ? ? ? ? 1 ? ? ? ? ? ? ? ? ? ? ? ? ? ? 1 ? ? ?

Gnathosaurus_subulatus 1 - 1 0 1 1 0 0 0 2 1 0 1 ? - 1 1 - - 1 1 0 0 0 4 1 1 0 - 1 1 1 0 3 1 0 0 0 0 - - - - 0 0 1 0 0 0 - - 1 0 0 0 - 1 1 1 3 2 0 1 0 1 0 0 0 1 0 - 1 2 1 1 0 0 ? ? 0 - - 0 1 0 1 ? 1 1 0 2 1 0 - 1 0 0 ? 0 0 1 - ? ? ? ? ? 1 ? ? ? 0 - - 1 0 2 0 - 2 0 1 1 1 2 0 ? 0 1 2 3 1 - - 1 0 0 ? 0 ? ? ? ? ? ? ? ? ? ? ? ? ? ? ? ? ? ? ? ? ? ? ? ? ? ? ? ? ? ? ? ? ? ? ? ? ? ? ? ? ? ? ? ? ? ? ? ? ? ? ? ? ? ? ? ? ? ? ? ? ? ? ? ? ? ? ? ? ? ? ? ? ? ? ? ? ? ? ? ? ? ? ? ? ? ? ? ? ? ? ?

Gnathosaurus_macrurus ? ? ? ? ? 1 0 ? 0 ? ? 0 1 ? ? ? ? ? ? ? ? ? ? ? ? ? ? ? ? ? ? ? ? ? ? ? ? ? ? ? ? ? ? ? ? ? ? ? ? ? ? ? ? ? ? ? ? ? ? ? ? ? ? ? ? ? ? ? ? ? ? ? ? ? ? ? ? ? ? ? ? ? ? ? ? ? ? ? 1 0 2 1 0 - 1 0 ? ? 0 0 1 - 1 0 0 0 - 1 0 ? 2 ? ? ? ? 0 2 0 - ? ? ? 1 1 2 0 ? ? ? 2 3 ? ? ? ? ? 0 0 ? ? 0 1 ? 2 2 1 0 1 ? ? ? ? ? ? ? ? ? ? ? ? ? ? ? ? ? ? ? ? ? ? ? ? ? ? ? ? ? ? ? ? ? ? ? ? ? ? ? ? ? ? ? ? ? ? ? ? ? ? ? ? ? ? ? ? ? ? ? ? ? ? ? ? ? ? ? ? ? ? ? ? ? ? ? ? ? ? ? ? ? ?

Plataleorhynchus_streptophorodon 1 - 1 0 1 0 0 0 0 2 ? 0 1 ? ? 1 ? - - 1 ? ? ? ? ? ? ? ? - ? ? ? ? ? ? ? ? ? ? ? - ? ? ? ? ? ? ? ? ? ? ? ? ? ? ? ? ? 1 ? ? ? ? ? ? ? ? ? ? ? - ? ? ? ? ? ? ? 1 0 - - 0 ? ? ? ? ? 1 ? ? ? ? ? ? ? ? ? ? ? ? ? ? ? ? ? ? ? ? ? ? ? ? ? ? 0 2 ? ? ? ? ? 1 1 2 0 ? ? ? 1 3 ? ? ? 1 0 0 0 0 ? ? ? ? ? ? ? ? ? ? ? ? ? ? ? ? ? ? ? ? ? ? ? ? ? ? ? ? ? ? ? ? ? ? ? ? ? ? ? ? ? ? ? ? ? ? ? ? ? ? ? ? ? ? ? ? ? ? ? ? ? ? ? ? ? ? ? ? ? ? ? ? ? ? ? ? ? ? ? ? ? ? ? ? ? ? ? ? ? ? ?

Huanhepterus_quingyangensis 1 - 1 ? 1 0 0 0 0 0 1 0 1 ? ? ? ? ? ? ? ? ? ? ? ? ? ? ? ? ? 1 1 0 3 ? 0 0 0 ? ? ? ? ? ? ? ? ? ? ? ? ? ? ? ? ? ? ? ? ? ? ? ? ? ? ? ? ? ? ? ? ? ? ? ? ? ? ? ? ? ? ? ? ? ? ? ? ? 1 1 0 0 1 0 - 1 0 0 0 0 0 ? ? ? ? ? ? ? ? ? ? ? 0 - - ? 0 2 0 - 2 0 1 0 1 2 0 1 0 1 1 3 1 - - 1 0 0 0 0 ? ? ? ? 2 2 ? 0 1 1 0 0 0 1 ? ? 0 ? ? ? 0 ? ? 2 ? ? ? ? ? ? 1 ? 0 ? 1 ? 1 ? ? 1 0 5 0 ? ? ? ? ? ? ? ? ? 0 ? 1 ? ? ? ? ? ? ? ? 1 3 0 1 0 0 1 ? 0 1 ? 1 ? ? 0 ? ? ? 0 1 0 0 0 0 1 0 0 1

Moganopterus_zhuiana 1 - 1 0 0 - 0 0 0 0 1 ? ? 2 - 1 1 - - 1 0 1 0 0 4 1 1 0 - 1 1 1 0 1 0 0 0 ? 0 - - - - 0 0 1 1 0 0 - - 0 ? 0 1 3 1 1 ? 3 2 0 1 ? ? 0 ? 0 ? 0 ? ? 2 ? 1 0 0 ? ? ? ? ? ? ? ? ? ? ? 1 2 0 0 0 - 1 0 0 0 0 0 1 - ? ? ? ? ? 0 0 0 2 0 - - 1 0 2 0 - 2 0 1 0 1 2 0 1 1 1 1 1 1 - - 1 0 0 0 0 ? ? ? ? 2 2 1 ? ? ? ? ? ? ? ? ? ? ? ? ? ? ? ? ? ? ? ? ? ? ? ? ? ? ? ? ? ? ? ? ? ? ? ? ? ? ? ? ? ? ? ? ? ? ? ? ? ? ? ? ? ? ? ? ? ? ? ? ? ? ? ? ? ? ? ? ? ? ? ? ? ? ? ? ? ? ? ? ? ? ? ?

Elanodactylus_prolatus ? ? ? ? ? ? ? ? ? ? ? ? ? ? ? ? ? ? ? ? ? ? ? ? ? ? ? ? ? ? ? ? ? ? ? ? ? ? ? ? ? ? ? ? ? ? ? ? ? ? ? ? ? ? ? ? ? ? ? ? ? ? ? ? ? ? ? ? ? ? ? ? ? ? ? ? ? ? ? ? ? ? ? ? ? ? ? ? ? ? ? ? ? ? ? ? ? ? ? ? ? ? ? ? ? ? ? ? ? ? ? ? ? ? ? ? ? ? ? ? ? ? ? ? ? ? ? ? ? ? ? ? ? ? ? ? ? ? ? ? ? ? ? 2 2 1 0 1 1 0 0 0 ? ? ? ? ? 0 0 0 0 0 2 ? 1 0 1 0 0 1 ? 0 1 1 1 1 ? 0 1 0 5 0 0 0 0 0 0 ? 0 4 0 0 1 1 0 0 0 ? 0 3 0 1 1 3 0 1 0 0 1 1 0 ? 0 1 1 ? 0 1 3 3 0 1 0 0 0 1 0 0 1

Ctenochasma_elegans 1 - 1 0 0 - 0 0 0 0 0 1 1 2 - 1 1 - - 1 0 0 0 0 4 1 1 0 - 1 [01] 1 0 3 ? 0 0 0 0 - - - - 0 0 1 2 0 0 - - 1 1 0 0 - 1 1 1 3 2 0 1 0 1 0 0 0 1 0 - 1 2 ? ? 0 0 ? 1 0 - - 0 ? 0 ? ? 1 1 0 0 0 0 - 1 0 0 0 0 0 1 - 1 0 0 0 - 0 0 1 2 0 - - 1 0 1 0 - 2 0 1 1 1 1 0 1 0 1 2 1 1 - - 1 0 0 0 0 ? 0 ? ? 1 0 1 0 0 1 0 0 0 1 ? ? 0 0 0 0 0 0 0 2 ? ? 0 0 0 0 2 0 0 0 1 1 1 ? ? 1 0 5 0 0 0 0 0 0 ? ? 4 ? ? 1 1 0 0 ? 0 ? 3 0 1 1 3 0 1 0 0 1 ? 0 1 0 1 1 0 0 1 3 3 0 1 0 0 0 0 1 0 0 1

Pterodaustro_guinazui 1 - 0 0 0 - 0 0 0 0 1 1 1 2 - 1 1 - - 1 0 0 0 0 4 1 1 0 - 1 0 - - - - - - - 0 - - - - 0 0 1 2 0 0 - - 1 1 0 0 - 1 1 1 3 1 0 1 ? 1 0 0 0 0 0 - 1 2 ? ? 0 0 ? 1 0 - - 0 ? ? ? ? 1 0 0 0 0 0 - 1 0 0 0 0 0 1 - 1 0 0 ? ? 2 0 0 2 0 - - 1 0 1 0 - 2 0 1 0 2 0 0 2 0 1 0 0 1 - - 0 0 0 0 0 ? 0 1 0 1 0 1 0 ? 1 0 ? ? 1 1 0 0 0 0 0 0 0 0 2 1 1 0 0 0 0 2 0 0 0 ? 1 1 0 ? 1 0 5 0 ? ? 0 0 0 0 0 4 ? ? ? 1 0 0 0 0 0 3 ? ? 1 3 0 1 0 0 1 1 0 1 0 1 1 1 ? 1 3 3 0 1 0 0 0 0 0 0 0 1

Beipiaopterus_chenianus ? ? ? ? ? ? ? ? ? ? ? ? ? ? ? ? ? ? ? ? ? ? ? ? ? ? ? ? ? ? ? ? ? ? ? ? ? ? ? ? ? ? ? ? ? ? ? ? ? ? ? ? ? ? ? ? ? ? ? ? ? ? ? ? ? ? ? ? ? ? ? ? ? ? ? ? ? ? ? ? ? ? ? ? ? ? ? ? ? ? ? ? ? ? ? ? ? ? ? ? ? ? ? ? ? ? ? ? ? ? ? ? ? ? ? ? ? ? ? ? ? ? ? ? ? ? ? ? ? ? ? ? ? ? ? ? ? ? ? ? ? ? ? 1 0 1 0 ? 1 0 0 ? 1 ? 1 0 0 ? 0 0 0 2 2 ? ? ? ? 0 0 ? 0 0 ? 1 1 ? ? ? 1 0 5 0 ? ? ? 0 ? ? ? 4 ? ? ? 1 0 0 0 0 ? ? ? ? ? ? ? ? ? ? ? ? ? 1 ? ? ? ? ? 1 3 3 0 1 0 0 0 1 0 0 1

Gegepterus_changae ? ? 0 ? ? ? ? 0 0 0 1 1 1 2 - 1 1 - - 1 0 0 0 0 ? 1 1 0 - 1 1 ? ? 3 0 0 0 0 0 - - - - 0 0 1 2 0 0 - - 1 1 0 0 - 1 ? 1 3 1 0 1 0 1 0 0 0 0 0 - 1 2 ? ? ? 0 ? 1 0 - - 0 ? ? ? ? 1 0 ? ? ? ? ? 0 0 ? ? 0 0 1 - ? ? 0 ? ? 0 0 0 2 0 - - 1 0 1 0 - 2 0 1 0 ? 1 ? 1 0 1 2 0 1 - - ? 0 ? ? 0 0 0 1 ? 1 0 1 0 0 ? ? ? ? ? ? ? ? ? ? ? ? ? 2 2 ? ? ? ? 0 0 2 ? ? ? ? ? ? ? ? ? ? ? ? ? ? ? ? ? ? ? ? ? [01] ? ? ? 0 ? 0 0 3 ? ? ? ? 0 ? ? ? ? ? ? 1 ? ? ? 0 ? ? ? ? 0 1 0 0 0 1 0 0 1

Feilongus_youngi 1 - 1 0 0 - 0 0 0 0 1 0 ? 2 - 1 1 - - 1 0 1 ? 0 4 1 1 0 - 1 1 1 0 1 0 0 0 0 0 - - - - 0 0 1 0 0 0 - - 0 1 0 1 0 1 1 1 3 ? 0 1 0 1 0 0 0 1 0 - 1 2 ? 1 0 0 ? 1 0 - - 0 ? ? ? 0 1 1 2 0 0 0 - 1 0 0 0 0 0 1 - ? ? ? ? ? 0 0 0 2 0 - - 1 0 2 0 - 2 0 1 0 1 2 0 1 1 1 1 1 1 - - 1 0 0 0 0 ? ? ? ? 2 2 1 0 1 ? ? ? ? ? ? ? ? ? ? ? ? ? ? ? ? ? ? ? ? ? ? ? ? ? ? ? ? ? ? ? ? ? ? ? ? ? ? ? ? ? ? ? ? ? ? ? ? ? ? ? ? ? ? ? ? ? ? ? ? ? ? ? ? ? ? ? ? ? ? ? ? ? ? ? ? ? ? ? ? ?

Cycnorhamphus_suevicus 1 - 0 0 0 - 1 0 0 2 1 0 1 2 - 1 1 - - 1 0 0 0 0 2 1 1 0 - 1 [01] 1 2 2 2 0 0 0 0 - - - - 0 0 1 1 0 0 - - 1 1 0 1 1 1 1 1 3 2 0 1 0 1 0 0 0 1 0 - 1 2 1 1 ? 0 ? 1 0 - - 0 1 0 1 0 1 0 0 2 0 0 - 1 0 0 1 0 0 1 - ? ? ? 0 - 0 1 1 1 1 - - 1 0 - - - 2 0 1 0 1 - - 1 0 0 0 1 1 - - 1 - - 0 0 0 0 0 0 0 0 0 0 1 ? 0 ? ? 1 ? ? 0 0 0 0 0 ? 0 2 ? 1 1 0 0 0 2 0 0 0 1 1 1 ? ? 1 0 8 0 0 0 0 0 0 ? ? 4 ? 0 1 1 0 0 0 0 0 3 0 1 1 3 0 1 0 0 0 0 0 1 0 1 1 0 0 1 3 3 0 1 0 0 0 1 0 0 1

Ardeadactylus_longicollum 1 - 1 0 1 0 0 0 0 0 1 0 1 2 - 1 1 - - 1 1 0 0 0 4 1 1 0 - 1 0 - - - - - - - 0 - - - - 0 0 1 ? 0 0 - - 1 1 0 0 - 1 1 0 3 2 0 1 0 1 0 0 0 1 0 1 1 2 1 ? 0 0 ? 1 0 - - 0 1 0 1 0 1 1 0 0 1 0 - 1 ? ? 0 0 0 1 - 1 0 0 1 0 1 0 1 2 0 - - 1 0 2 0 - 5 0 1 0 1 2 0 1 0 1 1 3 1 - - 1 0 0 0 0 0 0 1 0 2 2 1 0 1 1 0 0 0 1 ? ? 0 0 0 0 0 0 0 2 ? ? 0 1 0 0 1 1 0 ? 0 1 1 0 ? 1 0 5 0 0 0 0 0 0 ? ? 4 ? 0 1 1 0 0 ? 0 ? 3 0 1 1 3 0 1 0 0 1 1 0 1 0 1 1 0 0 1 ? ? 0 1 0 0 0 1 0 0 1

Pterodactylus_antiquus 1 - 1 0 0 - 0 0 0 0 1 0 1 1 - 1 1 - - 1 0 0 0 0 4 1 1 0 - 1 0 - - - - - - - 0 - - - - 0 0 1 1 0 0 - - 1 1 0 0 - 1 1 1 3 2 0 1 0 1 0 0 0 1 0 - 1 2 ? ? ? 0 ? 1 0 - - 0 ? ? ? ? 1 1 0 0 0 0 - 1 0 0 0 0 0 1 - 0 ? 0 0 - 0 0 1 3 0 - - 1 0 2 0 - 3 0 0 0 1 2 0 1 0 0 0 1 1 - - 1 0 0 0 0 0 0 0 ? 1 0 0 0 1 1 0 0 0 1 0 1 0 0 0 0 0 0 0 2 0 1 0 1 0 0 4 0 0 0 1 1 1 0 0 1 0 5 0 0 0 0 0 0 0 0 4 0 0 1 1 0 0 1 0 0 3 0 1 1 3 0 1 0 0 1 1 0 1 0 1 1 0 0 1 3 3 0 1 0 0 0 1 0 0 1

Lonchodectes_compressirostris ? ? 1 0 ? - ? 0 1 3 1 0 1 1 - ? ? ? ? ? ? 0 ? ? ? ? ? ? ? ? 0 - - - - - - - ? ? ? ? ? ? ? ? ? ? ? ? ? ? ? ? ? ? ? ? ? ? ? 0 ? ? ? ? ? ? ? ? ? ? ? ? ? ? ? ? 0 1 0 0 ? ? ? ? ? 1 1 ? 3 0 ? ? ? ? ? ? ? ? ? ? ? 0 0 ? ? ? ? ? ? 1 - - ? 0 0 0 - ? ? ? 0 ? 1 0 ? ? ? 0 3 ? ? ? ? 0 0 0 0 ? ? ? ? ? ? ? ? ? ? ? ? ? ? ? ? ? ? ? ? ? ? ? ? ? ? ? ? ? ? ? ? ? ? ? ? ? ? ? ? ? ? ? ? ? ? ? ? ? ? ? ? ? ? ? ? ? ? ? ? ? ? ? ? ? ? ? ? ? ? ? ? ? ? ? ? ? ? ? ? ? ? ? ? ? ? ? ? ? ? ?

Lonchodraco_giganteus 1 - 1 ? 0 - 0 0 ? 0 1 0 1 1 - ? ? - - 1 ? 0 ? ? ? ? ? 0 - ? 1 1 0 5 ? 0 1 1 ? - ? - - ? ? ? ? ? ? ? ? ? ? ? ? ? ? ? ? ? ? ? ? ? ? ? ? ? ? ? ? ? ? ? ? ? ? ? ? 1 0 0 0 ? ? ? ? 1 1 0 0 0 0 - 1 0 0 0 0 ? ? ? 1 ? 1 ? ? ? ? ? ? 2 0 0 ? 0 0 1 2 5 0 0 0 2 1 0 1 0 1 0 3 1 - - 1 0 0 0 0 ? ? ? ? ? ? ? ? ? ? ? ? ? ? ? ? ? ? 1 ? ? ? ? 2 ? ? ? ? ? ? ? ? 0 ? ? ? 1 ? ? 2 0 7 1 ? ? ? ? ? ? ? ? ? ? ? ? ? ? ? ? ? ? ? ? ? ? ? ? ? ? ? ? ? ? ? ? ? ? ? ? ? ? 0 0 0 1 0 ? 0 ? 0 1

Serradraco_sagittirostris 1 - 1 ? 0 - 0 0 ? 0 1 0 1 1 - ? ? - - 1 ? 0 ? ? ? ? ? 0 - ? 1 1 0 5 ? 0 1 1 ? - ? - - ? ? ? ? ? ? ? ? ? ? ? ? ? ? ? ? ? ? ? ? ? ? ? ? ? ? ? ? ? ? ? ? ? ? ? ? 1 0 0 0 ? ? ? ? 1 1 0 0 0 0 - 1 0 0 0 0 ? ? ? 1 ? 1 ? ? ? ? ? ? 2 0 0 ? 0 0 1 2 5 0 0 0 2 1 0 1 0 1 0 3 1 - - 1 0 0 0 0 ? ? ? ? ? ? ? ? ? ? ? ? ? ? ? ? ? ? 1 ? ? ? ? 2 ? ? ? ? ? ? ? ? 0 ? ? ? 1 ? ? 2 0 7 1 ? ? ? ? ? ? ? ? ? ? ? ? ? ? ? ? ? ? ? ? ? ? ? ? ? ? ? ? ? ? ? ? ? ? ? ? ? ? 0 0 0 1 0 ? 0 ? 0 1

Prejanopterus_curvirostris 1 - 1 0 0 - 0 0 0 0 1 0 1 1 - 1 1 - - 1 0 0 0 0 4 1 1 0 - 1 0 - - - - - - - 0 - - - - 0 0 1 1 0 0 - - 1 1 0 0 - 1 1 1 3 2 0 1 0 1 0 0 0 1 0 - 1 2 ? ? ? 0 ? 1 0 - - 0 ? ? ? ? 1 1 0 0 0 0 - 1 0 0 0 0 0 1 - 0 ? 0 0 - 0 0 1 3 0 - - 1 0 2 0 - 3 0 0 0 1 2 0 1 0 0 0 1 1 - - 1 0 0 0 0 0 0 0 ? 1 0 0 0 1 1 0 0 0 1 0 1 0 0 0 0 0 0 0 2 0 1 0 1 0 0 4 0 0 0 1 1 1 0 0 1 0 5 0 0 0 0 0 0 0 0 4 0 0 1 1 0 0 1 0 0 3 0 1 1 3 0 1 0 0 1 1 0 1 0 1 1 0 0 ? ? ? 0 0 0 1 0 0 0 0 0 1

Parirau_ataroa ? ? ? ? ? ? ? ? ? ? ? ? ? ? ? ? ? ? ? ? ? ? ? ? ? ? ? ? ? ? ? ? ? ? ? ? ? ? ? ? ? ? ? ? ? ? ? ? ? ? ? ? ? ? ? ? ? ? ? ? ? ? ? ? ? ? ? ? ? ? ? ? ? ? ? ? ? ? ? ? ? ? ? ? ? ? ? ? ? ? ? ? ? ? ? ? ? ? ? ? ? ? ? ? ? ? ? ? ? ? ? ? ? ? ? ? ? ? ? ? ? ? ? ? ? ? ? ? ? ? ? ? ? ? ? ? ? ? ? ? ? ? ? ? ? ? ? ? ? ? ? ? ? ? ? ? ? ? ? ? ? ? ? ? ? ? ? ? ? ? ? ? ? ? ? ? ? ? ? ? ? ? ? ? 0 0 ? ? ? ? ? ? ? ? ? ? ? ? ? ? ? ? ? ? ? ? ? ? ? ? ? ? ? ? ? ? ? ? ? ? 0 0 0 1 ? ? 0 ? ? ?

Yixianopterus_jingangshanensis 1 - 1 ? 0 - 0 0 ? 0 1 0 1 1 - ? ? - - 1 ? 0 ? ? ? ? ? 0 - ? 1 1 0 5 ? 0 1 1 ? - ? - - ? ? ? ? ? ? ? ? ? ? ? ? ? ? ? ? ? ? ? ? ? ? ? ? ? ? ? ? ? ? ? ? ? ? ? ? 1 0 0 0 ? ? ? ? 1 1 0 0 0 0 - 1 0 0 0 0 ? ? ? 1 ? 1 ? ? ? ? ? ? 2 0 0 ? 0 0 1 2 5 0 0 0 2 1 0 1 0 1 0 3 1 - - 1 0 0 0 0 ? ? ? ? ? ? ? ? ? ? ? ? ? ? ? ? ? ? 1 ? ? ? ? 2 ? ? ? ? ? ? ? ? 0 ? ? ? 1 ? ? 2 0 7 1 ? ? ? ? ? ? ? ? ? ? ? ? ? ? ? ? ? ? ? ? ? ? ? ? ? ? ? ? ? 1 0 1 1 0 0 ? ? ? 0 0 0 1 0 0 0 0 0 1

Navajodactylus_boerei ? ? ? ? ? ? ? ? ? ? ? ? ? ? ? ? ? ? ? ? ? ? ? ? ? ? ? ? ? ? ? ? ? ? ? ? ? ? ? ? ? ? ? ? ? ? ? ? ? ? ? ? ? ? ? ? ? ? ? ? ? ? ? ? ? ? ? ? ? ? ? ? ? ? ? ? ? ? ? ? ? ? ? ? ? ? ? ? ? ? ? ? ? ? ? ? ? ? ? ? ? ? ? ? ? ? ? ? ? ? ? ? ? ? ? ? ? ? ? ? ? ? ? ? ? ? ? ? ? ? ? ? ? ? ? ? ? ? ? ? ? ? ? ? ? ? ? ? ? ? ? ? ? ? ? ? ? ? ? ? ? ? ? ? ? ? ? ? ? ? ? ? ? ? ? ? ? ? ? ? ? ? ? ? 0 0 ? ? ? ? ? ? ? ? ? ? ? ? ? ? ? ? ? ? ? ? ? ? ? ? ? ? ? ? ? ? ? ? ? ? 0 0 0 1 ? ? 0 ? ? ?

Barbaridactylus_grandis ? ? ? ? ? ? ? ? ? ? ? ? ? ? ? ? ? ? ? ? ? ? ? ? ? ? ? ? ? ? ? ? ? ? ? ? ? ? ? ? ? ? ? ? ? ? ? ? ? ? ? ? ? ? ? ? ? ? ? ? ? ? ? ? ? ? ? ? ? ? ? ? ? ? ? ? ? ? ? ? ? ? ? ? ? ? ? ? ? ? ? ? ? ? ? ? ? ? ? 1 1 - ? ? ? ? ? ? ? 1 0 ? ? ? ? ? ? ? ? ? ? ? ? ? ? ? ? ? ? ? ? ? ? ? ? ? ? ? ? ? ? ? ? ? ? 1 0 1 ? ? ? ? ? ? ? ? ? ? ? ? ? ? 2 ? ? ? ? ? ? ? 1 0 0 0 1 ? ? ? 3 1 6 0 1 0 0 0 ? ? ? ? ? ? ? ? ? ? ? ? ? ? ? ? ? ? ? ? ? ? ? ? ? 1 1 1 1 ? 1 ? ? ? 1 0 0 0 0 0 1 1 1

Alcione_elainus ? ? ? ? ? ? ? ? ? ? ? ? ? ? ? ? ? ? ? ? ? ? ? ? ? ? ? ? ? ? ? ? ? ? ? ? ? ? ? ? ? ? ? ? ? ? ? ? ? ? ? ? ? ? ? ? ? ? ? ? ? ? ? ? ? ? ? ? ? ? ? ? ? ? ? ? ? ? ? ? ? ? ? ? ? ? ? ? ? ? ? ? ? ? ? ? ? ? ? ? ? ? ? ? ? ? ? ? ? ? ? ? ? ? ? ? ? ? ? ? ? ? ? ? ? ? ? ? ? ? ? ? ? ? ? ? ? ? ? ? ? ? ? ? ? ? ? ? ? ? ? ? ? ? ? ? ? 1 1 0 ? 0 2 ? 0 1 1 1 0 ? 0 0 0 0 0 0 0 0 3 1 6 0 1 0 0 0 ? ? ? ? ? ? ? 1 0 0 ? 1 0 ? ? ? ? ? ? ? ? ? ? ? ? 1 1 1 1 0 1 ? ? ? 1 0 0 0 0 0 1 1 1

Simurghia_robusta ? ? ? ? ? ? ? ? ? ? ? ? ? ? ? ? ? ? ? ? ? ? ? ? ? ? ? ? ? ? ? ? ? ? ? ? ? ? ? ? ? ? ? ? ? ? ? ? ? ? ? ? ? ? ? ? ? ? ? ? ? ? ? ? ? ? ? ? ? ? ? ? ? ? ? ? ? ? ? ? ? ? ? ? ? ? ? ? ? ? ? ? ? ? ? ? ? ? ? ? ? ? ? ? ? ? ? ? ? ? ? ? ? ? ? ? ? ? ? ? ? ? ? ? ? ? ? ? ? ? ? ? ? ? ? ? ? ? ? ? ? ? ? ? ? ? ? ? ? ? ? ? ? ? ? ? ? ? ? ? ? ? ? ? ? ? ? ? ? ? ? ? ? 0 0 ? ? ? ? ? 6 0 ? 0 ? ? ? ? ? ? ? ? ? ? ? ? ? ? ? ? ? ? ? ? ? ? ? ? ? ? ? ? ? ? ? ? ? ? ? ? ? ? ? ? ? ? 0 ? 1 1

Nyctosaurus_gracilis 2 - 1 0 0 - 1 0 1 0 1 0 0 1 - - 0 - - 1 1 0 0 0 2 1 1 0 - 1 0 - - - - - - - 0 - - - - 1 - - - - 1 2 0 1 1 0 1 1 0 0 1 2 3 0 1 1 1 0 0 0 1 0 - 1 1 1 1 0 0 ? 0 0 - - 0 1 ? 1 1 1 1 1 0 0 0 - 1 1 0 1 0 1 1 - 0 0 0 1 0 0 1 1 0 1 - - 1 1 - - - - - - - - - - - - - - - - - - - - - - 0 1 0 0 1 0 3 1 0 1 0 1 1 1 1 1 1 0 1 1 1 0 0 0 2 0 0 1 1 1 0 1 1 0 0 0 1 1 0 0 3 1 6 0 1 0 0 0 0 1 1 1 ? 2 1 1 0 0 ? 1 0 2 0 1 1 2 0 1 1 0 1 1 0 1 1 1 1 0 1 1 ? ? 1 0 0 0 0 0 1 1 1

Nyctosaurus_nanus ? ? ? ? ? ? ? ? ? ? ? ? ? ? ? ? ? ? ? ? ? ? ? ? ? ? ? ? ? ? ? ? ? ? ? ? ? ? ? ? ? ? ? ? ? ? ? ? ? ? ? ? ? ? ? ? ? ? ? ? ? ? ? ? ? ? ? ? ? ? ? ? ? ? ? ? ? ? ? ? ? ? ? ? ? ? ? 1 ? ? ? ? ? ? ? ? ? ? ? 1 1 - ? ? ? ? ? 0 ? 1 0 ? ? ? 1 ? ? ? ? ? ? ? ? ? ? ? ? ? ? ? ? ? ? ? ? ? ? ? ? ? ? ? ? ? ? ? ? ? ? ? 1 ? ? ? ? ? ? ? ? ? ? 0 2 ? ? ? ? 1 0 ? 1 0 0 0 1 ? 0 0 ? 1 6 0 1 0 ? ? ? ? ? ? ? ? ? ? ? ? ? ? ? ? ? ? ? ? ? ? ? ? ? ? ? ? ? ? ? ? ? ? ? ? 1 0 0 0 0 0 1 1 1

Nyctosaurus_lamegoi ? ? ? ? ? ? ? ? ? ? ? ? ? ? ? ? ? ? ? ? ? ? ? ? ? ? ? ? ? ? ? ? ? ? ? ? ? ? ? ? ? ? ? ? ? ? ? ? ? ? ? ? ? ? ? ? ? ? ? ? ? ? ? ? ? ? ? ? ? ? ? ? ? ? ? ? ? ? ? ? ? ? ? ? ? ? ? ? ? ? ? ? ? ? ? ? ? ? ? ? ? ? ? ? ? ? ? ? ? ? ? ? ? ? ? ? ? ? ? ? ? ? ? ? ? ? ? ? ? ? ? ? ? ? ? ? ? ? ? ? ? ? ? ? ? ? ? ? ? ? ? ? ? ? ? ? ? ? ? ? ? ? ? ? ? ? ? ? ? ? ? ? ? 0 1 ? ? ? ? 1 6 0 ? ? ? ? ? ? ? ? ? ? ? ? ? ? ? ? ? ? ? ? ? ? ? ? ? ? ? ? ? ? ? ? ? ? ? ? ? ? ? ? ? ? ? ? 1 ? ?

Muzquizopteryx_coahuilensis ? ? ? ? ? ? ? ? 1 ? 1 ? ? ? - - ? - - 1 1 0 0 0 2 1 1 0 - 1 0 - - - - - - - 0 - - - - 1 - - - 0 0 - - 1 1 0 1 1 0 0 1 2 2 0 1 ? 1 0 0 0 1 0 - 1 1 ? ? ? 0 ? ? ? ? ? ? ? ? ? ? 1 ? ? ? ? ? ? ? ? ? ? ? 1 1 - ? ? ? ? ? 0 1 0 3 ? ? ? 1 1 - - - - - - - - - - - - - - - - - - - - - - 0 1 ? ? ? 0 3 ? 0 1 ? ? 1 1 1 ? 1 0 ? 1 1 0 0 0 ? ? 0 ? 1 ? ? 1 ? 0 ? 0 0 1 ? ? ? 1 6 0 1 0 ? ? ? ? ? 1 0 2 ? 1 ? ? ? ? ? ? ? ? 1 2 0 ? 1 0 1 ? ? 1 ? 1 1 ? 1 1 4 ? 1 0 0 0 0 0 1 1 1

Haopterus_gracilis 1 - 1 0 0 - 0 0 0 0 1 0 1 1 - 1 ? - - 1 ? 0 0 0 ? ? 1 0 - ? 0 - - - - - - - 0 - - - - 0 1 0 0 0 0 - - ? 0 0 0 - 0 ? 1 2 3 0 ? ? 1 ? ? ? ? 0 - 1 ? ? ? ? 0 ? 0 ? ? ? ? ? ? ? ? 1 1 2 0 0 0 - 1 ? 0 ? 0 ? 1 - ? ? ? ? ? 1 0 1 0 0 - - ? 0 2 0 - 3 0 0 0 2 2 0 1 0 0 0 1 1 - - 1 0 0 0 0 ? ? ? ? ? ? 0 0 1 ? 0 0 ? ? ? ? ? ? 1 0 0 0 0 2 0 1 0 1 ? ? 2 0 0 0 0 1 1 ? ? ? 0 7 1 0 0 ? ? ? ? ? 4 ? 0 1 ? 0 0 1 0 0 ? ? ? ? ? ? ? ? ? ? ? ? ? ? ? ? ? ? 1 - - 0 0 1 0 0 0 0 1 0 0

Mimodactylus_libanensis 1 - 1 0 0 - 0 0 0 0 1 0 1 1 - 1 0 - - 1 0 0 0 0 1 1 1 0 - ? 0 - - - - - - - 0 - - - - 0 1 0 0 0 0 - - ? 0 0 0 - 0 ? 1 2 3 0 ? ? 1 ? ? ? ? 0 - 1 ? ? ? ? 0 ? 0 ? ? ? ? ? ? ? ? 1 1 2 0 0 0 - 1 ? 0 ? 0 ? 1 - ? ? ? ? ? 1 0 1 0 0 - - ? 0 2 0 - 3 0 0 0 2 2 0 1 0 0 0 1 1 - - 1 0 0 0 1 1 0 1 ? 0 1 1 0 1 ? ? 1 ? ? ? ? ? ? ? 1 1 1 0 2 1 1 1 0 1 0 2 0 0 0 0 0 1 0 1 2 0 7 1 1 0 1 0 ? 1 ? 4 ? 1 ? 1 0 0 1 1 ? 3 ? ? ? ? ? ? 0 0 1 1 0 1 ? 1 1 0 1 1 1 - - 0 0 1 0 0 0 0 1 0 0

Hongshanopterus_lacustris 2 - 1 0 0 - 1 0 1 0 ? 0 1 ? - ? ? - - 1 1 0 ? 0 ? 1 1 0 - ? ? ? ? ? ? ? ? ? 0 - - - ? 0 1 0 0 0 ? ? ? ? ? ? ? ? 0 ? 1 ? [23] 0 ? ? 1 ? 0 0 1 ? - 1 1 1 1 0 ? ? 0 1 0 0 0 1 ? 1 1 1 ? ? ? ? ? ? ? ? ? ? ? ? ? ? ? ? ? ? ? ? ? ? ? ? ? ? ? 0 2 0 - 0 0 0 0 2 2 0 ? 0 1 0 3 1 - - 1 0 0 - 0 1 0 1 ? 0 1 1 0 1 ? ? ? ? ? ? ? ? ? ? ? ? ? ? ? ? ? ? ? ? ? ? ? ? ? ? ? ? ? ? ? ? ? ? ? ? ? ? ? ? ? ? ? ? ? ? ? ? ? ? ? ? ? ? ? ? ? ? ? ? ? ? ? ? ? ? ? ? ? ? ? ? ? ? ? ? ? ? ? ? ? ?

Nurhachius_ignaciobritoi 1 - 1 ? 0 - 0 0 0 0 1 0 1 1 - 1 0 - - 1 0 0 1 0 ? 1 1 0 - 1 0 - - - - - - - 0 - - - - 0 1 0 0 0 0 - - 1 1 1 0 - 0 0 1 3 3 0 1 ? 1 0 0 1 1 0 - 1 1 ? ? ? 0 ? ? 1 1 1 0 ? 0 ? 0 1 1 2 0 0 0 - 1 0 0 0 0 0 1 - 1 0 0 0 - 1 0 1 [02] 0 - - 1 0 2 0 - 4 0 0 0 1 1 0 1 0 1 0 1 1 - - 1 0 0 0 1 1 0 1 ? 0 1 1 0 1 ? ? 1 ? ? ? ? ? ? ? 1 1 ? 0 2 1 ? 1 0 1 0 2 0 0 ? 0 0 1 0 ? ? 0 7 1 1 0 1 0 ? 1 ? 4 ? 1 ? 1 0 0 1 1 ? 3 ? ? ? ? ? ? 0 0 1 1 0 1 ? 1 ? 0 1 ? 4 – 0 0 1 0 0 0 0 1 0 0

Liaoxipterus_brachyognathus ? ? ? ? ? - ? ? ? ? ? 1 1 ? ? ? ? ? ? ? ? ? ? ? ? ? ? ? ? ? ? ? ? ? ? ? ? ? ? ? ? ? ? ? ? ? ? ? ? ? ? ? ? ? ? ? ? ? ? ? ? ? ? ? ? ? ? ? ? ? ? ? ? ? ? ? ? ? ? ? ? ? ? ? ? ? ? ? 1 ? 2 0 0 - 1 0 ? ? 0 ? 1 - ? ? ? 0 ? 1 0 1 2 ? ? ? ? 0 2 0 - 4 0 0 0 0 0 0 ? 0 1 0 1 1 - - ? ? 0 0 ? ? ? ? ? ? ? ? ? ? ? ? ? ? ? ? ? ? ? ? ? ? ? ? ? ? ? ? ? ? ? ? ? ? ? ? ? ? ? ? ? ? ? ? ? ? ? ? ? ? ? ? ? ? ? ? ? ? ? ? ? ? ? ? ? ? ? ? ? ? ? ? ? ? ? ? ? ? ? ? ? ? ? ? ? ? ? ? ? ? ? ?

Istiodactylus_sinensis 1 - 1 ? 0 - 0 0 0 2 1 1 ? 1 - 1 0 - - 1 0 0 0 1 2 1 1 0 - 1 0 - - - - - - - 0 - - - - 1 - - - - 0 - - 1 ? 1 0 - 0 0 1 3 4 0 1 1 1 0 0 1 2 1 - 1 1 ? ? ? 0 ? ? 1 1 1 0 ? 0 ? 0 1 1 2 ? 0 0 - 1 0 0 0 0 0 1 - ? ? ? ? ? 1 0 1 2 0 - - 1 0 2 0 - 4 3 0 0 0 0 0 1 0 1 0 1 1 - - 1 0 0 0 1 1 ? ? ? 0 1 ? 0 1 ? ? 1 ? ? ? ? ? ? ? ? 1 ? 0 2 ? ? ? ? 0 0 ? ? 0 0 0 0 1 ? ? ? 0 7 1 ? ? ? 0 0 ? ? ? 0 1 ? ? 0 0 ? ? 0 3 1 1 ? ? ? ? ? ? ? ? ? 1 ? 1 ? 0 ? ? ? ? 0 0 1 0 0 0 0 1 0 0

Istiodactylus_latidens 1 - 1 ? 0 - 0 0 0 2 1 1 1 1 - 1 0 - - 1 0 0 0 1 2 1 1 0 - ? 0 - - - - - - - 0 - - - - 1 - - - - 0 - - 1 1 1 0 ? 0 0 1 3 4 0 1 ? 1 0 0 1 2 1 - 1 1 ? ? ? ? 1 0 ? ? ? ? ? 0 ? 0 1 1 2 2 0 0 - 1 0 0 0 0 0 1 - ? ? ? 0 - 1 0 1 2 0 - - 1 0 2 0 - 4 3 0 0 0 0 0 1 0 1 0 1 1 - - 1 0 0 0 1 ? ? ? ? 0 ? 1 0 ? ? 1 1 1 ? ? ? ? ? 1 1 1 1 0 2 1 0 1 0 1 0 ? 0 0 ? 0 0 1 0 1 2 0 7 1 1 0 1 0 0 1 1 4 ? ? ? 1 0 0 ? 1 ? 3 1 1 ? ? ? ? ? ? ? ? ? 1 1 1 1 0 1 ? ? ? 0 0 1 0 0 0 0 1 0 0

Gwawinapterus_beardi 1 - 1 ? 0 - 0 0 0 2 1 1 1 1 - 1 0 - - 1 0 0 0 1 2 1 1 0 - ? 0 - - - - - - - 0 - - - - 1 - - - - 0 - - 1 1 1 0 ? 0 0 1 3 4 0 1 ? 1 0 0 1 2 1 - 1 1 ? ? ? ? ? 0 ? ? ? ? ? 0 ? 0 1 1 2 2 0 0 - ? ? ? ? ? ? ? - ? ? ? ? - ? ? ? ? ? - - ? ? ? ? - ? ? ? ? ? ? ? ? ? ? ? ? ? - - ? ? ? ? ? ? ? ? ? ? ? ? ? ? ? ? ? ? ? ? ? ? ? ? ? ? ? ? ? ? ? ? ? ? ? ? ? ? ? ? ? ? ? ? ? ? ? ? ? ? ? ? ? ? ? ? ? ? ? ? ? ? ? ? ? ? ? ? ? ? ? ? ? ? ? ? ? ? ? ? ? ? ? ? ? ? ? ? ? ? ? ? ? ?

Aetodactylus_halli ? - ? ? ? 0 0 0 ? 2 ? 1 0 ? ? ? ? ? ? ? ? ? ? ? ? ? ? ? ? ? 0 - - - - - - - ? ? ? ? ? ? ? ? ? ? ? ? ? ? ? ? ? ? ? ? ? ? ? ? ? ? ? ? ? ? ? ? ? ? ? ? ? ? ? ? ? 1 0 0 ? ? ? ? 1 1 1 0 2 1 0 - 1 0 2 0 0 0 1 - 1 0 0 0 - 1 ? 1 0 1 - - 1 0 1 0 - 5 1 ? 0 1 2 0 1 0 1 1 3 1 - - ? ? 0 0 1 ? ? ? ? ? ? ? ? ? ? ? ? ? ? ? ? ? ? ? ? ? ? ? ? ? ? ? ? ? ? ? ? ? ? ? ? ? ? ? ? ? ? ? ? ? ? ? ? ? ? ? ? ? ? ? ? ? ? ? ? ? ? ? ? ? ? ? ? ? ? ? ? ? ? ? ? ? ? ? ? ? ? ? ? ? ? ? ? ? ? ?

Cimoliopterus_cuvieri 1 - 1 0 1 0 0 0 ? 0 1 1 ? 1 ? ? ? ? ? ? ? ? ? ? ? ? ? ? ? ? 1 1 0 4 ? 0 1 1 ? ? ? ? ? ? ? ? ? ? ? ? ? ? ? ? ? ? ? ? ? ? ? ? ? ? ? ? ? ? ? ? ? ? ? ? ? ? ? ? ? 1 0 0 ? ? ? ? ? 1 ? ? ? ? ? ? ? ? ? ? ? ? ? ? ? ? ? ? ? ? ? ? ? ? ? ? ? 0 2 0 - 5 1 0 0 1 2 0 ? 0 1 1 3 1 - - 1 2 0 ? 1 ? ? ? ? ? ? ? ? ? ? ? ? ? ? ? ? ? ? ? ? ? ? ? ? ? ? ? ? ? ? ? ? ? ? ? ? ? ? ? ? ? ? ? ? ? ? ? ? ? ? ? ? ? ? ? ? ? ? ? ? ? ? ? ? ? ? ? ? ? ? ? ? ? ? ? ? ? ? ? ? ? ? ? ? ? ? ? ? ? ? ?

Cimoliopterus_dunni 1 - 1 0 1 0 0 0 ? 0 1 1 ? 1 ? ? ? ? ? ? ? ? ? ? ? ? ? ? ? ? 1 1 0 4 ? 0 1 1 ? ? ? ? ? ? ? ? ? ? ? ? ? ? ? ? ? ? ? ? ? ? ? ? ? ? ? ? ? ? ? ? ? ? ? ? ? ? ? ? ? 1 0 0 ? ? ? ? ? 1 ? ? ? ? ? ? ? ? ? ? ? ? ? ? ? ? ? ? ? ? ? ? ? ? ? ? ? 0 2 0 - 5 ? ? 0 1 2 0 ? 0 1 1 3 1 - - 1 2 0 ? 1 ? ? ? ? ? ? ? ? ? ? ? ? ? ? ? ? ? ? ? ? ? ? ? ? ? ? ? ? ? ? ? ? ? ? ? ? ? ? ? ? ? ? ? ? ? ? ? ? ? ? ? ? ? ? ? ? ? ? ? ? ? ? ? ? ? ? ? ? ? ? ? ? ? ? ? ? ? ? ? ? ? ? ? ? ? ? ? ? ? ? ?

Boreopterus_cuiae 1 - 1 ? 0 - 1 0 1 0 1 ? ? 1 - 1 ? - - 1 1 0 1 0 ? 1 1 0 - 1 0 - - - - - - - 0 - - - - ? ? ? ? ? 1 0 2 1 1 ? 1 0 0 0 1 2 2 0 1 ? 1 0 0 0 1 0 - 1 1 ? ? ? ? ? ? ? ? ? ? ? ? ? ? 1 1 2 0 0 0 - 1 0 0 1 0 1 1 - ? ? ? ? ? 0 0 1 0 [01] - - 1 0 0 0 - 2 0 1 0 2 2 0 1 0 0 1 1 1 - - 1 0 0 0 0 ? ? ? ? 0 1 1 0 1 ? ? ? ? ? ? 1 0 ? ? ? ? ? 0 ? ? ? ? ? ? ? ? ? ? 1 0 ? ? ? ? ? ? ? ? ? ? ? ? ? ? ? ? ? 1 1 ? ? ? 1 ? 0 ? ? ? ? ? ? ? ? ? ? ? ? 1 ? 1 1 ? 1 1 4 - 1 0 0 0 0 0 0 1 1 0

Guidraco_venator 1 - 1 ? 1 0 0 0 1 0 1 ? ? 1 - 1 0 - - 1 1 0 1 0 2 1 1 0 - 1 0 - - - - - - - 0 - - - - ? ? ? ? ? 1 2 1 1 1 1 1 2 0 0 1 2 2 0 1 1 1 0 0 0 0 0 - 1 1 ? ? ? 0 ? ? ? ? ? 0 ? 0 ? 1 1 1 ? 0 1 0 - 1 0 0 0 0 1 1 - ? ? ? ? ? 1 0 1 0 0 - - 1 0 2 1 2 5 1 1 0 1 2 1 1 0 1 1 1 1 - - 1 0 0 0 1 ? 0 1 ? ? ? 1 0 1 ? ? ? ? ? ? ? ? ? ? ? ? ? ? ? ? ? ? ? ? ? ? ? ? ? ? ? ? ? ? ? ? ? ? ? ? ? ? ? ? ? ? ? ? ? ? ? ? ? ? ? ? ? ? ? ? ? ? ? ? ? ? ? ? ? ? ? ? ? ? ? ? ? ? ? ? ? ? ? ? ? ?

Zhenyuanopterus_longirostris 1 - 1 ? 0 - 0 0 1 0 1 ? ? 1 - 1 1 - - 1 1 0 1 0 4 1 1 0 - 1 1 1 2 3 1 0 ? ? 0 - - - - 0 1 1 0 0 1 0 2 1 1 0 1 0 0 0 1 2 2 0 1 0 1 0 0 0 1 0 - 1 1 ? ? ? 0 ? ? ? ? ? ? ? 0 ? ? 1 1 ? 0 0 0 - 1 0 0 0 0 1 1 - ? ? ? ? ? 0 0 1 0 [01] - - 1 0 0 0 - 2 0 1 0 2 2 0 1 0 0 0 1 1 - - 1 0 0 0 0 ? ? 1 ? 0 1 1 0 1 ? 1 1 1 1 ? 1 0 ? 1 1 1 1 0 2 ? ? ? ? ? ? 2 ? 0 ? 0 0 1 ? ? 2 0 7 1 1 0 ? ? ? ? ? 4 ? ? 1 1 0 ? 1 ? ? ? ? ? ? ? 0 ? ? ? ? ? ? 1 ? 1 1 ? 1 1 4 - 1 0 0 0 0 0 0 1 1 0

Anhanguera_santanae 1 - 1 0 1 0 0 0 1 0 1 0 0 1 - 1 0 - - 1 1 0 1 0 2 1 1 0 - 1 1 1 0 4 0 ? 1 1 0 - - - - 0 1 1 0 1 1 0 2 1 1 1 1 0 0 0 1 2 3 0 1 1 1 0 1 0 1 0 - 1 1 1 1 0 0 1 0 1 0 0 0 1 1 1 1 1 1 0 0 ? 0 - 1 0 0 0 0 0 1 - 1 0 1 0 - 0 0 1 0 2 ? 1 1 0 2 1 2 5 1 ? 0 1 2 1 1 1 1 1 3 1 - - 1 0 1 0 1 1 0 1 1 0 1 1 0 1 0 1 0 0 1 0 1 0 ? 1 1 1 0 0 2 ? ? ? ? 1 1 ? 0 0 1 0 0 ? ? ? 2 0 7 1 1 0 0 0 0 1 1 4 0 1 ? 1 ? ? ? ? 0 2 1 1 ? ? 1 1 1 0 1 0 0 ? 0 1 1 ? ? ? ? ? 1 0 0 0 ? ? 0 1 1 0

Anhanguera_piscator 1 - 1 ? 1 0 0 0 1 0 1 0 0 1 - 1 0 - - 1 1 0 1 0 2 1 1 0 - 1 1 1 0 4 0 0 1 1 0 - - - - 0 1 1 0 1 1 0 2 1 1 1 1 0 0 0 1 2 3 0 1 1 1 0 1 0 1 0 - 1 1 1 1 0 0 1 0 ? ? ? ? 1 ? ? 1 1 1 0 0 1 0 - 1 0 0 0 0 0 1 - ? ? ? 0 - 0 0 1 0 2 0 1 1 0 2 1 2 5 1 1 0 1 2 1 1 1 1 1 3 1 - - 1 0 1 0 1 1 0 1 1 0 1 1 0 1 0 1 0 0 1 0 1 0 1 ? 1 1 0 0 2 0 0 1 0 1 1 2 0 0 1 0 0 1 0 1 2 0 7 1 1 0 0 ? 0 1 1 4 ? 1 ? 1 0 1 1 1 0 2 1 1 ? ? ? 1 1 0 1 1 0 1 0 1 1 0 1 ? 4 - 1 0 0 0 0 0 0 1 1 0

Anhanguera_blittersdorffi 1 - 1 0 1 0 0 0 1 0 1 0 0 1 - 1 0 - - 1 1 0 1 0 2 1 1 0 - 1 1 1 0 4 0 0 1 1 0 - - - - 1 - - - - 1 0 2 1 1 1 1 0 0 0 1 2 3 0 1 1 1 0 1 0 1 0 - 1 1 1 1 0 0 1 0 1 0 0 0 1 1 1 1 1 1 0 0 1 0 - 1 0 0 ? 0 ? 1 - 1 0 ? ? ? ? 0 1 ? 2 0 1 ? 0 2 1 2 5 1 1 0 1 2 1 1 1 1 1 3 1 - - 1 0 1 0 1 ? ? ? ? ? ? ? ? ? ? ? ? ? ? ? ? ? ? ? ? ? ? ? ? ? ? ? ? ? ? ? ? ? ? ? ? ? ? ? ? ? ? ? ? ? ? ? ? ? ? ? ? ? ? ? ? ? ? ? ? ? ? ? ? ? ? ? ? ? ? ? ? ? ? ? ? ? ? ? ? ? ? ? ? ? ? ? ? ? ? ?

Anhanguera_araripensis 1 - 1 0 1 0 ? 0 1 0 1 0 0 1 - 1 0 - - 1 1 0 1 0 2 1 1 0 - 1 1 1 0 4 0 0 1 1 0 - - - - ? - - - - 1 0 2 1 1 1 1 0 0 0 1 2 3 0 1 1 1 0 1 0 1 0 - 1 1 1 1 0 0 1 0 1 0 0 0 1 1 1 1 1 ? ? 0 ? ? ? 1 0 ? ? 0 0 1 - ? ? ? 0 - 0 0 1 0 ? ? ? 1 0 2 1 2 5 ? ? 0 1 2 1 1 1 1 1 3 1 - - 1 0 1 ? 1 ? ? ? ? ? ? ? ? ? ? ? ? ? ? ? ? ? ? ? ? ? ? ? ? ? ? ? ? ? ? ? 0 0 1 0 0 1 0 1 2 0 ? 1 1 0 ? 0 0 1 1 4 ? 1 ? 1 ? ? ? ? ? ? ? ? ? ? ? ? ? ? ? ? ? ? ? ? ? ? ? ? ? ? 1 0 0 0 0 0 0 1 1 0

Liaoningopterus_gui 1 - 1 0 1 0 0 0 1 0 1 0 ? 1 - 1 0 - - 1 1 0 1 [02] 2 1 1 0 - ? 1 1 0 4 0 0 1 1 0 - - - - ? ? ? ? ? ? ? ? ? ? 1 ? ? 0 0 1 2 3 0 1 ? 1 0 1 0 1 0 - 1 ? ? ? ? ? ? ? 1 0 0 ? ? ? ? 1 1 1 0 0 ? 0 - ? 0 0 ? 0 0 1 - ? ? ? ? ? 0 0 1 0 2 ? ? ? 0 2 1 2 5 1 1 0 1 2 1 1 0 1 1 3 1 - - 1 0 1 0 1 ? 0 1 ? ? ? 1 0 1 ? ? ? ? ? ? ? ? ? ? ? ? ? ? ? ? ? ? ? ? ? ? ? ? ? ? ? ? ? ? ? ? ? ? ? ? ? ? ? ? ? ? ? ? ? ? ? ? ? ? ? ? ? ? ? ? ? ? ? ? ? ? ? ? ? ? ? ? ? ? ? ? 1 0 0 0 0 0 0 1 1 0

Siroccopteryx_moroccensis 0 - 1 0 1 1 0 0 ? 0 ? 0 0 ? ? ? ? ? ? ? ? ? ? ? ? ? ? ? ? ? 1 0 0 4 ? 0 1 ? ? ? ? ? ? ? ? ? ? ? ? ? ? ? ? ? ? ? ? ? ? ? ? ? ? ? ? ? ? ? ? ? ? ? ? ? ? ? ? ? ? 1 0 1 ? ? ? ? ? 1 ? ? ? ? ? ? ? ? ? ? ? ? ? ? ? ? ? ? ? ? ? ? ? ? ? ? ? 0 2 ? 2 5 ? 0 0 1 2 ? ? 0 1 1 3 1 - - 2 ? 0 ? ? ? ? ? ? ? ? ? ? ? ? ? ? ? ? ? ? ? ? ? ? ? ? ? ? ? ? ? ? ? ? ? ? ? ? ? ? ? ? ? ? ? ? ? ? ? ? ? ? ? ? ? ? ? ? ? ? ? ? ? ? ? ? ? ? ? ? ? ? ? ? ? ? ? ? ? ? ? ? ? ? ? ? ? ? ? ? ? ? ? ? ?

Tropeognathus_mesembrinus 1 - 1 0 1 0 0 0 1 0 1 0 0 1 - 1 0 - - 1 1 0 1 0 2 1 1 0 - ? 1 0 0 4 0 0 1 1 0 - - - - 1 - - - - 1 0 2 1 1 1 1 0 0 0 1 2 3 0 1 1 1 0 1 0 1 0 - 1 1 1 1 0 0 1 0 1 0 0 0 1 1 1 1 1 1 0 0 1 0 - 1 0 0 0 0 0 1 - 1 0 0 0 - 0 0 1 3 2 0 1 ? 0 2 1 2 5 1 ? 0 1 2 1 1 0 1 1 3 1 - - 1 0 0 0 1 ? ? ? ? ? ? ? ? ? ? ? ? ? ? ? ? ? ? ? ? ? ? ? ? ? ? ? ? ? ? ? ? ? ? ? ? ? ? ? ? ? ? ? ? ? ? ? ? ? ? ? ? ? ? ? ? ? ? ? ? ? ? ? ? ? ? ? ? ? ? ? ? ? ? ? ? ? ? ? ? ? 1 0 0 0 0 0 0 1 1 0

Coloborhynchus_clavirostris 0 - 1 1 1 2 0 ? ? 0 ? ? ? ? ? ? ? ? ? ? ? ? ? ? ? ? ? ? ? ? 1 0 0 4 ? 0 1 1 ? ? ? ? ? ? ? ? ? ? ? ? ? ? ? ? ? ? ? ? ? ? ? ? ? ? ? ? ? ? ? ? ? ? ? ? ? ? ? ? ? 1 0 1 ? ? ? ? ? 1 ? ? ? ? ? ? ? ? ? ? ? ? ? ? ? ? ? ? ? ? ? ? ? ? ? ? ? 0 2 ? 2 5 ? ? 0 1 2 1 ? ? ? 1 3 ? ? ? 2 ? 0 ? ? ? ? ? ? ? ? ? ? ? ? ? ? ? ? ? ? ? ? ? ? ? ? ? ? ? ? ? ? ? ? ? ? ? ? ? ? ? ? ? ? ? ? ? ? ? ? ? ? ? ? ? ? ? ? ? ? ? ? ? ? ? ? ? ? ? ? ? ? ? ? ? ? ? ? ? ? ? ? ? ? ? ? ? ? ? ? ? ? ? ? ?

Coloborhynchus_wadleighi 0 - 1 1 1 2 0 ? ? 0 1 0 0 ? ? ? ? ? ? ? ? ? ? ? ? ? ? ? ? ? 1 0 0 4 ? 0 1 1 ? ? ? ? ? ? ? ? ? ? ? ? ? ? ? ? ? ? ? ? ? ? ? ? ? ? ? ? ? ? ? ? ? ? ? ? ? ? ? ? ? 1 0 1 ? ? ? ? ? 1 ? ? ? ? ? ? ? ? ? ? ? ? ? ? ? ? ? ? ? ? ? ? ? ? ? ? ? 0 2 ? 2 5 ? ? 0 1 2 1 ? ? 1 1 3 1 - - 2 ? 0 ? ? ? ? ? ? ? ? ? ? ? ? ? ? ? ? ? ? ? ? ? ? ? ? ? ? ? ? ? ? ? ? ? ? ? ? ? ? ? ? ? ? ? ? ? ? ? ? ? ? ? ? ? ? ? ? ? ? ? ? ? ? ? ? ? ? ? ? ? ? ? ? ? ? ? ? ? ? ? ? ? ? ? ? ? ? ? ? ? ? ? ? ?

Ornithocheirus_simus 0 - 1 0 1 0 0 ? ? 0 ? ? ? ? ? ? ? ? ? ? ? ? ? ? ? ? ? ? ? ? 1 0 0 4 ? 0 1 1 ? ? ? ? ? ? ? ? ? ? ? ? ? ? ? ? ? ? ? ? ? ? ? ? ? ? ? ? ? ? ? ? ? ? ? ? ? ? ? ? ? 1 0 1 ? ? ? ? ? 1 ? ? ? ? ? ? ? ? ? ? ? ? ? ? ? ? ? ? ? ? ? ? ? ? ? ? ? 0 2 ? 2 5 ? ? 0 1 ? ? ? ? ? 0 3 ? ? ? 1 ? ? ? ? ? ? ? ? ? ? ? ? ? ? ? ? ? ? ? ? ? ? ? ? ? ? ? ? ? ? ? ? ? ? ? ? ? ? ? ? ? ? ? ? ? ? ? ? ? ? ? ? ? ? ? ? ? ? ? ? ? ? ? ? ? ? ? ? ? ? ? ? ? ? ? ? ? ? ? ? ? ? ? ? ? 1 ? ? ? ? ? ? ? ? ?

Brasileodactylus_araripensis ? ? ? ? ? 0 ? ? ? ? ? 0 0 ? ? ? ? ? ? ? ? ? ? ? ? ? ? ? ? ? ? ? ? ? ? ? ? ? ? ? ? ? ? ? ? ? ? ? ? ? ? ? ? ? ? ? ? ? ? ? ? ? ? ? ? ? ? ? ? ? ? ? ? ? ? ? ? ? ? ? ? ? ? ? ? ? ? 1 1 0 0 1 0 - 1 0 0 0 0 0 1 ? 1 0 1 ? ? ? ? ? ? 1 - - ? 0 2 1 2 5 0 ? 0 1 2 1 ? 0 1 1 3 1 - - ? ? 0 0 1 ? ? ? ? ? ? ? ? ? ? ? ? ? ? ? ? ? ? ? ? ? ? ? ? ? ? ? ? ? ? ? ? ? ? ? ? ? ? ? ? ? ? ? ? ? ? ? ? ? ? ? ? ? ? ? ? ? ? ? ? ? ? ? ? ? ? ? ? ? ? ? ? ? ? ? ? ? ? ? ? ? ? ? ? ? ? ? ? ? ? ?

Ludodactylus_sibbicki 1 - 1 0 1 0 0 0 1 0 1 ? ? 1 - 1 0 - - 1 1 0 1 0 2 1 1 0 - 1 0 - - - - - - - 0 - - - - 0 1 1 0 1 1 1 1 1 1 1 1 2 0 0 1 2 2 0 1 1 1 0 1 0 0 0 - 1 1 1 1 0 0 ? 0 1 0 1 0 1 0 ? 1 1 1 0 0 1 0 - 1 0 0 0 0 1 1 - 1 0 0 ? ? 1 0 1 0 0 0 - 1 0 2 1 2 5 1 1 0 1 2 1 1 0 1 1 3 1 - - 1 0 0 0 1 ? ? ? ? ? ? ? ? ? ? ? ? ? ? ? ? ? ? ? ? ? ? ? ? ? ? ? ? ? ? ? ? ? ? ? ? ? ? ? ? ? ? ? ? ? ? ? ? ? ? ? ? ? ? ? ? ? ? ? ? ? ? ? ? ? ? ? ? ? ? ? ? ? ? ? ? ? ? ? ? ? ? ? ? ? ? ? ? ? ? ?

Cearadactylus_atrox 1 - 1 0 1 0 0 0 1 0 1 0 0 1 - 1 0 - - 1 1 0 1 0 ? 1 1 0 - ? 1 ? 0 1 0 0 1 1 0 - - - - 0 1 1 0 0 ? ? ? ? ? ? ? ? ? 0 1 2 3 0 1 0 1 0 1 0 1 ? - 1 ? ? ? ? ? ? 0 1 0 0 0 1 1 1 1 1 1 0 0 1 0 - 1 0 0 0 0 0 1 - 1 0 1 0 - 1 0 1 0 1 - - 1 0 2 1 2 5 0 1 0 1 2 1 1 0 1 1 3 1 - - 1 0 0 0 1 ? ? ? ? ? ? ? ? ? ? ? ? ? ? ? ? ? ? ? ? ? ? ? ? ? ? ? ? ? ? ? ? ? ? ? ? ? ? ? ? ? ? ? ? ? ? ? ? ? ? ? ? ? ? ? ? ? ? ? ? ? ? ? ? ? ? ? ? ? ? ? ? ? ? ? ? ? ? ? ? ? ? ? ? ? ? ? ? ? ? ?

Piksi_barbarulna ? ? ? ? ? ? ? ? ? ? ? ? ? ? ? ? ? ? ? ? ? ? ? ? ? ? ? ? ? ? ? ? ? ? ? ? ? ? ? ? ? ? ? ? ? ? ? ? ? ? ? ? ? ? ? ? ? ? ? ? ? ? ? ? ? ? ? ? ? ? ? ? ? ? ? ? ? ? ? ? ? ? ? ? ? ? ? ? ? ? ? ? ? ? ? ? ? ? ? ? ? ? ? ? ? ? ? ? ? ? ? ? ? ? ? ? ? ? ? ? ? ? ? ? ? ? ? ? ? ? ? ? ? ? ? ? ? ? ? ? ? ? ? ? ? ? ? ? ? ? ? ? ? ? ? ? ? ? ? ? ? ? ? ? ? ? ? ? ? ? ? ? ? ? ? 1 0 0 2 ? ? ? ? ? 0 0 ? ? ? ? ? ? ? ? ? ? ? ? ? ? ? ? ? ? ? ? ? ? ? ? ? ? ? ? ? ? ? ? ? ? 1 0 0 0 ? ? 0 ? ? ?

Pteranodon_longiceps 2 - 0 0 0 - 0 0 1 0 1 0 0 2 - - 0 - - 1 1 0 1 0 4 1 1 0 - 1 0 - - - - - - - 0 - - - - 0 1 0 ? 0 1 2 0 0 1 0 1 2 0 0 1 2 4 0 1 1 1 0 1 0 1 0 - 1 1 1 1 0 0 1 0 0 - - 0 1 0 1 1 1 1 1 0 0 0 - 1 1 0 0 0 1 1 - 0 0 0 1 1 0 0 1 0 1 - - 1 1 - - - - - - - - - - - - - - - - - - - - - - 0 1 0 1 1 0 1 1 0 1 0 1 1 1 1 1 1 0 1 1 1 0 1 0 2 0 0 1 0 1 0 1 1 0 0 0 0 0 1 0 3 0 7 1 1 0 0 0 0 1 1 4 0 2 1 1 0 0 1 1 0 1 0 1 1 2 0 1 1 0 1 1 0 1 1 1 1 0 1 1 4 - 1 0 0 0 0 0 0 1 1 0

Pteranodon_sternbergi 2 - 0 ? 0 - ? 0 1 0 1 0 0 2 - - 0 - - 1 1 0 1 0 4 1 1 0 - 1 0 - - - - - - - 0 - - - - 0 1 0 0 ? 1 2 0 0 1 0 1 2 0 0 1 2 3 0 1 1 1 0 1 0 1 0 - 1 1 ? ? 0 0 ? ? ? ? ? ? ? 0 ? ? 1 1 1 0 0 0 - 1 1 0 ? 0 1 1 - ? ? ? ? ? 0 0 1 0 1 ? - ? 1 - - - - - - - - - - - - - - - - - - - - - - ? ? ? ? ? ? ? ? 0 ? ? ? ? ? 1 1 ? ? 1 ? ? ? ? ? 2 0 ? 1 0 1 0 1 ? 0 ? 0 ? 0 1 ? 3 ? 7 ? ? ? ? 0 ? 1 ? 2 0 ? ? 1 0 0 ? ? 0 ? ? ? ? ? ? ? ? ? ? ? ? 1 ? ? ? 0 1 ? ? ? 1 0 0 0 0 0 0 1 1 0

Tethydraco_regalis ? ? ? ? ? ? ? ? ? ? ? ? ? ? ? ? ? ? ? ? ? ? ? ? ? ? ? ? ? ? ? ? ? ? ? ? ? ? ? ? ? ? ? ? ? ? ? ? ? ? ? ? ? ? ? ? ? ? ? ? ? ? ? ? ? ? ? ? ? ? ? ? ? ? ? ? ? ? ? ? ? ? ? ? ? ? ? ? ? ? ? ? ? ? ? ? ? ? ? ? ? ? ? ? ? ? ? ? ? ? ? ? ? ? ? ? ? ? ? ? ? ? ? ? ? ? ? ? ? ? ? ? ? ? ? ? ? ? ? ? ? ? ? ? ? ? ? ? ? ? ? ? ? ? ? ? ? ? ? ? ? ? ? ? ? ? ? ? ? ? ? 0 ? 0 0 1 1 0 3 0 7 1 1 0 ? ? ? ? ? ? ? ? ? ? ? ? ? ? ? ? ? ? ? ? ? ? ? ? ? ? ? ? ? ? ? ? ? ? ? ? 1 0 0 0 0 0 0 1 1 0

Alamodactylus_byrdi ? ? ? ? ? ? ? ? ? ? ? ? ? ? ? ? ? ? ? ? ? ? ? ? ? ? ? ? ? ? ? ? ? ? ? ? ? ? ? ? ? ? ? ? ? ? ? ? ? ? ? ? ? ? ? ? ? ? ? ? ? ? ? ? ? ? ? ? ? ? ? ? ? ? ? ? ? ? ? ? ? ? ? ? ? ? ? ? ? ? ? ? ? ? ? ? ? ? ? ? ? ? ? ? ? ? ? ? ? ? ? ? ? ? ? ? ? ? ? ? ? ? ? ? ? ? ? ? ? ? ? ? ? ? ? ? ? ? ? ? ? ? ? ? ? ? ? ? ? ? ? ? ? ? ? ? ? ? ? ? ? ? ? ? ? ? ? ? ? ? 1 0 0 0 0 1 0 ? 3 0 6 1 1 0 ? ? ? ? ? ? ? ? ? ? 0 0 ? ? ? ? ? ? ? ? ? ? ? ? ? ? ? ? ? ? ? ? ? ? ? ? ? ? ? ? ? ? 0 ? 1 0

Tupandactylus_navigans 2 - 2 1 0 - 1 1 0 0 1 0 ? 0 - - 1 - - 1 0 0 1 2 4 1 1 0 - 1 1 1 1 0 1 1 1 0 0 - - - - 0 0 1 1 0 ? ? ? 0 1 0 ? 0 0 0 1 2 3 0 1 0 1 0 0 1 1 0 - 1 1 ? ? 0 ? ? 0 ? ? ? ? ? 1 ? ? ? ? ? ? ? ? ? ? ? ? ? ? ? ? ? ? ? ? ? ? ? ? ? ? ? ? ? ? 1 - - - - - - - - - - - - - - - - - - - - - - 0 ? ? ? ? ? ? ? ? ? ? ? ? ? ? ? ? ? ? ? ? ? ? ? ? ? ? ? ? ? ? ? ? ? ? ? ? ? ? ? ? ? ? ? ? ? ? ? ? ? ? ? ? ? ? ? ? ? ? ? ? ? ? ? ? ? ? ? ? ? ? ? ? ? ? ? ? ? ? ? ? ? ? ? ? ? 0 ? ? ? ? ?

Tupandactylus_imperator 2 - 2 ? 0 - 1 ? 0 0 1 0 ? 0 - - ? - - 1 ? 0 1 2 3 1 1 0 - 1 1 1 1 0 1 1 1 0 0 - - - - 1 - - - - 1 1 2 0 ? 0 1 3 0 0 1 2 3 0 1 0 1 0 0 1 1 0 - 1 1 ? ? ? 1 ? 0 ? ? ? ? ? ? ? ? 1 2 2 0 0 1 0 1 0 ? ? ? ? 1 - 0 1 0 ? ? 0 0 1 ? 2 1 0 ? 1 - - - - - - - - - - - - - - - - - - - - - - 0 ? ? ? ? ? ? ? ? ? ? ? ? ? ? ? ? ? ? ? ? ? ? ? ? ? ? ? ? ? ? ? ? ? ? ? ? ? ? ? ? ? ? ? ? ? ? ? ? ? ? ? ? ? ? ? ? ? ? ? ? ? ? ? ? ? ? ? ? ? ? ? ? ? ? ? ? ? ? ? ? ? ? ? ? ? 0 ? ? ? ? ?

Tapejara_wellnhoferi 2 - 2 1 0 - 1 1 0 0 1 0 0 0 - - 1 - - 1 0 0 1 2 4 1 1 0 - 1 1 1 1 0 1 0 1 1 0 - - - - 0 0 1 1 0 1 1 2 0 1 0 1 3 0 0 1 2 3 0 1 0 1 0 0 1 1 0 - 1 1 1 1 0 1 1 1 0 - - 0 1 0 1 0 1 2 1 0 0 1 1 1 0 0 1 1 1 1 - 0 1 0 1 0 0 0 1 0 2 1 0 0 1 - - - - - - - - - - - - - - - - - - - - - - 0 ? 0 1 1 0 0 1 0 1 0 1 0 0 ? ? ? ? 0 1 0 0 1 2 2 1 1 1 0 0 0 1 1 0 1 0 1 1 0 0 1 0 8 0 1 1 1 1 1 0 0 4 1 ? ? 1 0 0 0 1 0 3 1 1 ? ? 0 0 1 1 1 ? 1 1 1 1 1 1 0 1 4 - 1 0 0 0 0 0 0 1 0 0

Europejara_olcadesorum ? ? ? ? ? - ? ? 0 ? ? 0 ? ? ? ? ? - - 1 ? ? ? ? 4 1 1 ? - ? ? ? ? ? ? ? ? ? 0 - - - - ? ? ? ? ? ? ? ? ? 1 ? ? ? 0 0 ? 2 ? 0 ? 0 1 0 ? ? ? 0 ? 1 ? ? ? ? ? ? 1 ? ? ? 0 1 0 ? 0 1 2 1 0 0 1 1 1 0 ? 1 1 1 1 - 0 1 ? ? ? ? 0 1 0 2 1 0 ? 1 - - - - - - - - - - - - - - - - - - - - - - 0 ? ? ? ? ? ? ? ? ? ? ? ? ? ? ? ? ? ? ? ? ? ? ? ? ? ? ? ? ? ? ? ? ? ? ? ? ? ? ? ? ? ? ? ? ? ? ? ? ? ? ? ? ? ? ? ? ? ? ? ? ? ? ? ? ? ? ? ? ? ? ? ? ? ? ? ? ? ? ? ? ? ? ? ? ? 0 ? ? ? ? ?

Vectidraco_daisymorrisae ? ? ? ? ? ? ? ? ? ? ? ? ? ? ? ? ? ? ? ? ? ? ? ? ? ? ? ? ? ? ? ? ? ? ? ? ? ? ? ? ? ? ? ? ? ? ? ? ? ? ? ? ? ? ? ? ? ? ? ? ? ? ? ? ? ? ? ? ? ? ? ? ? ? ? ? ? ? ? ? ? ? ? ? ? ? ? ? ? ? ? ? ? ? ? ? ? ? ? ? ? ? ? ? ? ? ? ? ? ? ? ? ? ? ? ? ? ? ? ? ? ? ? ? ? ? ? ? ? ? ? ? ? ? ? ? ? ? ? ? ? ? ? ? ? ? ? ? ? ? ? ? 1 0 ? ? ? ? ? ? ? ? ? ? ? ? ? ? ? ? ? ? ? ? ? ? ? ? ? ? ? ? ? ? ? ? ? ? ? ? ? ? ? ? ? ? ? ? ? 3 1 1 ? ? ? 0 1 1 1 1 1 ? ? ? ? ? ? ? ? ? ? ? ? ? ? ? ? ? ? ?

Caiuajara_dobruskii 2 - 2 1 0 - 1 1 0 0 1 0 0 0 - - ? - - 1 0 0 1 2 4 1 1 0 - ? 1 1 1 0 1 1 1 1 0 - - - - ? ? ? ? ? 1 1 2 ? ? ? 1 3 ? 0 1 ? [34] 0 ? 0 1 0 0 1 1 0 - 1 1 ? 1 0 1 ? 0 0 - - 0 ? ? ? 0 1 2 2 0 0 1 1 1 0 0 1 0 1 1 - 0 1 0 ? ? 0 0 1 0 2 1 0 0 1 - - - - - - - - - - - - - - - - - - - - - - 0 ? ? 1 1 0 0 1 0 ? ? ? 0 ? 1 ? 1 0 0 1 0 0 1 2 2 ? 1 ? 0 0 ? 2 1 ? 1 0 1 ? ? ? 1 0 8 0 1 1 ? ? 1 0 0 4 ? ? ? 1 0 0 ? 1 0 2 ? ? 1 2 ? ? 1 1 1 ? 1 1 1 1 1 1 0 ? ? ? 1 0 0 0 0 0 0 1 0 0

Huaxiapterus_benxiensis 2 - 2 ? 0 - 1 ? 0 0 1 0 ? 0 - - ? - - 1 0 0 ? ? ? ? 1 0 - 1 1 1 2 6 1 0 1 1 0 - - - - 0 - 1 0 ? 1 1 2 0 ? 0 1 3 ? 0 1 3 3 0 1 0 1 0 0 1 1 0 - 1 1 ? ? ? ? ? ? ? ? ? ? ? ? ? ? 1 2 1 0 0 1 0 1 0 0 1 1 1 1 - 0 1 0 ? ? 2 0 1 2 2 0 1 ? 1 - - - - - - - - - - - - - - - - - - - - - - 0 ? 0 1 ? ? ? 1 0 1 ? ? 0 ? ? ? ? ? ? ? ? ? ? ? ? ? ? ? ? ? ? ? ? ? ? 0 1 ? ? ? ? 0 8 0 ? ? ? ? ? ? 0 ? ? 2 1 1 0 ? 1 1 0 ? ? ? ? ? ? ? ? ? ? ? ? 1 ? ? ? ? ? ? 4 - 1 0 0 0 0 0 0 1 0 0

Huaxiapterus_corallatus 2 - 2 ? 0 - 1 1 0 0 1 0 ? 0 - - ? - - 1 ? 0 ? 2 ? ? ? 0 - 1 1 1 2 6 1 0 1 1 0 - - - - ? ? ? ? ? ? ? ? 0 ? ? 1 0 ? ? ? ? ? ? ? 0 1 ? ? ? ? ? ? 1 ? ? ? ? 1 ? ? ? ? ? ? ? ? ? ? 1 2 1 0 0 1 0 1 0 0 1 ? 1 1 - ? ? ? ? ? 2 0 ? 2 2 0 1 1 1 - - - - - - - - - - - - - - - - - - - - - - 0 ? 0 1 ? ? ? 1 0 1 ? ? 0 ? ? ? ? ? ? ? 0 0 ? 2 2 ? ? 1 0 0 0 ? ? ? ? 0 1 ? ? ? ? 0 8 0 1 ? 1 1 1 ? ? 4 ? 2 1 1 0 ? 1 1 0 ? ? ? 1 3 ? ? ? 1 1 ? 1 1 1 ? 1 ? ? 1 4 - 1 0 0 0 0 0 0 1 1 0

Eopteranodon_lii 2 - 2 1 0 - 1 1 0 0 1 0 0 0 - - ? - - 1 0 0 ? ? ? ? ? 0 - ? 1 1 0 4 1 0 1 1 0 - - - - ? ? ? ? ? ? ? ? ? ? ? ? ? ? ? 1 ? ? 0 ? 0 1 ? 0 1 1 ? - 1 1 ? ? ? ? ? ? ? - - ? ? ? ? 1 ? 2 1 0 0 1 0 1 0 ? ? 1 ? 1 - 0 0 ? ? ? ? ? 1 ? ? ? ? 1 1 - - - - - - - - - - - - - - - - - - - - - - 0 ? ? ? ? 0 0 1 0 1 ? ? ? ? ? ? ? ? ? ? ? 0 ? 2 2 ? ? ? ? 0 0 ? ? 0 ? 0 1 ? ? ? 1 0 8 0 1 ? ? 1 ? ? ? 4 ? ? ? ? ? 0 ? ? 0 2 0 1 1 3 0 1 1 1 1 1 1 1 ? 1 1 1 0 ? ? ? 1 0 0 0 0 0 0 1 0 0

Huaxiapterus_jii 2 - 2 1 0 - 1 ? ? 0 1 0 ? 0 - - ? - - 1 0 0 ? ? 4 1 1 0 - ? 1 1 0 4 1 0 ? 1 0 - - - - 0 0 ? 1 0 ? ? ? ? ? ? ? 0 ? ? 1 3 ? 0 1 ? 1 ? 0 1 1 ? - 1 ? ? ? ? ? ? ? 0 - - 0 ? ? ? ? 1 2 1 0 0 1 0 1 0 0 ? ? ? 1 - 0 0 ? ? ? 2 0 1 2 2 0 1 1 1 - - - - - - - - - - - - - - - - - - - - - - 0 ? ? ? ? ? ? ? 0 1 ? ? 0 ? ? ? 1 ? ? ? ? ? ? 2 2 ? ? 1 0 0 0 2 1 0 ? 0 1 ? ? ? 1 0 8 0 1 1 ? ? ? ? ? 4 1 1 1 1 0 ? 1 1 0 ? ? ? 1 3 ? ? ? ? ? ? ? 1 ? 1 1 1 0 ? ? ? 1 0 0 0 0 0 0 1 0 0

Sinopterus_dongi 2 - 2 ? 0 - 1 1 0 0 1 0 ? 0 - - 1 - - 1 0 0 0 2 ? 1 ? 0 - 1 1 1 0 4 1 0 1 1 0 - - - - 0 0 1 1 0 1 1 2 0 1 ? 1 3 0 ? 1 3 3 ? 1 0 1 ? 0 1 1 ? - 1 1 ? ? ? 1 ? ? ? ? ? ? ? ? ? ? 1 2 1 0 0 1 0 1 0 0 1 ? 1 1 - ? ? ? ? ? 2 0 1 2 2 0 1 1 1 - - - - - - - - - - - - - - - - - - - - - - 0 ? ? ? ? 0 0 1 0 1 ? ? 0 ? ? 1 1 0 ? 1 ? 0 ? 2 2 ? 1 1 0 0 0 2 1 0 ? 0 1 1 ? ? 1 0 8 0 1 1 1 1 1 0 0 4 1 2 1 1 0 0 1 1 0 ? 0 ? 1 3 ? 1 1 1 1 1 1 1 ? ? ? 1 ? 1 4 - 1 0 0 0 0 0 0 1 0 0

Bennettazhia_oregonensis ? ? ? ? ? ? ? ? ? ? ? ? ? ? ? ? ? ? ? ? ? ? ? ? ? ? ? ? ? ? ? ? ? ? ? ? ? ? ? ? ? ? ? ? ? ? ? ? ? ? ? ? ? ? ? ? ? ? ? ? ? ? ? ? ? ? ? ? ? ? ? ? ? ? ? ? ? ? ? ? ? ? ? ? ? ? ? ? ? ? ? ? ? ? ? ? ? ? ? ? ? ? ? ? ? ? ? ? ? ? ? ? ? ? ? ? ? ? ? ? ? ? ? ? ? ? ? ? ? ? ? ? ? ? ? ? ? ? ? ? ? ? ? ? ? ? ? ? ? ? 1 0 ? ? ? ? ? ? ? ? ? ? ? ? ? ? ? ? ? ? 1 0 1 0 0 1 0 0 1 0 8 0 1 1 ? ? ? ? ? ? ? ? ? ? ? ? ? ? ? ? ? ? ? ? ? ? ? ? ? ? ? ? ? ? ? ? ? ? ? ? ? ? ? ? ? ? 0 ? 0 0

Microtuban_altivolans ? ? ? ? ? ? ? ? ? ? ? ? ? ? ? ? ? ? ? ? ? ? ? ? ? ? ? ? ? ? ? ? ? ? ? ? ? ? ? ? ? ? ? ? ? ? ? ? ? ? ? ? ? ? ? ? ? ? ? ? ? ? ? ? ? ? ? ? ? ? ? ? ? ? ? ? ? ? ? ? ? ? ? ? ? ? ? ? ? ? ? ? ? ? ? ? ? ? ? ? ? ? ? ? ? ? ? ? ? ? ? ? ? ? ? ? ? ? ? ? ? ? ? ? ? ? ? ? ? ? ? ? ? ? ? ? ? ? ? ? ? ? ? ? ? ? ? ? ? 1 ? ? ? ? ? ? ? 1 0 0 ? 2 2 ? ? ? ? ? ? ? ? 0 0 0 1 1 0 ? ? 0 8 0 1 1 ? ? 1 ? ? 4 ? 2 1 1 0 ? ? ? 0 ? ? ? ? ? ? ? ? ? ? ? ? ? 1 1 2 ? 0 ? ? ? 1 0 0 0 ? ? 0 1 0 0

Chaoyangopterus_zhangi 2 - 1 0 0 - 1 0 0 0 1 0 ? 2 - - ? - - 1 ? 0 ? ? ? ? ? 1 - ? 0 - - - - - - - 0 - - - - ? ? ? ? ? ? ? ? ? ? ? ? ? ? ? 1 ? ? ? ? ? ? ? ? ? ? ? - ? ? ? ? ? ? ? ? ? ? ? ? ? ? ? ? 1 1 1 0 0 0 - 1 1 0 1 0 0 1 - ? ? ? ? ? ? ? ? ? 1 - - ? 1 - - - - - - - - - - - - - - - - - - - - - - 0 ? 0 0 ? 1 2 1 1 1 ? ? ? ? ? ? ? ? ? ? 0 0 ? 2 2 1 1 1 0 0 0 2 ? ? 0 0 1 1 ? ? 1 0 ? ? ? ? 1 ? 1 ? ? 4 ? 1 1 1 0 0 ? ? 0 2 0 1 1 3 0 0 0 0 0 ? 0 1 ? 1 2 1 0 1 4 - 1 0 0 0 0 0 0 1 0 0

Jidapterus_edentus 2 - 1 0 0 - 1 0 0 0 1 0 0 2 - - 1 - - 1 ? 0 ? ? ? ? ? 1 - ? 0 - - - - - - - 0 - - - - ? ? ? ? ? ? ? ? ? ? ? ? ? ? ? 1 ? ? ? 1 0 1 ? ? ? ? ? - 1 ? ? ? ? ? ? 0 0 ? ? ? ? ? ? ? 1 1 1 0 0 0 - 1 1 0 1 0 0 1 - ? ? ? ? ? 0 ? 1 0 1 - - 1 1 - - - - - - - - - - - - - - - - - - - - - - 0 ? 0 0 ? 1 2 1 1 1 ? ? ? ? 1 ? ? ? ? ? ? 0 ? ? 2 1 ? 1 0 0 0 1 ? ? ? 0 1 ? ? 0 ? ? ? ? ? ? ? 1 1 0 ? 4 ? 1 1 1 0 0 1 1 ? 3 ? ? 1 3 ? ? ? 1 1 ? 1 1 1 1 2 1 ? ? 3 3 1 0 0 0 0 0 0 1 0 0

Samrukia_nessovi 2 - 1 0 0 - 1 0 0 0 1 0 ? 2 - - ? - - ? ? ? ? ? ? ? ? ? - ? ? - - - - - - - ? - - - - ? ? ? ? ? ? ? ? ? ? ? ? ? ? ? ? ? ? ? ? ? ? ? ? ? ? ? - ? ? ? ? ? ? ? ? ? ? ? ? ? ? ? ? ? ? ? ? ? ? - ? ? ? ? ? ? ? - ? ? ? ? ? ? ? ? ? ? - - ? ? - - - - - - - - - - - - - - - - - - - - - - ? ? ? ? ? ? ? ? ? ? ? ? ? ? ? ? ? ? ? ? ? ? ? ? ? ? ? ? ? ? ? ? ? ? ? ? ? ? ? ? ? ? ? ? ? ? ? ? ? ? ? ? ? ? ? ? ? ? ? ? ? ? ? ? ? ? ? ? ? ? ? ? ? ? ? ? ? ? ? ? ? - ? ? ? ? ? ? ? ? ? ?

Shenzhoupterus_chaoyangensis 2 - 1 ? 0 - ? ? ? 0 1 ? ? 2 - - 1 - - 1 0 0 1 2 ? 1 1 1 - 1 0 - - - - - - - 0 - - - - 1 - - - ? 1 1 0 1 ? 0 1 3 0 1 ? 2 3 0 1 ? 1 0 0 0 2 0 ? 1 1 ? ? ? 1 ? ? ? ? ? ? ? ? ? ? 1 1 1 0 0 0 - 1 1 0 ? ? ? 1 - ? ? ? ? ? 0 0 1 ? 1 - - ? 1 - - - - - - - - - - - - - - - - - - - - - - 0 ? ? ? ? 1 2 ? 1 1 ? 1 0 ? ? ? ? ? ? ? 0 0 ? 2 2 ? ? ? ? ? ? ? ? ? ? 0 1 ? ? ? 1 0 8 0 ? ? ? ? ? ? 0 ? ? 2 1 ? 0 ? 1 ? 0 ? ? ? ? ? ? ? ? ? ? ? ? 1 ? 1 ? ? 0 1 4 - 1 0 0 0 0 0 0 1 0 0

Cretornis_hlavaci ? ? ? ? ? ? ? ? ? ? ? ? ? ? ? ? ? ? ? ? ? ? ? ? ? ? ? ? ? ? ? ? ? ? ? ? ? ? ? ? ? ? ? ? ? ? ? ? ? ? ? ? ? ? ? ? ? ? ? ? ? ? ? ? ? ? ? ? ? ? ? ? ? ? ? ? ? ? ? ? ? ? ? ? ? ? ? ? ? ? ? ? ? ? ? ? ? ? ? ? ? ? ? ? ? ? ? ? ? ? ? ? ? ? ? ? ? ? ? ? ? ? ? ? ? ? ? ? ? ? ? ? ? ? ? ? ? ? ? ? ? ? ? ? ? ? ? ? ? ? ? ? ? ? ? ? ? ? ? ? ? ? ? ? ? ? ? ? ? ? 1 0 0 0 0 1 0 0 3 1 ? 1 1 0 ? 1 0 ? ? ? ? 2 ? ? 0 0 ? ? 0 ? ? ? ? ? ? ? ? ? ? ? ? ? ? ? ? ? ? ? ? ? 1 0 0 0 0 0 0 1 0 0

Eoazhdarcho_liaoxiensis ? ? ? ? ? - ? ? ? ? ? 0 0 ? ? ? ? ? ? ? ? ? ? ? ? ? ? ? ? ? ? ? ? ? ? ? ? ? ? ? ? ? ? ? ? ? ? ? ? ? ? ? ? ? ? ? ? ? ? ? ? 0 ? ? ? ? ? ? ? ? ? ? ? ? ? ? ? ? ? ? - - ? ? ? ? ? ? 1 1 0 0 0 - 1 ? ? 1 0 0 1 - 0 0 0 1 0 ? ? 1 0 ? - ? 1 1 - - - - - - - - - - - - - - - - - - - - - - ? ? 0 0 ? 1 2 ? 1 1 ? ? 1 ? ? ? ? ? ? ? ? 0 ? 2 2 ? ? ? ? 0 0 ? ? 0 ? 0 1 ? ? 0 ? 0 8 0 ? ? 1 1 1 ? ? 4 ? 2 1 1 0 ? ? 1 0 ? ? ? ? ? ? ? ? ? ? ? ? 1 ? ? ? 1 ? ? ? ? 1 0 0 0 0 0 0 1 0 0

Radiodactylus_langstoni ? ? ? ? ? ? ? ? ? ? ? ? ? ? ? ? ? ? ? ? ? ? ? ? ? ? ? ? ? ? ? ? ? ? ? ? ? ? ? ? ? ? ? ? ? ? ? ? ? ? ? ? ? ? ? ? ? ? ? ? ? ? ? ? ? ? ? ? ? ? ? ? ? ? ? ? ? ? ? ? ? ? ? ? ? ? ? ? ? ? ? ? ? ? ? ? ? ? ? ? ? ? ? ? ? ? ? ? ? ? ? ? ? ? ? ? ? ? ? ? ? ? ? ? ? ? ? ? ? ? ? ? ? ? ? ? ? ? ? ? ? ? ? ? ? ? ? ? ? ? ? ? ? ? ? ? ? ? ? ? ? ? ? ? ? ? ? ? ? ? 1 0 0 0 1 ? 0 1 1 0 8 0 1 1 ? ? ? ? ? ? ? ? ? ? ? ? ? ? ? ? ? ? ? ? ? ? ? ? ? ? ? ? ? ? ? ? ? ? ? ? ? ? ? ? ? ? ? ? ? ?

Montanazhdarcho_minor ? ? ? ? ? - ? ? ? ? ? 0 ? ? ? ? ? ? ? ? ? ? ? ? ? ? ? ? ? ? ? ? ? ? ? ? ? ? ? ? ? ? ? ? ? ? ? ? ? ? ? ? ? ? ? ? ? ? ? ? ? ? ? ? ? ? ? ? ? ? ? ? ? ? ? ? ? ? ? ? ? ? ? ? ? ? ? 1 1 ? 0 0 0 - 1 ? 0 1 0 0 ? ? 0 0 0 ? ? ? ? ? ? 0 - - ? 1 - - - - - - - - - - - - - - - - - - ? ? - - ? ? 0 0 ? 1 2 ? 1 1 ? ? ? ? ? ? ? ? ? 1 0 0 1 2 2 ? 0 1 0 ? ? ? 1 1 0 0 1 1 0 1 1 0 8 0 0 1 ? 1 1 0 ? ? 1 ? ? 1 0 0 ? ? 0 ? ? ? ? ? ? ? ? ? ? ? ? ? ? ? ? ? ? ? ? ? 1 0 0 0 0 0 0 1 0 0

RBCM_EH_2009_019_0001 ? ? ? ? ? ? ? ? ? ? ? ? ? ? ? ? ? ? ? ? ? ? ? ? ? ? ? ? ? ? ? ? ? ? ? ? ? ? ? ? ? ? ? ? ? ? ? ? ? ? ? ? ? ? ? ? ? ? ? ? ? ? ? ? ? ? ? ? ? ? ? ? ? ? ? ? ? ? ? ? ? ? ? ? ? ? ? ? ? ? ? ? ? ? ? ? ? ? ? ? ? ? ? ? ? ? ? ? ? ? ? ? ? ? ? ? ? ? ? ? ? ? ? ? ? ? ? ? ? ? ? ? ? ? ? ? ? ? ? ? ? ? ? ? ? ? ? ? ? ? ? ? ? ? ? ? ? ? ? ? ? ? ? ? ? ? ? ? ? ? 1 0 0 0 0 1 0 0 3 1 ? 1 1 0 ? 1 0 ? ? ? ? 2 ? ? 0 0 ? ? 0 ? ? ? ? ? ? ? ? ? ? ? ? ? ? ? ? ? ? ? ? ? ? ? ? ? 0 0 0 1 0 0

Palaeocursornis_corneti ? ? ? ? ? ? ? ? ? ? ? ? ? ? ? ? ? ? ? ? ? ? ? ? ? ? ? ? ? ? ? ? ? ? ? ? ? ? ? ? ? ? ? ? ? ? ? ? ? ? ? ? ? ? ? ? ? ? ? ? ? ? ? ? ? ? ? ? ? ? ? ? ? ? ? ? ? ? ? ? ? ? ? ? ? ? ? ? ? ? ? ? ? ? ? ? ? ? ? ? ? ? ? ? ? ? ? ? ? ? ? ? ? ? ? ? ? ? ? ? ? ? ? ? ? ? ? ? ? ? ? ? ? ? ? ? ? ? ? ? ? ? ? ? ? ? ? ? ? ? ? ? ? ? ? ? ? ? ? ? ? ? ? ? ? ? ? ? ? ? 1 0 0 0 1 ? 0 1 1 0 8 0 1 1 ? ? ? ? ? ? ? ? ? ? ? ? ? ? ? ? ? ? ? ? ? ? ? ? ? ? ? ? ? ? ? ? ? ? ? ? ? ? ? ? ? ? ? ? ? ?

Tupuxuara_longicristatus ? ? ? 0 0 - 1 0 0 0 1 0 ? ? - - ? - - 1 0 0 ? ? ? ? ? 0 - 1 1 0 0 5 ? 0 1 2 ? - - - - ? ? ? ? ? ? ? ? ? ? ? ? ? ? ? 1 ? ? ? ? 0 ? ? 0 ? ? ? - ? ? ? ? ? ? ? 1 1 1 1 0 ? ? ? ? ? ? ? ? ? ? ? ? ? ? ? ? ? ? ? ? ? ? ? ? ? ? ? ? ? ? ? ? 1 - - - - - - - - - - - - - - - - - - - - - - 0 ? ? ? ? ? ? ? 0 ? ? ? ? ? ? ? ? ? ? ? ? ? ? ? ? ? ? ? ? ? 0 ? ? ? ? ? ? ? ? ? ? ? ? ? ? ? ? ? ? ? ? ? ? 2 1 ? 0 ? ? 1 ? ? ? ? ? ? ? ? ? ? ? ? ? ? ? ? ? ? ? ? ? ? ? ? ? ? ? ? ? ? ? ?

Tupuxuara_leonardii 2 - 1 0 0 - 1 0 0 0 1 0 0 1 - - 1 - - 1 0 0 1 2 4 1 1 0 - 1 1 0 0 5 3 0 1 2 0 - - - - 1 - - - - 1 1 0 0 0 0 1 2 2 0 1 2 3 0 1 0 1 0 0 1 1 0 - 1 1 1 1 0 1 1 1 1 1 1 0 1 1 1 1 1 1 1 0 0 0 - 1 0 0 1 0 1 1 - 0 0 0 1 0 0 0 1 0 1 - - 1 1 - - - - - - - - - - - - - - - - - - - - - - 0 1 0 1 1 0 0 1 0 1 ? ? 1 1 ? ? ? ? ? ? 0 0 1 1 2 1 ? 1 0 0 0 1 1 0 0 ? ? 1 1 ? 1 0 8 0 1 1 1 1 1 0 0 4 ? ? ? 1 0 0 1 1 0 ? ? ? ? ? ? ? ? ? ? ? ? 1 1 1 2 1 0 1 ? ? 1 0 0 0 0 0 0 1 0 0

Alanqa_saharica 2 - 1 0 0 - 1 0 ? 3 1 ? ? 1 ? ? ? ? ? ? ? ? ? ? ? ? ? ? ? ? ? ? ? ? ? ? ? ? ? ? ? ? ? ? ? ? ? ? ? ? ? ? ? ? ? ? ? ? ? ? ? ? ? ? ? ? ? ? ? ? ? ? ? ? ? ? ? ? 0 1 2 1 ? ? ? ? ? 0 1 ? ? 0 0 - 1 ? ? 1 0 1 ? ? 1 0 1 ? ? ? ? ? ? ? ? ? ? 1 - - - - - - - - - - - - - - - - - - - - - ? 0 ? ? ? ? ? ? ? ? ? ? ? ? ? ? ? ? ? ? ? ? ? ? ? ? ? ? ? ? ? ? ? ? ? ? ? ? ? ? ? ? ? ? ? ? ? ? ? ? ? ? ? ? ? ? ? ? ? ? ? ? ? ? ? ? ? ? ? ? ? ? ? ? ? ? ? ? ? ? ? ? ? ? ? ? ? ? ? ? ? ? ?

Aerotitan_sudamericanus 2 - 1 0 0 - 1 0 ? 3 1 ? ? 1 ? ? ? ? ? ? ? ? ? ? ? ? ? ? ? ? ? ? ? ? ? ? ? ? ? ? ? ? ? ? ? ? ? ? ? ? ? ? ? ? ? ? ? ? ? ? ? ? ? ? ? ? ? ? ? ? ? ? ? ? ? ? ? ? 0 1 2 ? ? ? ? ? ? 0 ? ? ? ? ? ? ? ? ? ? ? ? ? ? ? ? ? ? ? ? ? ? ? ? ? ? ? 1 - - - - - - - - - - - - - - - - - - - - - ? 0 ? ? ? ? ? ? ? ? ? ? ? ? ? ? ? ? ? ? ? ? ? ? ? ? ? ? ? ? ? ? ? ? ? ? ? ? ? ? ? ? ? ? ? ? ? ? ? ? ? ? ? ? ? ? ? ? ? ? ? ? ? ? ? ? ? ? ? ? ? ? ? ? ? ? ? ? ? ? ? ? ? ? ? ? ? ? ? ? ? ? ?

Argentinadraco_barrealensis 2 - 1 0 0 - 1 0 ? 3 1 ? ? 1 ? ? ? ? ? ? ? ? ? ? ? ? ? ? ? ? ? ? ? ? ? ? ? ? ? ? ? ? ? ? ? ? ? ? ? ? ? ? ? ? ? ? ? ? ? ? ? ? ? ? ? ? ? ? ? ? ? ? ? ? ? ? ? ? 0 1 2 ? ? ? ? ? ? 0 ? ? ? ? ? ? ? ? ? ? ? ? ? ? ? ? ? ? ? ? ? ? ? ? ? ? ? 1 - - - - - - - - - - - - - - - - - - - - - ? 0 ? ? ? ? ? ? ? ? ? ? ? ? ? ? ? ? ? ? ? ? ? ? ? ? ? ? ? ? ? ? ? ? ? ? ? ? ? ? ? ? ? ? ? ? ? ? ? ? ? ? ? ? ? ? ? ? ? ? ? ? ? ? ? ? ? ? ? ? ? ? ? ? ? ? ? ? ? ? ? ? ? ? ? ? ? ? ? ? ? ? ?

Xericeps_curvirostris 2 - 1 0 0 - 1 0 ? 3 1 ? ? 1 ? ? ? ? ? ? ? ? ? ? ? ? ? ? ? ? ? ? ? ? ? ? ? ? ? ? ? ? ? ? ? ? ? ? ? ? ? ? ? ? ? ? ? ? ? ? ? ? ? ? ? ? ? ? ? ? ? ? ? ? ? ? ? ? ? ? ? ? ? ? ? ? ? ? ? ? ? 0 0 - ? ? ? ? ? ? ? ? ? ? ? ? ? ? ? ? ? ? ? ? ? ? - - - - - - - - - - - - - - - - - - - - - ? ? ? ? ? ? ? ? ? ? ? ? ? ? ? ? ? ? ? ? ? ? ? ? ? ? ? ? ? ? ? ? ? ? ? ? ? ? ? ? ? ? ? ? ? ? ? ? ? ? ? ? ? ? ? ? ? ? ? ? ? ? ? ? ? ? ? ? ? ? ? ? ? ? ? ? ? ? ? ? ? ? ? ? ? ? ? ? ? ? ? ? ?

Thalassodromeus_sethi 2 - 1 0 0 - 1 0 1 0 1 0 0 1 - - 1 - - 1 0 0 1 2 4 1 1 0 - 1 1 0 0 5 3 0 1 2 0 - - - - 0 1 0 0 0 1 1 0 0 0 0 1 2 2 0 1 2 3 0 1 0 1 0 0 1 1 0 - 1 1 ? 1 ? 1 1 0 1 1 1 0 1 1 ? 1 1 1 ? 0 0 0 - 1 0 0 ? 0 1 1 - 1 2 1 1 0 0 0 1 0 1 - - 1 1 - - - - - - - - - - - - - - - - - - - - - - 0 ? ? ? ? ? ? ? ? ? ? ? ? ? ? ? ? ? ? ? ? ? ? ? ? ? ? ? ? ? ? ? ? ? ? ? ? ? ? ? ? ? ? ? ? ? ? ? ? ? ? ? ? ? ? ? ? ? ? ? ? ? ? ? ? ? ? ? ? ? ? ? ? ? ? ? ? ? ? ? ? ? ? ? ? ? ? ? ? ? ? ?

Aralazhdarcho_bostobensis ? ? ? ? ? ? ? ? 0 ? ? ? ? ? ? ? 0 ? ? ? ? ? ? ? ? ? ? ? ? ? ? ? ? ? ? ? ? ? ? ? ? ? ? ? ? ? ? ? ? ? ? ? ? ? ? ? ? ? ? ? ? ? 1 ? 1 ? ? ? ? ? ? 1 ? ? ? ? ? ? ? ? ? ? ? ? ? ? 1 ? ? ? ? ? ? ? ? ? ? ? ? ? ? ? ? ? ? ? ? ? ? ? ? ? ? ? ? 1 - - - - - - - - - - - - - - - - - - - - - - ? ? ? ? 0 ? ? ? 1 ? ? ? ? ? ? ? ? ? ? 1 0 0 1 ? ? ? ? ? ? ? ? ? ? ? ? ? ? ? ? ? ? ? ? ? ? ? ? ? ? ? ? ? ? ? ? ? ? ? ? ? 3 ? ? ? ? ? ? ? ? ? ? ? ? ? 1 1 2 ? ? ? ? ? 1 0 0 0 0 0 0 1 0 0

Volgadraco_bogolubovi ? ? ? ? ? - ? ? ? ? ? 0 ? ? ? ? ? ? ? ? ? ? ? ? ? ? ? ? ? ? ? ? ? ? ? ? ? ? ? ? ? ? ? ? ? ? ? ? ? ? ? ? ? ? ? ? ? ? ? ? ? ? ? ? ? ? ? ? ? ? ? ? ? ? ? ? ? ? ? ? ? ? ? ? ? ? ? 1 1 1 0 0 0 - 1 1 0 1 ? ? ? ? ? 0 0 ? - ? ? ? ? 1 - - ? 1 - - - - - - - - - - - - - - - - - - ? ? - - ? ? 0 1 1 0 1 1 0 1 ? 1 1 ? ? ? ? ? ? ? ? ? ? ? ? ? ? ? ? ? ? ? ? ? ? ? ? ? ? ? ? ? ? ? ? ? ? ? ? ? ? ? ? ? ? ? ? ? ? ? ? ? ? ? ? ? ? ? ? ? ? ? ? ? ? 1 ? ? ? ? ? ? 1 0 0 0 0 0 0 1 0 0

Eurazhdarcho_langendorfensis ? ? ? ? ? ? ? ? ? ? ? ? ? ? ? ? ? ? ? ? ? ? ? ? ? ? ? ? ? ? ? ? ? ? ? ? ? ? ? ? ? ? ? ? ? ? ? ? ? ? ? ? ? ? ? ? ? ? ? ? ? ? ? ? ? ? ? ? ? ? ? ? ? ? ? ? ? ? ? ? ? ? ? ? ? ? ? ? ? ? ? ? ? ? ? ? ? ? ? ? ? ? ? ? ? ? ? ? ? ? ? ? ? ? ? ? ? ? ? ? ? ? ? ? ? ? ? ? ? ? ? ? ? ? ? ? ? ? ? ? 0 0 1 2 2 ? 1 1 ? ? ? ? ? ? ? ? ? ? ? ? ? ? ? ? ? ? ? ? ? ? ? ? ? ? ? ? ? ? ? ? ? ? ? ? ? ? ? ? ? ? ? ? ? 1 ? 0 ? ? 3 ? ? ? ? ? ? ? ? ? ? ? ? ? ? ? ? ? ? ? ? ? ? ? ? ? 0 ? ? ? ? ?

Phosphatodraco_mauritanicus ? ? ? ? ? ? ? ? ? ? ? ? ? ? ? ? ? ? ? ? ? ? ? ? ? ? ? ? ? ? ? ? ? ? ? ? ? ? ? ? ? ? ? ? ? ? ? ? ? ? ? ? ? ? ? ? ? ? ? ? ? ? ? ? ? ? ? ? ? ? ? ? ? ? ? ? ? ? ? ? ? ? ? ? ? ? ? ? ? ? ? ? ? ? ? ? ? ? ? ? ? ? ? ? ? ? ? ? ? ? ? ? ? ? ? ? ? ? ? ? ? ? ? ? ? ? ? ? ? ? ? ? ? ? ? ? ? ? ? ? 0 0 0 2 2 1 1 1 ? 1 ? ? ? ? ? ? ? ? ? ? ? ? ? ? ? ? ? ? ? ? ? ? ? ? ? ? ? ? ? ? ? ? ? ? ? ? ? ? ? ? ? ? ? ? ? ? ? ? ? ? ? ? ? ? ? ? ? ? ? ? ? ? ? ? ? ? ? ? ? ? ? ? ? ? ? ? ? ? ? ?

Zhejiangopterus_linhaiensis 2 - 1 0 0 - 1 0 0 0 1 ? ? 0 - - ? - - 1 0 0 1 2 4 1 1 0 - 1 0 - - - - - - - 0 - - - - 1 - - - - 0 - - ? ? 0 0 - 0 1 1 2 3 0 1 0 1 0 1 0 1 0 - 1 2 ? ? ? 1 ? ? ? ? ? ? ? 0 ? ? 1 1 1 0 0 0 - 1 1 0 1 0 0 1 - ? ? ? ? ? 0 1 1 0 1 - ? ? 1 - - - - - - - - - - - - - - - - - - - - - - 0 1 ? ? ? 2 2 ? 1 1 0 ? 1 1 1 1 1 0 0 ? 0 0 ? 2 2 ? ? 1 0 ? ? 1 ? ? ? 0 1 ? ? ? 1 1 8 0 ? ? 1 1 ? ? ? 4 ? 2 ? 1 0 ? 1 ? 3 1 0 1 1 3 ? 1 1 1 1 ? 1 1 ? 1 2 1 0 ? 4 - 1 0 0 0 0 0 0 1 0 0

Azhdarcho_lancicollis 2 - 1 0 0 - 1 0 0 0 1 0 ? 1 ? - ? - - 1 ? ? ? ? ? ? ? 0 - 1 0 - - - - - - - ? - ? - - ? ? ? ? ? ? ? ? ? ? ? ? ? ? ? ? ? ? 0 ? ? ? ? ? ? ? ? ? ? ? ? ? ? ? ? 0 0 - - ? ? ? ? 1 1 1 1 0 0 0 - 1 ? 0 1 ? ? ? ? 0 0 ? ? ? ? ? 1 ? 1 - - ? 1 - - - - - - - - - - - - - - - - - - - - - - 0 1 0 0 1 2 2 1 1 1 ? 1 1 ? ? ? ? ? ? 1 0 0 1 2 2 1 0 1 0 0 0 ? 1 ? 0 0 1 1 0 1 1 1 8 0 1 ? 1 1 1 0 0 ? ? ? ? ? 0 0 ? 1 3 ? ? ? ? ? ? ? ? ? ? ? ? 1 1 1 2 1 1 ? ? ? 1 0 0 0 0 0 0 1 0 0

Hatzegopteryx_thambena ? ? ? ? ? ? ? ? 0 ? ? ? ? ? ? ? ? ? ? ? ? ? ? ? ? ? ? ? ? ? ? ? ? ? ? ? ? ? ? ? ? ? ? ? ? ? ? ? ? ? ? ? ? ? ? ? ? ? 1 ? ? 0 1 ? 1 ? ? ? ? ? - 1 2 1 ? ? 0 ? ? ? ? ? 0 ? 1 ? 1 ? ? ? ? ? ? ? ? ? ? ? ? ? ? ? ? ? ? ? ? ? ? ? ? ? ? ? ? ? ? ? ? ? ? ? ? ? ? ? ? ? ? ? ? ? ? ? ? ? ? ? ? ? ? ? ? ? ? ? ? ? ? ? ? ? ? ? ? ? ? ? ? ? ? ? ? ? ? ? ? ? ? ? ? ? 0 0 1 ? ? ? ? 1 8 0 1 1 ? ? ? ? ? ? ? ? ? ? ? ? ? ? ? ? ? ? ? ? ? ? ? ? ? ? ? ? ? ? ? ? ? ? ? ? ? ? ? ? ? ? ? ? ? ?

Bakonydraco_galaczi ? ? ? ? ? - ? ? ? ? ? 0 0 ? ? ? ? ? ? ? ? ? ? ? ? ? ? ? ? ? ? ? ? ? ? ? ? ? ? ? ? ? ? ? ? ? ? ? ? ? ? ? ? ? ? ? ? ? ? ? ? ? ? ? ? ? ? ? ? ? ? ? ? ? ? ? ? ? ? ? ? ? ? ? ? ? 1 1 2 1 0 0 1 1 1 0 0 1 1 1 1 - 0 1 0 1 0 0 0 1 0 2 0 1 1 1 - - - - - - - - - - - - - - - - - - ? ? - - ? ? ? ? ? ? ? ? ? ? ? ? ? ? ? ? ? ? ? ? ? ? ? ? ? ? ? ? ? ? ? ? ? ? ? ? ? ? ? ? 0 0 1 ? ? ? ? 1 8 0 1 1 ? ? ? ? ? ? ? ? ? ? ? ? ? ? ? ? ? ? ? ? ? ? ? ? ? ? ? ? ? ? ? ? ? ? 0 ? ? ? ? ?

Albadraco_tharmisensis ? ? ? ? ? - ? ? ? ? ? 0 0 ? ? ? ? ? ? ? ? ? ? ? ? ? ? ? ? ? ? ? ? ? ? ? ? ? ? ? ? ? ? ? ? ? ? ? ? ? ? ? ? ? ? ? ? ? ? ? ? ? ? ? ? ? ? ? ? ? ? ? ? ? ? ? ? ? ? ? ? ? ? ? ? ? 1 1 2 1 0 0 1 1 1 0 0 1 1 1 1 - 0 1 0 1 0 0 0 1 0 2 0 1 1 1 - - - - - - - - - - - - - - - - - - ? ? - - ? ? ? ? ? ? ? ? ? ? ? ? ? ? ? ? ? ? ? ? ? ? ? ? ? ? ? ? ? ? ? ? ? ? ? ? ? ? ? ? 0 0 ? ? ? ? ? ? ? ? ? ? ? ? ? ? ? ? ? ? ? ? ? ? ? ? ? ? ? ? ? ? ? ? ? ? ? ? ? ? ? ? ? ? ? ? ? ? ? ? ?

Cryodrakon_boreas 2 - 1 0 0 - 1 0 0 0 1 0 0 0 - - 1 - - 1 0 0 1 2 ? ? 1 0 - 1 1 - 1 6 [23] 0 1 1 0 - - - - 1 - - - - ? ? ? 1 0 0 ? ? ? ? 1 2 3 0 1 ? 1 0 1 0 1 0 - 1 ? ? ? ? ? ? 0 0 - - 0 1 1 ? 1 1 1 1 0 0 0 - 1 0 0 1 0 0 1 - 0 0 0 1 0 0 1 1 0 1 - - 1 1 - - - - - - - - - - - - - - - - - - - - - - 1 2 2 1 1 1 0 1 1 1 0 0 1 2 2 1 0 0 1 2 2 1 1 1 0 0 0 ? 1 1 0 0 0 1 0 1 1 1 8 0 1 1 1 1 1 0 0 4 1 2 ? 1 0 0 ? 1 3 ? ? 1 1 3 ? ? 1 1 1 1 1 1 1 1 2 1 0 1 4 - 1 0 0 0 0 0 0 1 0 0

Arambourgiania_philadelphiae ? ? ? ? ? ? ? ? ? ? ? ? ? ? ? ? ? ? ? ? ? ? ? ? ? ? ? ? ? ? ? ? ? ? ? ? ? ? ? ? ? ? ? ? ? ? ? ? ? ? ? ? ? ? ? ? ? ? ? ? ? ? ? ? ? ? ? ? ? ? ? ? ? ? ? ? ? ? ? ? ? ? ? ? ? ? ? ? ? ? ? ? ? ? ? ? ? ? ? ? ? ? ? ? ? ? ? ? ? ? ? ? ? ? ? ? ? ? ? ? ? ? ? ? ? ? ? ? ? ? ? ? ? ? ? ? ? ? ? ? 0 0 1 2 2 1 1 ? ? ? ? ? ? ? ? ? ? ? ? ? ? ? ? ? ? ? ? ? ? ? ? ? ? ? ? ? ? ? ? ? ? ? ? ? ? ? ? ? ? ? ? ? ? ? ? ? ? 1 ? ? ? ? ? ? ? ? ? ? ? ? ? ? ? ? ? ? ? ? ? ? ? ? ? ? ? ? ? ? ? ?

WAM_60_57	? ? ? ? ? ? ? ? ? ? ? ? ? ? ? ? ? ? ? ? ? ? ? ? ? ? ? ? ? ? ? ? ? ? ? ? ? ? ? ? ? ? ? ? ? ? ? ? ? ? ? ? ? ? ? ? ? ? ? ? ? ? ? ? ? ? ? ? ? ? ? ? ? ? ? ? ? ? ? ? ? ? ? ? ? ? ? ? ? ? ? ? ? ? ? ? ? ? ? ? ? ? ? ? ? ? ? ? ? ? ? ? ? ? ? ? ? ? ? ? ? ? ? ? ? ? ? ? ? ? ? ? ? ? ? ? ? ? ? ? ? ? ? ? ? ? ? ? ? ? ? ? ? ? ? ? ? ? ? ? ? ? ? ? ? ? ? ? ? ? ? ? ? ? ? ? ? ? ? ? ? ? ? ? 1 0 ? ? ? ? ? ? ? ? ? ? ? ? ? ? ? ? ? ? ? ? ? ? ? ? ? ? ? ? ? ? ? ? ? ? 1 0 0 0 ? ? 0 ? ? ?

Quetzalcoatlus_spp 2 - 1 0 0 - 1 0 0 0 1 0 0 0 - - 1 - - 1 0 0 1 2 ? ? 1 0 - 1 1 - 1 6 [23] 0 1 1 0 - - - - 1 - - - - ? ? ? 1 0 0 ? ? ? ? 1 2 3 0 1 ? 1 0 1 0 1 0 - 1 ? ? ? ? ? ? 0 0 - - 0 1 1 ? 1 1 1 1 0 0 0 - 1 0 0 1 0 0 1 - 0 0 0 1 0 0 1 1 0 1 - - 1 1 - - - - - - - - - - - - - - - - - - - - - - 0 1 0 0 1 2 2 1 1 1 0 1 1 1 ? ? ? ? ? 1 0 0 1 2 2 1 1 1 0 0 0 ? 1 1 0 0 0 1 0 1 1 1 8 0 1 1 1 1 1 0 0 4 1 2 ? 1 0 0 ? 1 3 ? ? 1 1 3 ? ? 1 1 1 1 1 1 1 1 2 1 0 1 4 - 1 0 0 0 0 0 0 1 0 0

;

nstates stand;

ccode + 51 53 63 64 79 83 85 97 101 110 111 119 139 140 158 160 167 174 175 177 180 183 184 185 194 206 213 242 250 265 269 270;

ccode - 52 54 55 56 57 58 59 60 61 62 65 66 67 68 69 70 71 72 73 74 75 76 77 78 80 81 82 84 86 87 88 89 90 91 92 93 94 95 96 98 99 100 102 103 104 105 106 107 108 109 112 113 114 115 116 117 118 120 121 122 123 124 125 126 127 128 129 130 131 132 133 134 135 136 137 138 141 142 143 144 145 146 147 148 149 150 151 152 153 154 155 156 157 159 161 162 163 164 165 166 168 169 170 171 172 173 176 178 179 181 182 186 187 188 189 190 191 192 193 195 196 197 198 199 200 201 202 203 204 205 207 208 209 210 211 212 214 215 216 217 218 219 220 221 222 223 224 225 226 227 228 229 230 231 232 233 234 235 236 237 238 239 240 241 243 244 245 246 247 248 249 251 252 253 254 255 256 257 258 259 260 261 262 263 264 266 267 268;

cnames

{0 Skull_aspect_ratio,_length_relative_to_height_at_most_posterior_point_preserved_between_anterior_margin_of_external_naris_or_nasoantorbital_fenestra_and_jaw_articulation_exclusive_of_cranial_crests:_continuous;

{1 Skull,_length_to_squamosal_relative_to_dorsal_vertebra_length:_continuous;

{2 Mandble,_length_relative_to_skull_length_to_squamosal:_continuous;

{3 Rostrum,_length_to_external_naris_(or_nasoantorbital_fenestra)_relative_to_skull_length_to_squamosal:_continuous;

{4 External_naris,_length_relative_to_skull_length_to_squamosal:_continuous;

{5 External_naris,_length_relative_to_height:_continuous;

{6 Antorbital_fenestra,_length_relative_to_skull_length_to_squamosal:_continuous;

{7 Antorbital_fenestra,_length_relative_to_height:_continuous;

{8 Nasoantorbital_fenestra,_length_relative_to_skull_length_to_squamosal:_continuous;

{9 Nasoantorbital_fenestra,_length_relative_to_height:_continuous;

{10 Orbit,_length_relative_to_height:_continuous;

{11 Supratemporal_fenestra,_length_relative_to_skull_length_to_squamosal:_continuous;

{12 Subtemporal_fenestra,_length_relative_to_width:_continuous;

{13 Basipterygoid_processes,_angle_divided_by_100:_continuous;

{14 Rostral_tooth_row,_length_to_posterior_margin_relative_to_skull_length_to_squamosal:_continuous;

{15 Teeth,_maximum_number_divided_by_1000:_continuous;

{16 Mandibular_symphysis,_length_relative_to_mandible_length:_continuous;

{17 Mandible_length,_relative_to_ramus_mid-depth:_continuous;

{18 Mandibular_crest,_length_relative_to_mandible_length:_continuous;

{19 Mandibular_tooth_row,_length_relative_mandible_length:_continuous;

{20 Mid-cervical_vertebra,_maximum_length_relative_to_mid-width:_continuous;

{21 Mid-cervical_vertebra,_maximum_length_relative_to_dorsal_vertebra_length:_continuous;

{22 Dorsal_vertebra,_length_relative_to_maximum_diameter:_continuous;

{23 Synsacral_verebra,_number:_continuous;

{24 Caudal_vertebra,_length_relative_to_dorsal_vertebra_length:_continuous;

{25 Caudal_vertebra,_length_relative_to_diameter:_continuous;

{26 Scapula,_length_relative_to_coracoid_length:_continuous;

{27 Deep_coracoid_flange,_length_relative_to_coracoid_length:_continuous;

{28 Humerus,_length_relative_to_dorsal_vertebra_length:_continuous;

{29 Humerus,_deltopectoral_crest,_proximodistal_constriction_width_relative_to_anterior_terminus_proximodistal_width:_continuous;

{30 Ulna_or_radius,_length_relative_to_humerus_length:_continuous;

{31 Radius,_diameter_relative_to_ulna_diameter:_continuous;

{32 Pteroid,_length_relative_to_ulna_length:_continuous;

{33 Metacarpal_IV,_length_relative_to_humerus_length:_continuous;

{34 Metacarpal_IV_mid-width_relative_to_combined_ulna_and_radius_mid-width:_continuous;

{35 Metacarpal_IV_proximal_end,_dorsoventral_width_relative_to_mid-width:_continuous;

{36 Manual_digit_IV_first_phalanx,_length_relative_to_humerus_length:_continuous;

{37 Manual_digit_IV_second_phalanx,_length_relative_to_first_phalanx_length:_continuous;

{38 Manual_digit_IV_third_wing_phalanx,_length_relative_to_first_phalanx_length:_continuous;

{39 Manual_digit_IV_fourth_wing_phalanx,_length_relative_to_first_phalanx_length:_continuous;

{40 Prepubis,_length_relative_to_width:_continuous;

{41 Pubis,_dorsoventral_depth_relative_to_acetabulum_anteroposterior_length:_continuous;

{42 Ilium_anterior_process,_length_relative_to_posterior_process_length:_continuous;

{43 Femur,_length_relative_to_humerus_length:_continuous;

{44 Tibia,_length_relative_to_femur_length:_continuous;

{45 Fibula,_free_length_relative_to_tibia_length:_continuous;

{46 Metatarsal_III,_length_relative_to_tibia_length:_continuous;

{47 Pedal_digit_III_second_phalanx_length,_relative_to_mid-width:_continuous;

{48 Pedal_digit_IV_first_phalanx,_length_relative_to_mid-width:_continuous;

{49 Pedal_digit_IV_second_phalanx,_length_relative_to_mid-width:_continuous;

{50 Pedal_digit_IV_third_phalanx,_length_relative_to_mid-width:_continuous;

{51 Rostrum_anterior_margin,_shape: flat_surface blunt sharp_tip rostral_process _ _ _ _ _ _ ;

{52 Rostral_process_cross-section,_shape: triangular elliptical _ _ _ _ _ _ _ _ ;

{53 Rostrum_anterior_end,_orientation:_ordered upturned straight downturned _ _ _ _ _ _ _ ;

{54 Palate_anterior_end,_shape: absent present _ _ _ _ _ _ _ _ ;

{55 Rostrum,_anterior_end,_lateral_expansion: absent present _ _ _ _ _ _ _ _ ;

{56 Jaws,_anterior_expansion,_horizontal_outline_shape: elliptical triangular quadrangular _ _ _ _ _ _ _ ;

{57 Rostrum,_anterior_occlusal_margins,_shape: rounded_edges sharp_or_ridged _ _ _ _ _ _ _ _ ;

{58 Rostrum,_middle_expansion: absent present _ _ _ _ _ _ _ _ ;

{59 Rostrum,_posterior_occlusal_margins,_shape: rounded sharp_or_ridged _ _ _ _ _ _ _ _ ;

{60 Rostrum,_shape: laterally_attenuated anteroposteriorly_shortened dorsoventrally_depressed laterally_flattened _ _ _ _ _ _ ;

{61 Rostrum,_dorsal_taper: subparallel attenuated _ _ _ _ _ _ _ _ ;

{62 Jaws,_anterior_end,_lateral_taper: attenuated subparallel _ _ _ _ _ _ _ _ ;

{63 Skull,_entire_margin,_lateral_shape: concave straight_attenuated convex _ _ _ _ _ _ _ ;

{64 Skull_dorsal_margin,_curvature_exclusive_of_cranial_crests:_ordered convex straight concave _ _ _ _ _ _ _ ;

{65 External_naris_dorsal_and_ventral_margins,_orientation: acute_angle subparallel _ _ _ _ _ _ _ _ ;

{66 External_naris_(or_nasoantorbital_fenestra)_anterior_margin,_position_relative_to_premaxillary_toothrow: dorsal posterior _ _ _ _ _ _ _ _ ;

{67 Antorbital_(or_nasoantorbital)_fossa_on_jugal: present absent _ _ _ _ _ _ _ _ ;

{68 Antorbital_fenestra_dorsal_and_ventral_margins,_orientation: subparallel acute_angle _ _ _ _ _ _ _ _ ;

{69 Antorbital_fenestra_ventral_margin,_position_relative_to_external_naris_ventral_margin: same_level ventral _ _ _ _ _ _ _ _ ;

{70 External_naris_and_antorbital_fenestra,_configuration: separate confluent _ _ _ _ _ _ _ _ ;

{71 Antorbital_(or_nasoantorbital_fenestra)_posterior_margin,_shape: subangular beveled _ _ _ _ _ _ _ _ ;

{72 Nasoantorbital_fenestra_dorsal_and_ventral_margins,_orientation: acute_angle subparallel _ _ _ _ _ _ _ _ ;

{73 Orbit_outline,_shape: subcircular piriform inverted_triangle _ _ _ _ _ _ _ ;

{74 Orbit,_dorsal_position: middle_of_the_skull_with_the_ventral_margin_of_the_orbit_below_the_middle_of_the_antorbital_(or_nasoantorbital)_fenestra_and_the_dorsal_margin_of_the_orbit_above_the_dorsal_margin_of_the_antorbital_(or_nasoantorbital)_fenestra high_in_the_skull_with_the_ventral_margin_of_the_orbit_the_same_level_or_above_the_middle_of_the_antorbital_(or_nasoantorbital)_fenestra low_in_the_skull_with_the_entire_orbit_lower_than_the_dorsal_margin_of_the_antorbital_(or_nasoantorbital)_fenestra _ _ _ _ _ _ _ ;

{75 Infratemporal_fenestra,_shape: trapezoidal inverted_triangle upright_triangle oval elliptical _ _ _ _ _ ;

{76 Infratemporal_fenestra,_position_relative_to_orbit: posterior_to_orbit reaches_under_orbit _ _ _ _ _ _ _ _ ;

{77 Infratemporal_fenestra,_orientation: subvertical inclined _ _ _ _ _ _ _ _ ;

{78 Premaxillary_bar_(internasal_process),_width: wide narrow _ _ _ _ _ _ _ _ ;

{79 Premaxilla,_maxillary_process,_position:_ordered contacts_nasal reaches_posterior_half_of_external_naris anterior_to_middle_of_external_naris _ _ _ _ _ _ _ ;

{80 Premaxilla,_posterior_process,_posterior_margin_position: terminate_between_nasals contacts_frontals _ _ _ _ _ _ _ _ ;

{81 Premaxillary_crest: absent present _ _ _ _ _ _ _ _ ;

{82 Premaxillary_crest_anterior_margin,_position_relative_to_skull_anterior_margin: level posterior _ _ _ _ _ _ _ _ ;

{83 Premaxillary_crest_anterior_margin,_orientation:_ordered inclined_posteriorly subvertical curving_anterodorsally _ _ _ _ _ _ _ ;

{84 Premaxillary_crest,_shape: tall_triangle_decreasing_in_height_posteriorly low_blade low_with_anterior_humped_margin comb-like_with_straight_dorsal_margin semicircular tall_triangle_increasing_in_height_posteriorly rectangular _ _ _ ;

{85 Premaxillary_crest_posterior_margin,_position:_ordered anterior_to_external_naris_(or_nasoantorbital_fenestra)_anterior_margin between_external_naris_(or_nasoantorbital_fenestra)_anterior_margin_and_orbit above_orbit above_occipital_region _ _ _ _ _ _ ;

{86 Premaxillary_crest_dorsal_spine: absent present _ _ _ _ _ _ _ _ ;

{87 Premaxillary_crest,_thickness: single_plate two_plates_separated_by_trabeculae _ _ _ _ _ _ _ _ ;

{88 Premaxillary_crest,_texture: striated smooth branching_system_of_grooves _ _ _ _ _ _ _ ;

{89 Maxilla_posterior_end,_shape: narrow ventrally_expanded _ _ _ _ _ _ _ _ ;

{90 Maxilla_ascending_process,_shape: broad tapered slender _ _ _ _ _ _ _ ;

{91 Antorbital_fossa_on_maxilla: present absent _ _ _ _ _ _ _ _ ;

{92 Maxilla_and_nasal_contact,_position: maxilla_contacts_main_body_of_nasal maxilla_contacts_only_descending_process_of_nasal _ _ _ _ _ _ _ _ ;

{93 Maxilla,_premaxillary_and_jugal_processes,_shape: jugal_process_wider both_narrow premaxillary_process_wider both_wide _ _ _ _ _ _ ;

{94 Nasal_descending_process: present absent _ _ _ _ _ _ _ _ ;

{95 Nasal_descending_process,_position: lateral medial _ _ _ _ _ _ _ _ ;

{96 Nasal_descending_process,_length: short elongate _ _ _ _ _ _ _ _ ;

{97 Nasal_descending_process,_orientation:_ordered inclined_anteriorly ventral inclined_posteriorly _ _ _ _ _ _ _ ;

{98 Nasal_process,_lateral_pneumatic_foramen: absent present _ _ _ _ _ _ _ _ ;

{99 Frontal_crest: absent present _ _ _ _ _ _ _ _ ;

{100 Frontal_crest,_shape: low_and_blunt low_and_elongated high_and_expanded _ _ _ _ _ _ _ ;

{101 Frontal_crest_anterior_margin,_position: anterior_to_orbit above_orbit posterior_to_orbit _ _ _ _ _ _ _ ;

{102 Frontal_anterior_margin,_position_relative_to_preorbital_bar_anterior_margin: anterior posterior _ _ _ _ _ _ _ _ ;

{103 Lacrimal_foramen: absemt present _ _ _ _ _ _ _ _ ;

{104 Lacrimal_descending_process_posterior_margin,_shape: flat orbital_process _ _ _ _ _ _ _ _ ;

{105 Parietal_crest: absent present _ _ _ _ _ _ _ _ ;

{106 Parietal_crest,_shape: blunt expanded_into_rounded_margin tapered_into_triangular_process elongate_process _ _ _ _ _ _ ;

{107 Squamosal,_shape: unexpanded rounded expanded _ _ _ _ _ _ _ ;

{108 Squamosal,_position: above_base_of_lacrimal_process_of_jugal below_or_level_with_base_of_lacrimal_process_of_jugal _ _ _ _ _ _ _ _ ;

{109 Jugal_posterior_process: present absent _ _ _ _ _ _ _ _ ;

{110 Quadrate,_inclination_relative_to_ventral_margin_of_skull:_ordered anteriorly subvertical 120˚_posteriorly 150˚_posteriorly _ _ _ _ _ _ ;

{111 Mandibular_articulation,_position_relative_to_center_of_orbit:_ordered posterior_to_orbit posterior_to_center_below_orbit underneath_center_below_orbit anterior_to_center_below_orbit anterior_to_orbit _ _ _ _ _ ;

{112 Quadrate,_shape: wide thin_and_cylindrical _ _ _ _ _ _ _ _ ;

{113 Jugal_ventral_margin,_shape: straight concave _ _ _ _ _ _ _ _ ;

{114 Jugal_anterior_margin,_position_relative_to_nasoantorbital_fenestra_anterior_margin: posterior same_level_or_anterior _ _ _ _ _ _ _ _ ;

{115 Jugal_maxillary_process: absent present _ _ _ _ _ _ _ _ ;

{116 Jugal_postorbital_process_and_lacrimal,_configuration: do_not_contact contact_to_form_lower_orbital_bar _ _ _ _ _ _ _ _ ;

{117 Jugal_ascending_and_postorbital_processes,_shape: separated_by_distinct_angle infilled_by_concave_flange _ _ _ _ _ _ _ _ ;

{118 Jugal_ascending_process_base,_width: broad narrow _ _ _ _ _ _ _ _ ;

{119 Jugal_ascending_process,_inclination:_ordered anterodorsal vertical posterodorsal _ _ _ _ _ _ _ ;

{120 Jugal_postorbital_process_anterior_margin,_shape: flat orbital_process _ _ _ _ _ _ _ _ ;

{121 Jugal_posterior_process,_orientation: posterior ventral _ _ _ _ _ _ _ _ ;

{122 Jugal_maxillary_process: absent present _ _ _ _ _ _ _ _ ;

{123 Occiput,_orientation:_ordered posterior posteroventral ventral _ _ _ _ _ _ _ ;

{124 Basioccipital,_length_relative_to_width: shorter_than_wide longer_than_wide _ _ _ _ _ _ _ _ ;

{125 Basisphenoid_body,_length: shorter_than_wide at_least_longer_than_wide _ _ _ _ _ _ _ _ ;

{126 Elongate_basipterygoid_processes: absent present _ _ _ _ _ _ _ _ ;

{127 Supraoccipital_crest: absent present _ _ _ _ _ _ _ _ ;

{128 Supraoccipital,_pneumatic_foraminae: absent present _ _ _ _ _ _ _ _ ;

{129 Palate,_posterior_end,_shape: concave convex _ _ _ _ _ _ _ _ ;

{130 Palatal_ridge: absent present _ _ _ _ _ _ _ _ ;

{131 Palatal_ridge,_position: tapering_anteriorly confined_posteriorly _ _ _ _ _ _ _ _ ;

{132 Palatal_ridge_shape: narrow_strip strong_keel _ _ _ _ _ _ _ _ ;

{133 Palatines,_shape: broad_and_flat thin_bars _ _ _ _ _ _ _ _ ;

{134 Internal_nares_and_maxilla,_configuration: contact do_not_contact _ _ _ _ _ _ _ _ ;

{135 Pterygoids,_ventral_position_relative_to_jaw_margin: level_or_dorsal ventral _ _ _ _ _ _ _ _ ;

{136 Interpterygoid_opening,_length_relative_to_subtemporal_fenestra_length: at_least_subtemporal_fenestra shorter_than_subtemporal_fenestra _ _ _ _ _ _ _ _ ;

{137 Mandible_articulation_condyles,_orientation: parasagittal oblique _ _ _ _ _ _ _ _ ;

{138 Foramina_positioned_in_a_row_along_the_lateral_margin_of_the_jaws: present absent _ _ _ _ _ _ _ _ ;

{139 Mandible_anterior_end,_orientation:_ordered upturned straight downturned _ _ _ _ _ _ _ ;

{140 Mandible_anterior_margin,_shape: blunt sharp_tip prow _ _ _ _ _ _ _ ;

{141 Mandible_anterior_end,_shape: compressed_laterally shortened_anteroposteriorly depressed_dorsoventrally _ _ _ _ _ _ _ ;

{142 Mandible,_anterior_expansion: absent present _ _ _ _ _ _ _ _ ;

{143 Mandible_anterior_end_dorsal_jaw_margins,_shape: level eminence _ _ _ _ _ _ _ _ ;

{144 Mandible_anterior_end_distinct_eminence,_height: low high _ _ _ _ _ _ _ _ ;

{145 Mandibular_symphysis,_fusion: unfused fused _ _ _ _ _ _ _ _ ;

{146 Mandibular_symphysis,_orientation: subparallel_to_rami oblique_to_rami _ _ _ _ _ _ _ _ ;

{147 Mandible,_anterior_end_lateral_surfaces,_texture: flat large_foramina pitted _ _ _ _ _ _ _ ;

{148 Mandible,_anterior_occlusal_margins,_shape: rounded_edges sharp_or_ridged _ _ _ _ _ _ _ _ ;

{149 Mandible,_distinct_middle_expansion: absent present _ _ _ _ _ _ _ _ ;

{150 Mandible,_posterior_occlusal_margins,_shape: rounded sharp_or_ridged _ _ _ _ _ _ _ _ ;

{151 Mandibular_rami_distinct_dorsal_eminence: present absent _ _ _ _ _ _ _ _ ;

{152 Mandibular_rami_dorsal_eminence,_shape: rounded pointed _ _ _ _ _ _ _ _ ;

{153 Mandibular_sulcus: absent present _ _ _ _ _ _ _ _ ;

{154 Mandible_symphysis_occlusal_surface_anterior_end,_shape: flat_or_concave fossa keel _ _ _ _ _ _ _ ;

{155 Mandible,_dorsal_surface,_parasagittal_ridges: absent present _ _ _ _ _ _ _ _ ;

{156 Mandible,_symphyseal_cavity: absent present _ _ _ _ _ _ _ _ ;

{157 Mandibular_symphyseal_cavity,_dorsal_shelf,_posterior_margin_relative_to_ventral_symphysis_posterior_margin: dorsal_shelf_posterior_margin_extends_posterior_to_ventral_symphysis_posterior_margin ventral_shelf_posterior_margin_extends_at_least_posterior_to_dorsal_symphysis_posterior_margin _ _ _ _ _ _ _ _ ;

{158 Mandibular_rami_dorsal_margin,_shape:_ordered convex straight concave _ _ _ _ _ _ _ ;

{159 Mandibular_rami,_orientation: straight_to_upturned downcurved _ _ _ _ _ _ _ _ ;

{160 Retroarticular_process,_orientation_relative_to_mandible:_ordered posteroventral subhorizontal posterodorsal _ _ _ _ _ _ _ ;

{161 Retroarticular_process,_shape: triangular subcircular elongate blunt _ _ _ _ _ _ ;

{162 Mandible_ventral_margin,_shape: flat keel crest _ _ _ _ _ _ _ ;

{163 Mandibular_crest,_shape: blade_like_and_low massive_and_deep _ _ _ _ _ _ _ _ ;

{164 Mandibular_crest_anterior_margin,_position: posterior_to_mandible_anterior_margin mandible_anterior_margin _ _ _ _ _ _ _ _ ;

{165 Dentary,_length: do_not_separate_angular_and_surangular separate_angular_and_surangular _ _ _ _ _ _ _ _ ;

{166 Teeth: present absent _ _ _ _ _ _ _ _ ;

{167 Teeth,_spacing_along_jaws:_ordered mesial_teeth_spaced_wider_apart even_along_the_jaws distal_teeth_spaced_wider_apart _ _ _ _ _ _ _ ;

{168 Teeth,_variation_in_shape_along_tooth_row: isodont heterodont _ _ _ _ _ _ _ _ ;

{169 Mesial_heterodont_teeth,_shape: recurved_triangle slender_needle recurved_spike _ _ _ _ _ _ _ ;

{170 Teeth,_shape: recurved_triangle bulbous_triangle slender_needle recurved_cone labiolingually_compressed_triangle recurved_spike _ _ _ _ ;

{171 Teeth,_texture: smooth striated sharp_mesial_and_distal_keels medial_carinae _ _ _ _ _ _ ;

{172 Teeth,_maximum_crown_height_relative_to_basal_width: less_than_four_times_width at_least_four_times_width _ _ _ _ _ _ _ _ ;

{173 Teeth,_lateral_orientation: vertical inclined_laterally _ _ _ _ _ _ _ _ ;

{174 Mesial_teeth,_average_spacing_between_successive_teeth: nearly_touching at_most_diameter_of_teeth more_than_diameter_of_teeth _ _ _ _ _ _ _ ;

{175 Cheek_teeth,_average_spacing_between_successive_teeth:_ordered nearly_touching at_most_diameter_of_teeth more_than_diameter_of_teeth _ _ _ _ _ _ _ ;

{176 Teeth,_size_variation: transition_along_tooth_row disparity_in_size_between_mesial_and_distal_teeth _ _ _ _ _ _ _ _ ;

{177 Upper_dentition,_size_relative_to_lower_dentition:_ordered upper_dentition_significantly_larger subequal upper_dentition_significantly_smaller _ _ _ _ _ _ _ ;

{178 Teeth,_maximum_curvature: displacement_of_curvature_less_than_tooth_diameter displacement_of_curvature_at_least_tooth_diameter _ _ _ _ _ _ _ _ ;

{179 Teeth,_curvature_orientation: posterior lingual _ _ _ _ _ _ _ _ ;

{180 Teeth,_inclination:_ordered upright mesial_teeth_procumbent procumbent _ _ _ _ _ _ _ ;

{181 Cheek_alveoli,_shape: set_in_grooves low undulating_occlusal_margins raised_rims inflated _ _ _ _ _ ;

{182 Cheek_teeth,_denticles: present absent _ _ _ _ _ _ _ _ ;

{183 Teeth_largest_denticles,_shape:_ordered serrations cuspules crenulations low_cusps tall_cusps _ _ _ _ _ ;

{184 Teeth,_maximum_denticle_number:_ordered more_than_50 between_six_and_49 five _ _ _ _ _ _ _ ;

{185 Rostral_tooth_row,_anterior_end,_position: posterior_to_tip_of_rostrum tip_of_rostrum anterior_surface_of_rostrum _ _ _ _ _ _ _ ;

{186 Maxillary_teeth,_position_of_largest_teeth: mesial middle distal _ _ _ _ _ _ _ ;

{187 Fifth_and_sixth_teeth,_distinctly_smaller_than_fourth_and_seventh_and_subequal_in_size: absent present _ _ _ _ _ _ _ _ ;

{188 Mandibular_tooth_row,_anterior_end,_position: tip_of_mandible posterior_to_tip_of_mandible _ _ _ _ _ _ _ _ ;

{189 Occlusal_margin,_orientation_with_respect_to_jaw_margin: parallel dorsally_reflected _ _ _ _ _ _ _ _ ;

{190 Atlantoaxis,_fusion: unfused fused _ _ _ _ _ _ _ _ ;

{191 Mid-cervical_vertebra_neural_arch_lateral_surface,_pneumatic_foramen: absent present _ _ _ _ _ _ _ _ ;

{192 Mid-cervical_vertebra_centrum_lateral_surface,_pneumatic_foramen: absent present _ _ _ _ _ _ _ _ ;

{193 Cervical_vertebra_lateral_to_neural_canal,_pneumatic_foramina: absent present _ _ _ _ _ _ _ _ ;

{194 Mid-cervical_vertebra_neural_spines,_height:_ordered tall low extremely_reduced _ _ _ _ _ _ _ ;

{195 Mid-cervical_vertebrae_neural_spines,_lateral_outline_shape: blade triangular ridge fan _ _ _ _ _ _ ;

{196 Mid-cervical_vertebra,_postexapophyses: absent present _ _ _ _ _ _ _ _ ;

{197 Mid-cervical_vertebra_neural_arch_and_centrum,_configuration: distinct confluent _ _ _ _ _ _ _ _ ;

{198 Mid-cervical_vertebra_ribs,_shape: elongate reduced _ _ _ _ _ _ _ _ ;

{199 Cervical_8_neural_spine,_height: tall low _ _ _ _ _ _ _ _ ;

{200 Cervical_9,_shape: similar_to_dorsal_vertebrae similar_to_cervicals _ _ _ _ _ _ _ _ ;

{201 Notarium: absent present _ _ _ _ _ _ _ _ ;

{202 Anterior_dorsal_vertebra_neural_spines,_shape: unfused supraneural_plate _ _ _ _ _ _ _ _ ;

{203 Sacral_ribs,_configuration: contact_at_ilium contact_medial_to_ilium _ _ _ _ _ _ _ _ ;

{204 Synsacral_supraneural_plate: absent present _ _ _ _ _ _ _ _ ;

{205 Caudal_vertebra,_number: more_than_15 at_most_15 _ _ _ _ _ _ _ _ ;

{206 Caudal_vertebra_zygapophyses,_length: short elongate extremely_elongate _ _ _ _ _ _ _ ;

{207 Caudal_vertebra_centrum,_shape: single duplex _ _ _ _ _ _ _ _ ;

{208 Scapulocoracoid,_orientation_relative_to_vertebral_column: subparallel rotated_laterally _ _ _ _ _ _ _ _ ;

{209 Scapula_proximal_end,_shape: elongate_andcompressed suboval_and_expanded _ _ _ _ _ _ _ _ ;

{210 Scapula,_shape: elongate_process stout_with_constricted_shaft _ _ _ _ _ _ _ _ ;

{211 Scapula_articulates_with_vertebral_column: absent present _ _ _ _ _ _ _ _ ;

{212 Coracoid_ventral_margin,_shape: rounded broad_tubercle deep_flange _ _ _ _ _ _ _ ;

{213 Coracoid,_shape:_ordered semicircular broad_shaft narrow_shaft _ _ _ _ _ _ _ ;

{214 Sternocoracoid_articulations,_position_with_respect_to_one_another: lateral anterior_and_posterior _ _ _ _ _ _ _ _ ;

{215 Sternum,_constriction_posterior_to_sternocoracoid_articulations: present absent _ _ _ _ _ _ _ _ ;

{216 Cristopine,_shape: shallow deep _ _ _ _ _ _ _ _ ;

{217 Cristospine,_length: stout elongate _ _ _ _ _ _ _ _ ;

{218 Sternocoracoid_articulations,_shape: flattened oval _ _ _ _ _ _ _ _ ;

{219 Sternocoracoid_articulations,_posterior_expansion: absent present _ _ _ _ _ _ _ _ ;

{220 Sternum_plate,_shape: narrow quadrangular semicircular triangular laterally_expanded _ _ _ _ _ ;

{221 Humerus_proximal_end_ventral_surface,_pneumatic_foramen: absent present _ _ _ _ _ _ _ _ ;

{222 Humerus_proximal_end,_cross_section: crescent horseshoe _ _ _ _ _ _ _ _ ;

{223 Humerus_proximal_end_dorsal_surface,_pneumatic_foramen: absent present _ _ _ _ _ _ _ _ ;

{224 Humerus_shaft,_shape: straight bowed _ _ _ _ _ _ _ _ ;

{225 Humerus,_mid-shaft,_shape: tapered subcylindrical _ _ _ _ _ _ _ _ ;

{226 Humerus_entepicondyle,_anteroposterior_width_relative_to_ectepicondyle_anteroposterior_width: entepicondyle_wider_than_ectepicondyle ectepicondyle_at_most_entepicondyle_width _ _ _ _ _ _ _ _ ;

{227 Humerus,_between_distal_condyles,_pneumatic_foramen: absent present _ _ _ _ _ _ _ _ ;

{228 Humerus_distal_aspect,_pneumatic_foramen: absent present _ _ _ _ _ _ _ _ ;

{229 Humerus_distal_aspect,_shape: hourglass crescentic_or_D-shape triangular trapezoidal _ _ _ _ _ _ ;

{230 Deltopectoral_crest,_position_on_humerus: proximal more_distally_on_shaft _ _ _ _ _ _ _ _ ;

{231 Humerus_deltopectoral_crest,_shape: subtriangular_with_proximal_apex proximally_leaning_trapezoid proximally_curving_hook oblong_process_with_constricted_neck low_and_rectangular elongate_and_proximally_expanded hatchet-shape distally_leaning_trapezoid tall_rectangular_process _ ;

{232 Humerus_deltopectoral_crest,_curvature: parallel_to_shaft warped_distally _ _ _ _ _ _ _ _ ;

{233 Ulnar_crest_of_humerus,_size: reduced developed _ _ _ _ _ _ _ _ ;

{234 Humerus_ulnar_crest,_orientation: posterior ventral _ _ _ _ _ _ _ _ ;

{235 Ulna_shaft_proximal_anterior_surface,_shape: flat longitudinal_ridge _ _ _ _ _ _ _ _ ;

{236 Ulna_distal_tubercle,_position: middle_of_the_distal_end ventral_part_of_the_distal_end _ _ _ _ _ _ _ _ ;

{237 Radius_distal_end_cross-section,_shape: suboval subtriangular_with_large_anterior_process _ _ _ _ _ _ _ _ ;

{238 Distal_syncarpal_ventral_articular_facet_for_Metacarpal_IV,_size_relative_to_dorsal_facet: ventral_facet_larger subequal_in_size _ _ _ _ _ _ _ _ ;

{239 Distal_syncarpal,_cross-section_shape: rectangular triangular _ _ _ _ _ _ _ _ ;

{240 Pteroid,_shape: angled_at_midsection stout_hook straight_and_tapered_with_expanded_proximal_end straight_with_expanded_ends curved_slender_rod curved_and_subparallel-sided _ _ _ _ ;

{241 Preaxial_carpal,_shape: longer_than_wide at_most_long_as_wide _ _ _ _ _ _ _ _ ;

{242 Metacarpals,_number_articulating_with_carpus:_ordered four_or_more two one _ _ _ _ _ _ _ ;

{243 Metacarpals_I_to_III_distal_ends,_positions: disparate approximate _ _ _ _ _ _ _ _ ;

{244 Metacarpal_IV_proximal_cross-section,_shape: anteroposteriorly_compressed subrectangular _ _ _ _ _ _ _ _ ;

{245 Metacarpal_IV_shaft_cross-section,_shape: rounded_rectangle anteroposteriorly_compressed_oval _ _ _ _ _ _ _ _ ;

{246 Metacarpal_distal_end_between_condyles,_shape: flat median_ridge _ _ _ _ _ _ _ _ ;

{247 Manual_unguals,_size_relative_to_pedal_unguals: less_than_twice_the_size_of_pedal_unguals at_least_twice_the_size_of_pedal_unguals _ _ _ _ _ _ _ _ ;

{248 Manual_digit_IV_first_phalanx_proximal_end_ventral_surface,_pneumatic_foramen: absent present _ _ _ _ _ _ _ _ ;

{249 Manual_digit_IV_second_or_third_phalanges_shaft_cross-sections,_shape: round_to_subtriangular concave_posteriorly oval ventral_ridge _ _ _ _ _ _ ;

{250 Pubis_anterior_margin,_shape_in_lateral_view:_ordered convex straight slightly_concave deeply_concave _ _ _ _ _ _ ;

{251 Pubis_and_ischium_contact,_shape: confluent_along_length partially_separated_by_oval_opening _ _ _ _ _ _ _ _ ;

{252 Ischium_ventral_margin,_shape: straight convex _ _ _ _ _ _ _ _ ;

{253 Prepubis_shaft,_constriction: absent present _ _ _ _ _ _ _ _ ;

{254 Prepubis,_shape: elongate_paddle medially_curved_with_short_lateral_process triradiate expanded_fan _ _ _ _ _ _ ;

{255 Ilium_preactetabular_anterior_margin,_shape: rounded pointed _ _ _ _ _ _ _ _ ;

{256 Ilium_preacetabular_process,_orientation: straight dorsiflected _ _ _ _ _ _ _ _ ;

{257 Ilium_postacetabular_process,_orientation: subhorizontal posterodorsal _ _ _ _ _ _ _ _ ;

{258 Ilium_postacetabular_process_shaft,_constriction: absent present _ _ _ _ _ _ _ _ ;

{259 Ilium_postacetabular_process,_terminal_expansion: absent present _ _ _ _ _ _ _ _ ;

{260 Acetabulum,_shape: anteroposteriorly_ovate subcircular _ _ _ _ _ _ _ _ ;

{261 Ilium_postacetabular_process_apex,_shape: flat torus _ _ _ _ _ _ _ _ ;

{262 Femur,_shape: strongly_bowed slight_curvature _ _ _ _ _ _ _ _ ;

{263 Femur_proximal_end,_pneumatic_foramen: absent present _ _ _ _ _ _ _ _ ;

{264 Femoral_neck,_shape: indistinct constricted _ _ _ _ _ _ _ _ ;

{265 Greater_trochanter,_shape:_ordered reduced distinct_process anteriorly-curved_hook _ _ _ _ _ _ _ ;

{266 Femur_distal_end,_epicondyles_size: reduced_and_confluent_with_distal_condyles expanded_into_distinct_distal_flanges _ _ _ _ _ _ _ _ ;

{267 Femoral_head,_angle_relative_to_shaft: at_most_145° greater_than_145° _ _ _ _ _ _ _ _ ;

{268 Metatarsal_IV,_length_relative_to_metatarsals_I_to_III: subequal significantly_shorter _ _ _ _ _ _ _ _ ;

{269 Pedal_digit_V,_number_of_phalanges:_ordered four three two one zero _ _ _ _ _ ;

{270 Pedal_digit_V_ultimate_phalanx,_shape:_ordered straight curved bent_at_midsection nubbin _ _ _ _ _ _ ;

{271 Bulbous_anterior_ulnar_joint _ _ _ _ _ _ ;

{272 Uncinate_anterior_ulnar_joint _ _ _ _ _ _ ;

{273 Rectangular_anterior_ulnar_joint _ _ _ _ _ _ ;

{274 Obdurate_anterior_ulnar_joint _ _ _ _ _ _ ;

{275 Expanded_opisthotic_processes _ _ _ _ _ _ ;

{276 Strongly_curved_femur _ _ _ _ _ _ ;

{277 Non-pneumatic_limb_bones _ _ _ _ _ _ ;

{278 Vestigial_metacarpals_I-III:_ordered lost_contact_with_wrist _ _ _ _ _ _ ;

{279 Hatchet_shaped_deltopectoral_crest _ _ _ _ _ _ ;

{280 Scapulocoracoid_not_articulating_with_spine _ _ _ _ _ _ ;
